# PepForge: Hierarchical HELM-Based Peptide Generation

**DOI:** 10.64898/2026.05.29.728379

**Authors:** Qingxin Wang, Roderich D. Süssmuth

## Abstract

Peptides carrying special connections such as macrocyclizations and various other structural modifications constitute a major class among peptide therapeutics, yet their chemical space remains largely inaccessible to computational generation methods. Here we present PepForge, a deep learning platform for peptide generation that exploits Hierarchical Editing Language for Macromolecules (HELM) notation to access the chemical space of modified peptides, through a Layout-Content-Connection (LCC) cascade decomposing the generation task into block layout, monomer content, and special connection prediction. The LCC cascade is trained on 383,817 HELM peptides covering 425 monomers and nine connection types. Beyond *de novo* generation, the LCC cascade supports masked infilling for targeted scaffold modification and multi-level constrained generation. Both the monomer library and the connection-type set support user-defined extensions for exploring a broader chemical space. The prediction module is decoupled from generation and accepts arbitrary scoring heads for downstream tasks. As a demonstration, we built an antimicrobial potency ensemble predictor trained on 11,026 peptides with minimum inhibitory concentration (MIC) values, alongside the external PeptiVerse predictor. Applied at scale, we generated 4.78 million novel HELM peptides and obtained 799 structurally novel hit antimicrobial peptide (AMP) candidates after potency and safety filtering. All code, pre-trained models, and a web interface for interactive use are publicly available at https://github.com/wqx1999/PepForge.

## 1 Introduction

Peptide therapeutics constitute one of the fastest-growing drug classes, with >80 approved drugs and hundreds in clinical trials[1, 2]. Nearly all clinically approved peptide drugs carry modifications beyond the 20 canonical amino acids, such as non-proteinogenic building blocks, macrocyclizations, and acylations, all of which contribute to pharmacodynamic and pharmacokinetic properties such as target affinity, metabolic stability, and membrane permeability (Fig. 1a). However, the chemical space of such modified peptides remains largely unexplored by computational methods. Recent deep learning has expanded peptide generation coverage, but each existing method still spans only a limited region of the chemical space along two axes: monomer alphabet richness and special-connection diversity (Fig. 1b). Structure-based methods (RFpeptides[3], AfCycDesign[4]) are restricted to the proteinogenic-only corner, while string-based methods (HELM-GPT[5], PepINVENT[6]) accommodate non-proteinogenic monomers but cover only narrow slices of the connection-type space. Consequently, peptides that simultaneously demand both remain inaccessible to current generative models.

**Fig. 1:**
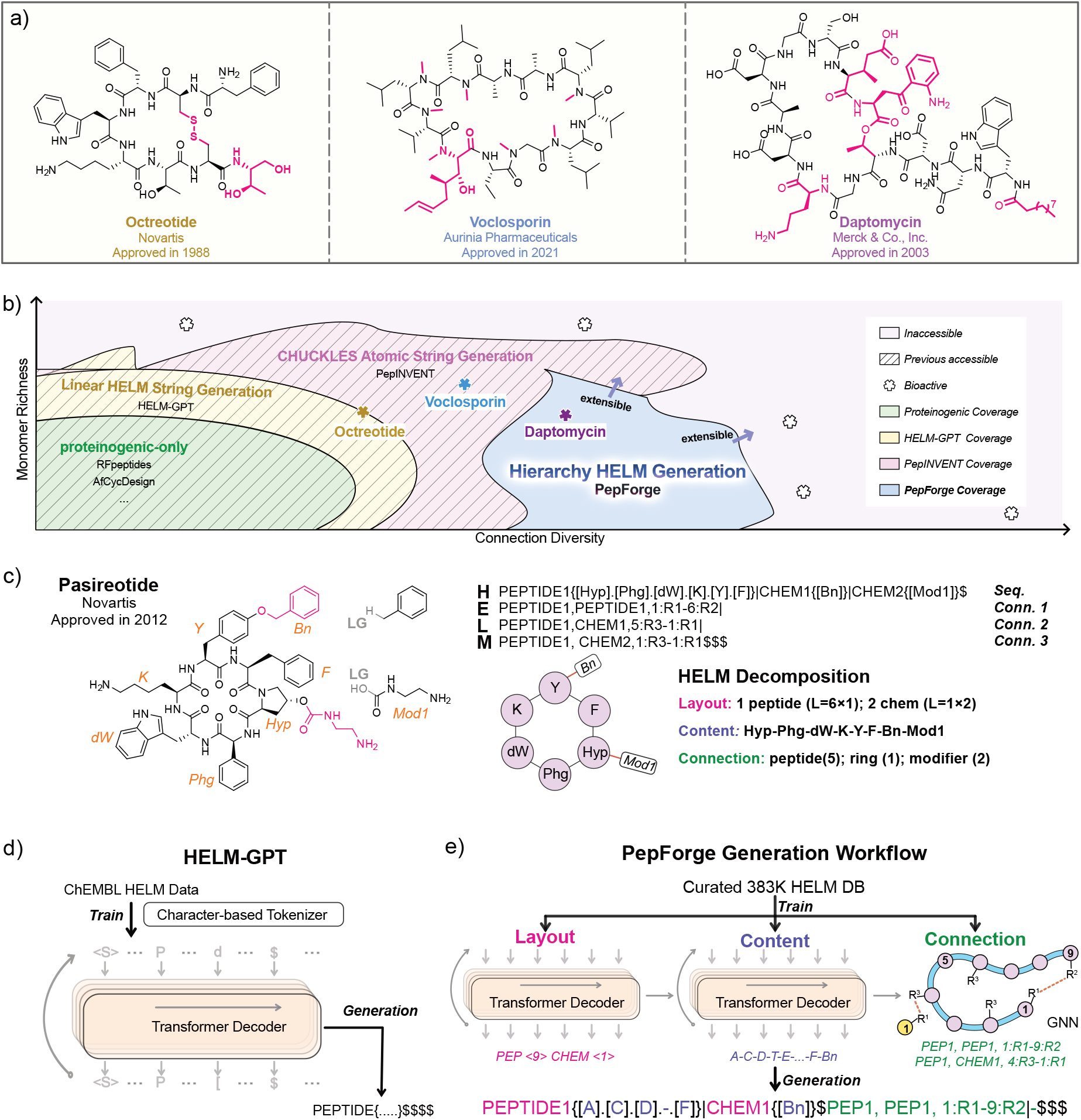
Representative peptide drugs and the HELM-based generation framework. **(a)** FDA-approved peptide drugs octreotide, daptomycin, and voclosporin with special connections, macrocyclizations and/or non-proteinogenic building blocks highlighted in magenta. **(b)** Coverage of four current peptide-generation paradigms along two axes: monomer richness and connection diversity. Star markers locate the three panel-(a) drugs within the region reachable by each paradigm. Open dotted circles mark literature-reported bioactive peptides, many falling in the hatched “Inaccessible” region beyond all current methods. **(c)** HELM decomposition of Pasireotide. HELM representation showing monomer sequences, chemical modifier (`CHEM`) blocks with their leaving-group (LG) annotations, and three special connections; together with the corresponding hierarchical decomposition into *Layout, Content*, and *Connection* levels, rendered as a monomer graph. **(d)** HELM-GPT, a prior end-to-end approach using a Transformer Decoder on flat HELM character sequences. **(e)** PepForge three-stage LCC cascade: *Layout* (block types and lengths) → *Content* (per-block monomer sequences) → *Connection* (special bonds, GNN). The HELM Parser provides bidirectional conversion between HELM strings and the (*Layout, Content, Connection*) three-tuple.

Underlying this gap is a descriptive mismatch: existing peptide notations differ in granularity and in their ability to encode modifications and special connections. Atom-level notations such as SMILES[7] and CHUCKLES[8] describe molecules at fine resolution but do not mark monomer boundaries within a peptide chain. Sequence-level notations such as FASTA encode peptides as one-letter codes, offering no syntax for modifications. Hierarchical notations such as BILN[9] and HELM[10] sit between these extremes by capturing peptides at the block, monomer, and inter-block-connection levels (Fig. 1c), natively accommodating non-proteinogenic monomers, macrocyclization, and chemical modifiers within a single string. HELM-GPT first applied HELM to generative modeling using an end-to-end Transformer decoder trained on ~22,000 HELM sequences from ChEMBL[11] (Fig. 1d), but only 70.8% of generated strings parsed into chemically valid molecules. Fine-tuning on task-specific datasets raised this rate to 100% but narrowed the generative distribution, a trade-off driven by limited data scale.

Rather than treating HELM as a flat token sequence, we exploit its inherent three-level hierarchy: *Layout* (block types and lengths), *Content* (per-block monomer sequences), and *Connection* (special bonds) (Fig. 1c). PepForge implements this insight as a *Layout*–*Content*–*Connection* (LCC) cascade, with each stage conditioned on the previous (Fig. 1e). The current LCC cascade covers nine diverse special connection types (Section 2.2) and supports constrained generation at each structural level. Both the monomer alphabet and the connection-type set are extensible, expanding the chemical space the LCC cascade can currently generate. To build the training database, we developed a bidirectional SMILES-to-HELM (S2H) conversion with roundtrip validation, curating 383,817 HELM peptides with 425 monomers and nine connection types from six public databases. Using the trained LCC cascade, we generated 4.78 million novel HELM peptides covering all nine connection types. To validate the utility of this generation capability, we additionally built a five-class antimicrobial potency predictor on 11,026 peptides with minimum inhibitory concentration (MIC) values as a heterogeneous four-model ensemble, implemented as a swappable plugin module with built-in support for third-party tools such as PeptiVerse[12]. Combining generation with the prediction layer, we apply the pipeline to large-scale antimicrobial candidate screening, yielding 799 structurally novel hit antimicrobial candidates (Section 2.3).

All code, pre-trained models, and a web interface are publicly available at https://github.com/wqx1999/PepForge.

## 2 Results and Discussion

### 2.1 Platform Architecture

Exploiting the three-level hierarchy described above, PepForge implements three integrated components: (1) a bidirectional S2H conversion pipeline that curates large-scale training data from public databases (Fig. 2a), (2) a three-stage LCC cascade architecture with models tailored to each structural level (Fig. 2b), and (3) a decoupled prediction module that can be replaced or extended with external scoring tools (Fig. 2c).

**Fig. 2:**
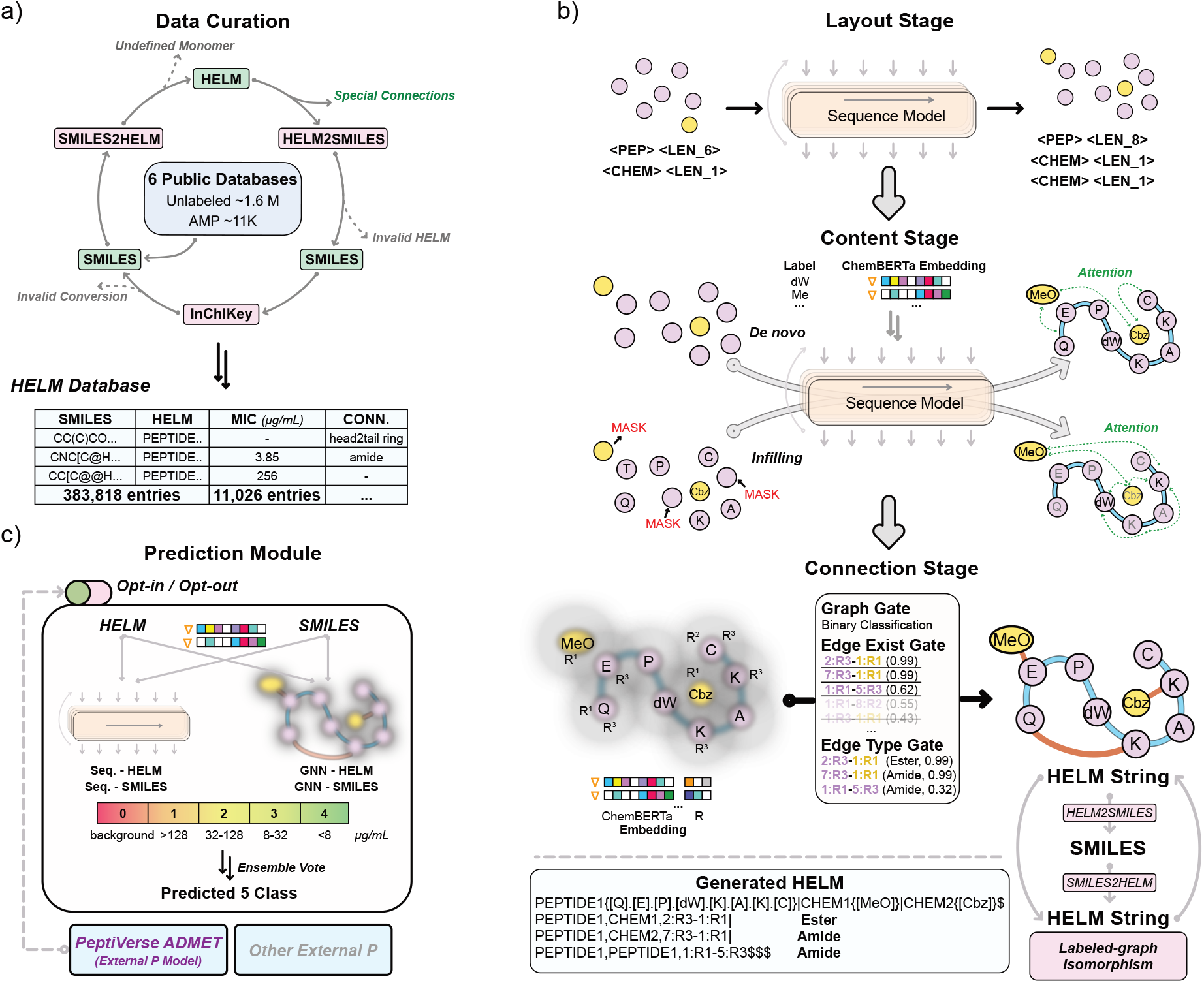
PepForge platform architecture. **(a)** Data curation. Six public databases are aggregated through a bidirectional S2H conversion with InChIKey roundtrip checking; three filters reject entries with undefined monomers, malformed HELM syntax, or InChIKey mismatch after roundtrip. The curated database retains 383,817 unique HELM entries, of which 11,026 carry quantitative antimicrobial biological data (MIC values). **(b)** Three-stage LCC cascade. *Layout* generates the block-level scaffold (block type, length). *Content* fills per-block monomers under *De novo* or Infilling modes. *Connection* predicts special bonds on the assembled monomer graph (R-group positions R^1^/R^2^/R^3^) through three sequential heads (graph gate, edge exist gate, edge type gate). The generated HELM string is validated by a second S2H roundtrip. **(c)** *Prediction* module (opt-in). A four-model heterogeneous ensemble combines two encodings (HELM, SMILES) with two model families (sequence model, GNN architecture), producing five-class antimicrobial predictions (four MIC-defined potency classes plus a background class) via MCC-weighted soft voting. External predictors (e.g., PeptiVerse) plug in as drop-in modules. Trainable ChemBERTa monomer embeddings (denoted ∇) initialize the *Content* and *Connection* stages and the GNN-based predictor.

#### Data curation

We developed a bidirectional S2H conversion pipeline with InChIKey-based roundtrip validation to transform existing molecular databases into HELM[10]-format training data (Fig. 2a, Supplementary Fig. S1). From six public databases, e.g., PubChem[13], ChEMBL[11], and DBAASP[14] (Section 4.1.1), the pipeline yielded 383,817 unique HELM peptides spanning nine special connection types (Supplementary Section S1.2), including 11,026 entries with measured antimicrobial bioactivities (MIC values), hereafter the antimicrobial peptide (AMP) subset (Supplementary Fig. S3). Invalid conversions are filtered at three checkpoints (undefined monomers, malformed HELM, and InChIKey mismatch after roundtrip), ensuring that only chemically faithful representations enter the training set. This curated HELM dataset, together with a 425-monomer library adapted from the Pistoia Alliance[15] (Supplementary Fig. S4), constitutes the shared infrastructure on which all downstream models are built. Pre-computed ChemBERTa[16] monomer embeddings further encode chemical similarity between monomers and are shared across all monomer-level models (Supplementary Figs. S5 and S6).

#### Hierarchical LCC cascade generation module

The LCC cascade generates HELM in three sequential stages, each trained independently (Fig. 2b). The *Layout* and *Content* stages are realized by sequence-model architectures, and the *Connection* stage by a graph neural network (GNN). For each stage, multiple architectures and model sizes were systematically compared (Supplementary Figs. S7–S13). The production LCC cascade configuration and selection criteria are detailed in Section 4.3. The *Layout* stage generates the block-level scaffold by predicting block types and lengths, defining the overall peptide skeleton. The *Content* stage takes this scaffold as input and fills each block with a monomer sequence, conditioned on all known blocks to ensure chemical coherence across the peptide. The *Content* stage supports two decoding modes: autoregressive GPT[17] generation for *de novo* design and BERT[18] masked-language-model infilling for targeted modification, where users fix known positions and the model predicts the rest. The *Connection* stage operates on the assembled monomer graph, where each monomer is a node and candidate edges are restricted to R-group-compatible pairs. Three prediction heads form a hierarchical cascade: (1) a GRAPH GATE determines whether the peptide contains any special connections, (2) an EDGE EXIST GATE identifies which candidate edges correspond to actual bonds, and (3) an EDGE TYPE GATE classifies each positive edge into one of nine connection types. Since each stage consumes the output of the previous one, errors in early stages propagate downstream. Generated peptides are therefore validated via SMILES roundtrip and labeled-graph isomorphism at the final stage (Supplementary Section S1.4), filtering structurally invalid outputs regardless of the failure source. This sequential design also enables users to enter the pipeline at any stage with a partial HELM peptide for layout-constrained, content-constrained, or connection-only generation.

#### Prediction module

To demonstrate a downstream application, we constructed a decoupled prediction module that scores generated peptides without biasing the generation process (Fig. 2c). The built-in default targets antimicrobial potency, formulated as five classes: four MIC-defined potency classes plus a background class (no reported antimicrobial activity) that flags generated peptides that are out-of-distribution (OOD) relative to known active peptides, rather than forcing them onto the potency scale, improving screening precision. The predictor is a four-model heterogeneous ensemble spanning two encoding schemes (HELM and SMILES) and two model families (sequence models and GNN architectures), where HELM encoding preserves hierarchical structural information while SMILES encoding captures atom-level chemical detail. Predictions are aggregated via Matthews correlation coefficient (MCC)-weighted soft voting. Beyond the built-in ensemble, the prediction interface is designed to be extensible: we integrated PeptiVerse[12], a third-party predictor for peptide safety and pharmacokinetic properties, demonstrating that external models can be incorporated without modifying the generation pipeline.

### 2.2 Generation Quality Evaluation

Subsequently, we evaluated the LCC cascade against a Flat GPT baseline, a single autoregressive transformer that we trained from scratch on the same HELM corpus as the LCC cascade, generating complete HELM strings without hierarchical decomposition (metric definitions in Section 4.8.1). We retrained rather than adopt the prior HELM-GPT model[5], whose 70.8% validity reflects both a much smaller training corpus of around 22,000 sequences and a character-level tokenizer that does not treat monomer codes as semantic units (e.g., `[Ala]` is split into the five tokens `[A l a]`). Our comparison thus isolates architecture choice from both data scale and tokenization. Both architectures substantially exceed the HELM-GPT prior model’s validity, confirming that the expanded dataset and improved tokenization, rather than the architecture, drive the gain. We compared the two approaches on core generation metrics and chemical space coverage, performed per-connection-type analysis, and evaluated the *Content* stage’s infilling accuracy with masked inputs.

#### Core metrics and chemical-space coverage

At the 100K generation scale, our LCC cascade achieved lower validity than Flat GPT (88.3% vs 99.6%) but markedly higher uniqueness (92.5% vs 85.5%) and novelty (66.5% vs 40.4%), while the internal diversity remained comparable (0.88–0.89 across both models and the training set) (Fig. 3a). The nearest-neighbor distance under MAP4C[19, 20], a stereochemistry-aware molecular fingerprint, confirmed this divergence. Flat GPT produced a median of 0.293, below the training set’s self-distance of 0.346, while the LCC cascade reached 0.432, exceeding the training distribution and exploring previously uncovered chemical space (Fig. 3b). Together with the low uniqueness and novelty, the compressed distance distribution points to training-set memorisation in the baseline model, heavily reproducing the training distribution. At the building-block level, the LCC cascade covered 385/386 training monomers (99.7%) plus 21 out-of-training monomers, whereas Flat GPT covered only 300/386 (77.7%) plus 1 out-of-training monomer (Supplementary Fig. S14), confirming that the LCC cascade’s broader exploration is grounded in vocabulary breadth rather than recombination of a narrow palette. This divergence became more pronounced for structurally complex peptides (Fig. 3c). As the number of special connections increased, the LCC cascade’s advantage in uniqueness widened progressively, while its novelty advantage remained large across all subgroups and was largest at the 3+ connections subgroup. Flat GPT’s validity remained near-perfect but its uniqueness dropped sharply, and its novelty stayed similar to or below the 1-connection level rather than rising with the broader exploration LCC cascade achieved, consistent with the inherent data imbalance where multi-connection peptides are underrepresented in the training set. However, LCC cascade’s broader exploration comes at a cost: its validity decreased with connection complexity, dropping to 49.3% for peptides with three or more connections.

**Fig. 3:**
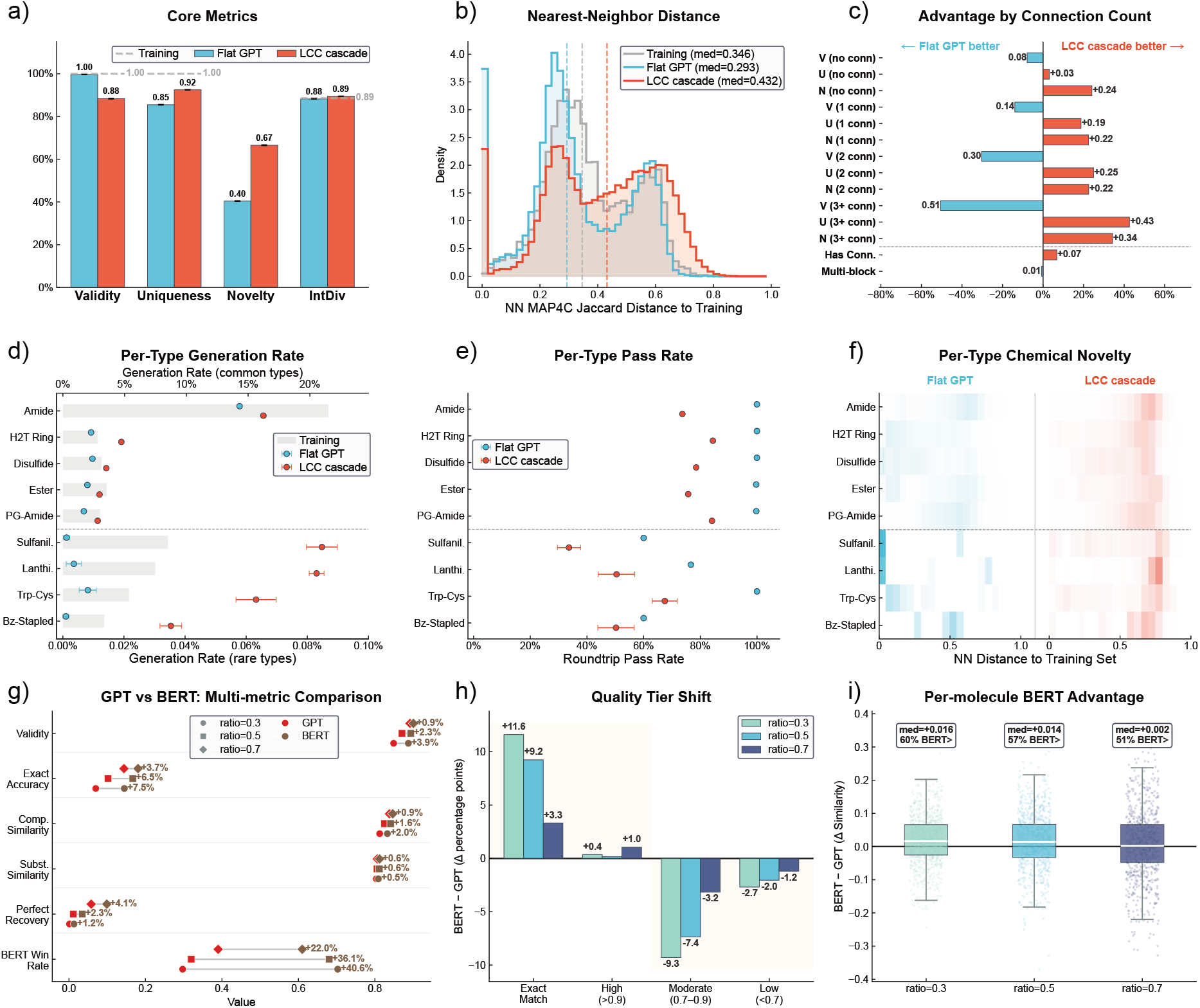
Generation quality evaluation. **(a–f)** LCC cascade vs Flat GPT, 100K peptides per run, 5 independent runs. **(g– i)** Comparison of two production models (BERT, GPT) in *Content*-stage infilling on 10K held-out peptides, evaluated at reveal ratios 0.3/0.5/0.7 (fraction of monomer positions retained as context). Metric definitions in Section 4.8.1 and Section 4.8.2. **(a)** Core generation metrics (**V**alidity, **U**niqueness, **N**ovelty, **Int**ernal **Div**ersity). Gray dashed lines: training-set reference. **(b)** Nearest-neighbor MAP4C Tanimoto distance to the training set. Dashed vertical lines: median of each distribution. **(c)** LCC cascade − Flat advantage (Δ V, Δ U, Δ N) stratified by the number of special connections per peptide. **(d)** Per-connection-type generation rate (fraction of outputs containing each connection type). **(e)** Per-connection-type roundtrip pass rate. Error bars are omitted for rare types where Flat GPT produced only 0–12 peptides per run. **(f)** Per-connection-type NN MAP4C distance heatmap (left: Flat GPT, right: LCC cascade), row-normalized. Near-zero values indicate training-set recall rather than novel generation. **(g)** BERT vs GPT dumbbell chart across six infilling metrics. Points mark each model’s value; bar length encodes the BERT − GPT gap. **(h)** Quality tier shift (BERT − GPT, Δpp), with the four tiers defined in Section 4.8.2. Positive bars indicate a larger share of predictions in that tier for BERT. **(i)** Per-peptide BERT − GPT compositional similarity Δ. Positive values mean BERT produces more faithful monomer compositions at that peptide.

#### Per-connection-type analysis

The LCC cascade and Flat GPT diverged fundamentally across connection types (Fig. 3d–f; connection types defined in Supplementary Table S1). For common connection types (amide, head-to-tail ring, disulfide, ester, PG-amide), both models achieved comparable generation rates, though LCC cascade consistently allocated a slightly larger proportion to each type (Fig. 3d). Four rare connection types separated the two models: sulfanilamide, lanthionin, Trp-Cys, and Bz_stapled (benzyl-thioether staple). LCC cascade maintained meaningful generation rates across all four, whereas Flat GPT produced near-zero counts, indicating that the end-to-end model largely failed to learn these underrepresented connection types. Flat GPT achieved higher pass rates across all connection types (Fig. 3e). However, for rare types, Flat GPT generated only 0–12 peptides per run, making its pass rate fluctuations a product of sampling noise from extremely small counts rather than a measure of model capability. LCC cascade, generating 35–102 rare-type peptides per run, provided more stable pass rate estimates across all types. The per-type nearest-neighbor distance heatmap (Fig. 3f) explains how Flat GPT achieves its high pass rates. Most outputs recapitulate training peptides rather than explore new chemistry, so passing roundtrip validation becomes a near-tautology for recalled structures. Amide and PG-amide showed the smallest Flat-vs-LCC gap, while for all other common types Flat GPT’s distance distributions shifted noticeably toward zero. Rare types amplified the gap: Flat GPT’s sulfanilamide and lanthionin outputs concentrated almost entirely at zero distance, while its Trp-Cys and Bz_stapled outputs either clustered near zero or had too few samples for reliable distribution shapes. LCC cascade, in contrast, produced distributions that spread broadly across all connection types, including the rare ones.

#### *Content*-stage infilling

Beyond *de novo* generation, the *Content* stage also supports constrained generation through masked infilling, where designated monomer positions are hidden and the model predicts replacements conditioned on the surrounding context (metric definitions in Section 4.8.2). Within the *Content* stage, we compared both architectures at three reveal ratios (0.3, 0.5, 0.7), defined as the fraction of monomer positions kept as context (Fig. 3g–i). BERT outperformed GPT on every metric evaluated (Fig. 3g). For validity, exact monomer accuracy, and substitution similarity, BERT’s lead was largest at the lowest reveal ratio, where the task is most context-starved and bidirectional attention pays off most, and narrowed as the reveal ratio increased and GPT’s causal context alone sufficed. Yet the advantage persisted at every ratio. Compositional similarity declined more slowly than position-level metrics, suggesting that BERT retains global compositional awareness even when most positions must be regenerated. At the position level, BERT consistently shifted the quality tier distribution toward higher similarity (Fig. 3h), with most of the shift concentrated in the lowest reveal ratio where bidirectional context matters most. At the peptide level (Fig. 3i), BERT produced higher compositional similarity than GPT across all ratios, with the per-peptide advantage narrowing toward parity at reveal ratio 0.7.

Beyond position-level infilling, the LCC cascade supports multi-level constrained generation: users can fix block layouts, designate monomer positions, or specify forced connections, and these constraints compose freely across stages (Supplementary Table S7). With both *de novo* and constrained generation validated, we next applied the LCC cascade to large-scale AMP screening.

### 2.3 Prediction and Candidate Discovery

Having established that the LCC cascade generates diverse peptides with broad connection-type coverage, we now apply it to a specific downstream task to demonstrate that generation quality translates into useful candidate discovery. The pipeline is task-agnostic by design, with the prediction layer decoupled from generation and swappable without retraining the LCC cascade. As an example, we constructed a built-in AMP potency predictor as the default scorer, while the prediction interface remains open to external tools. PeptiVerse[12], a third-party predictor for peptide safety and pharmacokinetic properties, was integrated to illustrate this extensibility.

The MCC scores reported below should be read as a baseline AMP classifier rather than a definitive predictor, establishing a reference against which future domain-specific predictors can be compared. For the AMP predictor, four strategies combining two encodings (HELM and SMILES) with two model families (sequence models and GNN architectures) were evaluated across multiple architectures and sizes (Supplementary Fig. S15, S16, with metric definitions in Section 4.8.3). Sequence models consistently outperformed GNNs by ~0.10 MCC (Fig. 4a), likely reflecting sequence-token models retaining positional context that node-edge conversion in graph models drops. Encoding had a smaller, family-dependent effect, with SMILES favoring sequence models and HELM favoring GNNs (per-cell ΔMCC in Supplementary Fig. S18), suggesting that monomer-level graph construction from HELM better aligns with the message-passing paradigm. To exploit this complementarity, predictions were aggregated via soft voting and MCC-weighted voting. Both aggregation strategies yielded comparable improvements (equal-weight 0.618, MCC-weighted 0.622, Fig. 4a), with the latter achieving a +0.035 improvement over the best individual model. The two aggregation strategies are statistically indistinguishable on the test set, with 95% confidence intervals (CIs) overlapping heavily. We adopt MCC-weighted voting as the production AMP predictor because it matches or exceeds soft voting on every reported metric. The complementary failure modes of the four ensemble members are detailed in Supplementary Fig. S17 and S19. This narrow margin between soft and weighted voting suggests that further headroom is more likely to come from new architectures or larger datasets than from additional refinements within the linear-pooling family of aggregators (Supplementary Section S5.4). Per-class analysis (Fig. 4b) shows the gain concentrates in active classes 1–4, while class 0 (the background class) shifts by approximately 0.01 (Supplementary Fig. S19).

**Fig. 4:**
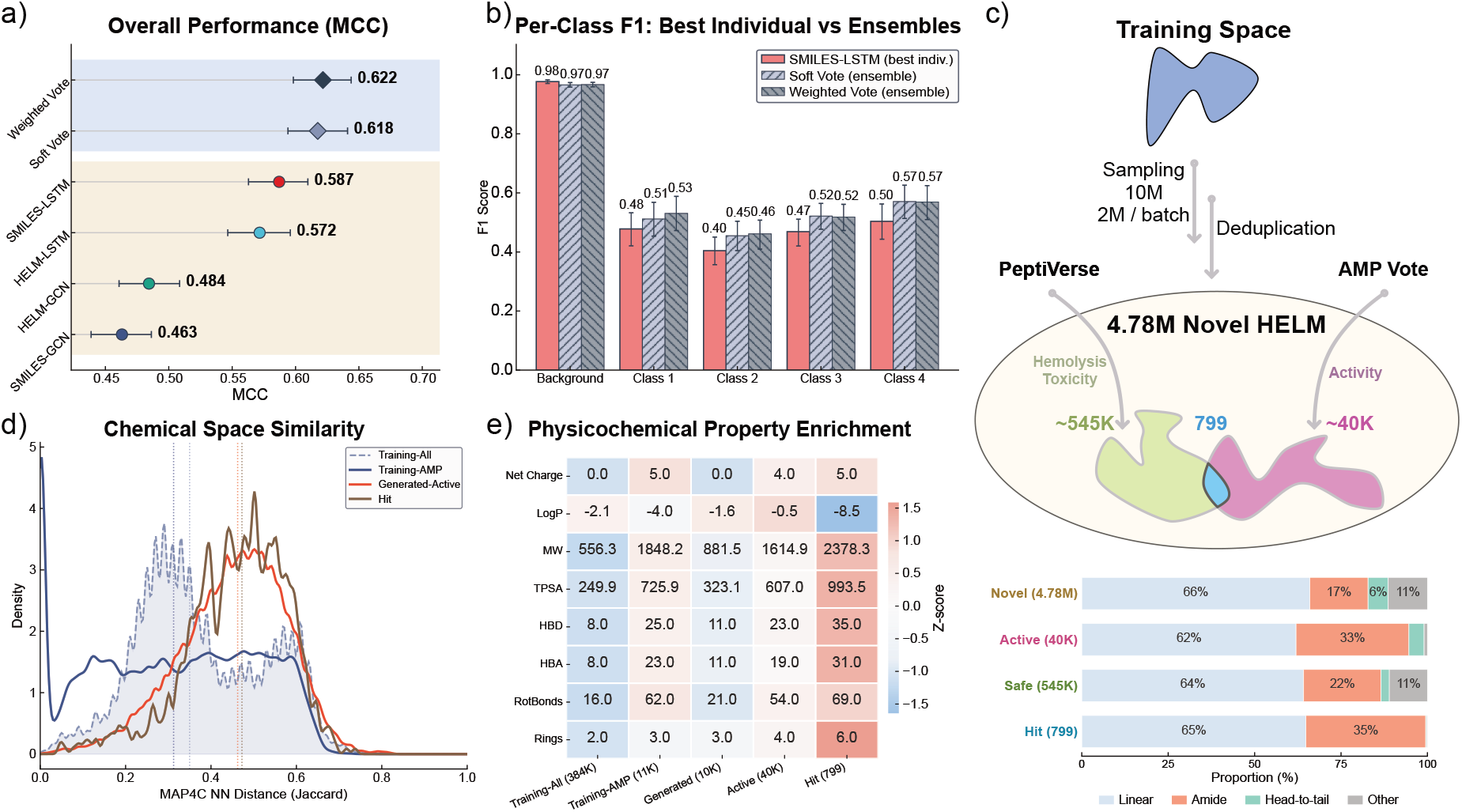
Prediction, screening, and candidate discovery. **(a)** Overall MCC for the four individual models (two encodings × two model families) and the soft-vote and MCC-weighted ensembles. Whiskers show 95% bootstrap CIs (1,000 paired resamples). **(b)** Per-class F1 for the best individual model (LSTM-SMILES) and both ensembles. **(c)** Screening pipeline. 10M sampled HELM peptides are deduplicated to 4.78M novel structures, then dual-screened by the AMP potency ensemble (~40K highly active, classes 3–4) and PeptiVerse safety filters (~545K non-toxic and non-hemolytic), yielding 799 hit candidates at the intersection. Bottom panel shows connection-type composition across the four screening stages. **(d)** Chemical-space novelty measured by MAP4C nearest-neighbour Tanimoto distance for four groups (all training peptides, training AMP subset, generated highly active candidates, and hit candidates). Training-All and Training-AMP show intra-set self-NN, while Generated-Active and Hit set show NN to Training-All. Dashed lines mark medians. **(e)** Physicochemical property enrichment across five sample populations. Medians coloured by row-wise Z-score (coral above row mean, blue below).

Applying this ensemble to large-scale screening, 10 million valid HELM peptides were sampled in five batches (2M each) from the LCC cascade pipeline, then deduplicated across batches and against the training set to yield 4.78 million novel unique structures. Two independent scoring criteria were applied in parallel. The AMP ensemble retained approximately 40,000 candidates in active classes 3 and 4 (predicted MIC *<* 32 *µ*g/mL), and PeptiVerse retained approximately 545,000 candidates as non-toxic and non-hemolytic. Their intersection yielded 799 hit AMP candidates (Fig. 4c). Through screening, amide connections enriched from 17% to 33% among active candidates, head-to-tail cyclization decreased modestly (6% → 4%), and all other connection types were almost eliminated (11% → 1%). This pattern tracks the composition of the AMP training subset, where amide chemistry overwhelmingly dominates and non-amide single-bond chemistries (head-to-tail rings, esters, and others) are vanishingly rare (Supplementary Fig. S3b), leaving the predictor with little training signal for non-amide motifs. Wet-lab assays will be required to test whether the predictor generalises beyond the training regime.

Chemical space analysis confirmed that the screened candidates occupy a distinct region of chemical space rather than recapitulating training structures (Fig. 4d). The nearest-neighbor MAP4C distance of generated highly active candidates (median 0.463) substantially exceeded the intra-set self-similarity of training AMP entries (median 0.313), and hit candidates remained at a comparable distance (median 0.473), confirming that the safety filter narrows the candidate set without collapsing it back onto training structures. Physicochemical property profiling across five sample populations (Fig. 4e) extended this picture. The most prominent transition occurred from the generated pool to highly active candidates. Median net charge rose from 0 to +4, consistent with the cationic character essential for electrostatic interaction with anionic bacterial membranes[21], while molecular weight (MW) and hydrogen bond donor count (HBD) increased substantially (881 → 1,615 Da and 11 → 23, respectively). LogP, by contrast, moved counter to the training AMP distribution, becoming less polar from generated to active (−1.6 → −0.5) rather than approaching the training AMP median (−4.0). Safety filtering then drove a second, distinct shift. Hit candidates became markedly less lipophilic (LogP −0.5 →−8.5) while retaining high net charge (+4 → +5), indicating that PeptiVerse filtered against hydrophobic surface area rather than against cationicity. The two screening axes act complementarily. The AMP ensemble pulls candidates toward the cationic, membrane-active regime, while PeptiVerse trims hydrophobic exposure within it, leaving a focused subset within the developable AMP window. The LCC cascade generation pipeline, combined with modular scoring, therefore navigates from millions of *de novo* HELM peptides to a focused set of structurally novel, physicochemically appropriate hit candidates warranting further usage.

## 3 Conclusion and Outlook

Modified peptides with special connections represent a pharmacologically important class that remains largely untouched by computational generation methods. PepForge addresses this gap through a three-stage LCC cascade that generates modified peptides in HELM format by decomposing the task into block layout, monomer content, and special connection prediction, trained on 383,817 HELM peptides curated via bidirectional S2H conversion. This hierarchical design supports *de novo* generation with faithful coverage of diverse connection types, masked infilling for targeted scaffold modification, and constrained generation where users can fix any structural level and let the model complete the rest. As a concrete demonstration, we paired it with a custom antimicrobial potency predictor and the external PeptiVerse[12] safety filter, applying the full pipeline to distill 4.78 million novel HELM peptides to 799 hits passing potency, hemolysis, and toxicity criteria. To support interactive exploration without scripting, we also built a web interface (Supplementary Section S6.3) that wraps the full pipeline.

The monomer library and special connection definitions are extensible, allowing users to register custom building blocks and define new bond types to broaden the accessible chemical space. The LCC generation cascade is task-agnostic by design, with two implications: the prediction module is a baseline screener limited by AMP training data dominated by linear unmodified peptides, and the LCC cascade itself is intended as a general-purpose HELM foundation usable beyond AMP screening, requiring large-scale sampling and post-hoc filtering at considerable computational cost. Building on this foundation, the pre-trained LCC generation cascade weights can be specialized through property-conditioned fine-tuning or reinforcement learning, directing generation toward specific targets and reducing this sampling overhead without rebuilding the pipeline from scratch. At a more fundamental level, integrating HELM-based sequence generation with three-dimensional structure prediction would enable target-aware design of modified peptides, bridging a gap that current structure-based methods cannot cross due to their limitation to canonical amino acids. The HELM data infrastructure, hierarchical generation framework, and open-source platform presented here provide a shared foundation for these advances and, we anticipate, will help bring the chemical space of modified peptides within systematic computational reach.

## 4 Methods

### 4.1 Data Collection and Curation

#### 4.1.1 Data Sources and Collection

Peptide information was collected from six public databases. PubChem[13] was queried via substructure search using a tripeptide backbone pattern (`O=C(C)NCC(NCC=O)=O`) to retrieve peptide-like SMILES; ChEMBL[11] was queried via application programming interface (API) for SMILES of peptides annotated as `protein` type; UniProt[22] provided reviewed short peptides (length 3–25 residues), converted to SMILES using RDKit[23]; DBAASP[14] provided single-chain antimicrobial peptide SMILES with measured MIC values; CycPeptMPDB[24] and Macrocycle-DB[25] were obtained as bulk downloads for SMILES. Salt fragments were removed by retaining the longest SMILES component, followed by InChIKey-based deduplication across all sources, yielding 1,461,836 unique peptides in SMILES format. Of these, 383,817 were successfully converted to HELM[10] strings via the S2H bidirectional pipeline described below, forming the final training dataset. Among the converted peptides, 11,026 entries from DBAASP with antimicrobial MIC values were used to train the antimicrobial potency prediction model. The number of peptides successfully converted from each source database and the MIC distribution of the DBAASP subset are reported in Supplementary Section S1.3 (Table S2, Fig. S3).

#### 4.1.2 SMILES2HELM Conversion Pipeline

To enable HELM-based model training from existing molecular databases, a bidirectional conversion pipeline between SMILES and HELM was developed. The forward direction adapts the backbone-detection and fragment-matching strategy of the `Structure2Sequence` module in the cyclicpeptide package[26], extended to emit HELM notation and to support special connection types beyond head-to-tail macrocyclization via a user-extensible SMARTS library. The reverse direction builds on the HELM-to-SMILES (H2S) reconstruction logic from HELM-GPT[5], extended to handle branched peptides and the same set of special connection types. A detailed diagram of the roundtrip pipeline is provided in Supplementary Fig. S1 (Section S1.1).

##### S2H conversion proceeds in four stages

(1) *Backbone detection*: the input SMILES is parsed by RDKit to identify the peptide backbone via amide bond patterns, with *α*-amino acid residues detected through iterative backbone walking from N-terminus to C-terminus; (2) *Monomer matching*: each detected residue is matched to known monomers in our curated HELM library (425 monomers, Supplementary Fig. S4) by InChIKey comparison, where the library includes canonical amino acids, non-proteinogenic amino acids, and chemical linkers, each annotated with R-group attachment points; (3) *Special connection detection*: eight types of special connections are identified using predefined SMARTS patterns, and head-to-tail macrocyclic linkages are detected when the C-terminal carboxyl group forms a peptide bond with the N-terminal amine (Supplementary Table S1); (4) *HELM assembly*: detected blocks (`PEPTIDE` and `CHEM`), monomer sequences, and special connections are assembled into valid HELM notation following the four-section HELM 2.0 format.

##### H2S reconstruction proceeds in three stages

(1) the HELM string is parsed to extract polymer blocks with their monomer SMILES sequences and connection records; (2) within each block, consecutive monomers are joined via peptide bonds using R-group atom mapping and the RDKit molzip algorithm; (3) cross-block special connections are formed by matching dummy atom pairs, removing leaving groups, and creating new single bonds. The final peptide is sanitized and converted to canonical SMILES.

##### Roundtrip validation

Each converted peptide undergoes a full SMILES-to-HELM-to-SMILES roundtrip. Conversion fidelity is verified by comparing the InChIKey of the original SMILES with that of the reconstructed SMILES. Peptides that fail this check are excluded from the dataset.

#### 4.1.3 Dataset Preparation

##### Monomer library

The monomer library contains 425 monomers adapted from the Pistoia Alliance HELM library[15], including all 20 canonical amino acids, non-proteinogenic amino acids, D-amino acids, N-methylated residues, and chemical connectors (Supplementary Section S2.1). Each monomer entry provides a canonical SMILES, an InChIKey identifier, and R-group definitions specifying valid attachment points and leaving groups. Custom monomers can be registered via the `Pipelines/Add_Monomer.py` utility, which enforces InChIKey deduplication. Special connection types are defined as pluggable SMARTS patterns, each specifying the substructure to match, the bond to cleave during fragment decomposition, and any heavy atom to restore at reconstruction (Supplementary Table S1).

##### Hierarchical data extraction

Each HELM peptide is parsed into three structural levels that correspond to the three generation stages. The layout level extracts block types and lengths. The content level extracts monomer sequences per block, with `CHEM` blocks retaining a reference to their preceding `PEPTIDE` context. The connection level records all special bonds as edges between monomer positions, annotated with R-group pairs and connection types.

##### Monomer embeddings

Pre-computed monomer embeddings (384 dimensions) were obtained from ChemBERTa-77M-MLM[16] using mean pooling over monomer SMILES token representations. These embeddings serve as optional initialization for token embedding layers in the *Content, Connection* stages, and prediction models, with a learned projection layer handling dimension mismatches. Beyond initialization, the embeddings provide chemically meaningful representations for rare or unseen monomers. The ChemBERTa featurization procedure, the physicochemical structure recovered by the embeddings, and the L2-norm rescaling applied before downstream use are described in Supplementary Section S2 (Figs. S5–S6).

##### Data splitting and tokenization

Both datasets (full HELM for generation, AMP subset for prediction) were randomly split into training/validation/test (80/10/10, seed 42). Distinct tokenizers were constructed for each model family from the training set. The *Layout* stage uses a structure tokenizer over block type and length tokens, the *Content* stage uses a monomer-symbol tokenizer, and the *Connection* stage operates on graphs and requires no tokenizer. The Flat GPT baseline uses a full-HELM tokenizer that retains block delimiters, monomer symbols, R-group labels, and connection notation as separate tokens. The prediction models support two encoding modes: a HELM mode that reuses monomer-level tokenization, and a SMILES mode that applies character-level tokenization.

### 4.2 Training Framework

All models share a unified training framework. The framework optimizes parameters using cosine annealing, with linear warmup applied over the first 5% of epochs when training for at least 100 epochs. Gradient clipping at maximum norm 1.0 and exponential moving average (EMA) of model parameters at decay 0.99 stabilize updates. Early stopping uses a patience of 20 epochs with strict improvement for generation models, and 15 epochs with a `min_delta` = 0.001 gate for prediction models. This asymmetry, together with per-stage best-checkpoint selection criteria, is detailed in SI Section S3.

PepForge provides unified pipeline runners (`Train_Generation.py` and `Train_Prediction.py`) that train multiple architecture and size combinations in a single invocation, automatically managing checkpoints, evaluation reports, and inference tests. All training was conducted on NVIDIA RTX 6000 Ada GPUs via SLURM job arrays. Default training settings shared across the three generation modules are listed in Supplementary Table S3, and the analogous defaults for the potency predictors in Supplementary Table S8.

### 4.3 Hierarchical LCC Cascade Generation Model

#### 4.3.1 Stage 1: *Layout*

The *Layout* stage generates the block-level structure as a token sequence specifying the type and length of each polymer block. The vocabulary contains 70 tokens (4 special, 2 block type, and 64 length tokens). Three autoregressive architectures were evaluated in a single small size variant: GPT[17], an LSTM[27], and a GRU[28]. GPT was selected for production based on the lowest validation loss. Architecture comparison and full hyperparameters are reported in Supplementary Section S3.1 (Table S4).

#### 4.3.2 Stage 2: *Content*

The *Content* stage generates monomer sequences for each block, conditioned on block type and target length from Stage 1. For multi-block peptides, `CHEM` blocks are further conditioned on preceding `PEPTIDE` blocks via a context-as-prefix mechanism. Four architectures (GPT, BERT[18], LSTM, GRU) were each evaluated in three size variants (Small, Medium, Large). Token embeddings were initialized from pre-computed ChemBERTa monomer embeddings, with a learned projection layer handling dimension alignment. The BERT variant is trained with random token masking using a wide rate range (15% to 85%, sampled uniformly per batch), enabling both partial infilling and nearly *de novo* generation at inference. GPT-large was selected as the production *de novo* Content backbone, and BERT-large as the production infilling backbone. Architecture*×*size sweep and full hyperparameters are reported in Supplementary Section S3.2 (Table S5).

#### 4.3.3 Stage 3: *Connection*

The *Connection* stage predicts special connections as a binary edge classification task on a molecular graph. Given a HELM peptide with filled monomer content from Stage 2, each monomer becomes a node and candidate edges are generated between all R-group-compatible monomer pairs. Node features combine monomer identity embeddings (ChemBERTa), block type, and positional encoding, while edge features encode the R-group pair type and the structural relationship (intra-block versus inter-block, terminal versus internal positions).

Five GNN architectures (GAT[29], GCN[30], GIN[31], MPNN[32], Graph Transformer[33]) were each evaluated in three size variants (Small, Medium, Large), and each architecture uses three hierarchical prediction heads. The GRAPH GATE first predicts whether the peptide contains any special connections (binary cross-entropy loss). For peptides that pass the gate, the EDGE EXIST GATE scores each candidate edge using binary focal loss (*γ*=2.0[34]) to handle the severe class imbalance between actual and candidate connections. Finally, the EDGE TYPE GATE classifies each predicted positive edge into one of nine special bond types using a weighted cross-entropy loss to address the long-tailed distribution of rare connection types.

To improve robustness against rare connection types, negative edges are dynamically resampled at each epoch using a deterministic per-graph seed, prioritizing “plausible-but-absent” pairs (monomer pairs that appear as positive connections in other training peptides) at a 4:1 negative-to-positive ratio. At inference, the three heads are applied sequentially with a fixed threshold of 0.5 for edge existence and argmax for edge type, followed by R-group constraint filtering against the HELM library to ensure chemical validity. GAT-large was selected as the production Connection backbone. GNN architecture comparison and full hyperparameters are reported in Supplementary Section S3.3 (Table S6).

### 4.4 Flat GPT Baseline

The Flat GPT baseline, an end-to-end GPT model, was trained to directly generate HELM strings as flat token sequences without hierarchical decomposition. It operates on a full-HELM vocabulary of 510 tokens (block delimiters, monomer symbols, R-group labels, and connection notation, see Section 4.1.3) and adopts the *Content* GPT-Medium architecture as a representative end-to-end baseline trained on the same dataset. The LCC cascade’s *Content* stage uses the larger GPT-Large backbone, so the comparison is conservative against the LCC cascade on raw validity. The reported gap (88.3% vs 99.6%) would, if anything, narrow with matched capacity, while the novelty and rare-type coverage advantages originate in the factorization rather than per-stage capacity. Full configuration and training hyperparameters are reported in Supplementary Section S7.1 (Table S10).

### 4.5 Antimicrobial Potency Prediction Model

#### 4.5.1 Task Formulation

Antimicrobial potency is predicted as a 5-class classification task based on MIC thresholds at inhibitory concentration [8, 32, 128] *µ*g/mL, applied regardless of bacterial strain or species (Supplementary Section S5.1): (0) background, no reported antimicrobial activity; (1) MIC *≥* 128 *µ*g/mL; (2) 32 *≤* MIC *<* 128; (3) 8 *≤* MIC *<* 32; (4) MIC *<* 8 *µ*g/mL. Class 0 is a positive-unlabeled (PU) negative sampled from peptides with no reported antimicrobial activity, and lies off the potency axis rather than below class 1. Routing OOD peptides here instead of forcing them onto the potency scale trades a small loss of recall for higher screening precision. To exploit the inherent ordering of MIC bins, the training loss uses focal cross-entropy (*γ*=2.0) with Gaussian-smoothed ordinal soft targets (*σ* =1.0)[35].

#### 4.5.2 Model Architectures

Eight base models were trained, spanning two encoding schemes (SMILES and HELM) and two model families. The language model family comprises GPT, LSTM, and GRU architectures applied to tokenized strings, with three size variants (Small, Medium, Large) per architecture. The GNN family comprises GAT, GCN, GIN, MPNN, and Graph Transformer applied in two modes: a HELM mode that operates on monomer-level graphs with ChemBERTa node features, and a SMILES mode that operates on atom-level graphs with Open Graph Benchmark (OGB)-style[36] atom and bond encoders. To mitigate the strong class imbalance between active and background peptides, each training epoch combines all active samples with an equal number of randomly sampled unlabeled peptides treated as the class-0 background. Architecture*×*size*×*encoding sweeps and full hyperparameters are reported in Supplementary Section S5 (Table S9).

#### 4.5.3 Ensemble Construction

The production AMP predictor uses a 4-model heterogeneous ensemble that combines the highest-MCC model from each (encoding *×* family) group: LSTM/Large (SMILES), LSTM/Medium (HELM), GCN/Large (HELM), and GCN/Large (SMILES). Predictions are aggregated by MCC-weighted soft voting, where each member’s validation-set MCC serves as its voting weight. The test set was held out from both member selection and weight assignment. Concretely, the ensemble class probability is

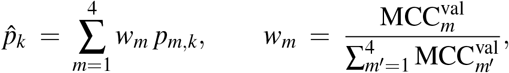

where *p*_*m,k*_ is member *m*’s predicted probability of class *k*, and the final prediction is 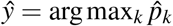. The soft-vote versus MCC-weighted comparison that motivates this choice is reported in Supplementary Section S5.4.

### 4.6 Inference Pipeline

The inference pipeline supports three entry levels: Level 0 (*de novo*, full LCC cascade), Level 1 (from a user-specified layout), and Level 2 (from a user-provided HELM with `[-]` mask tokens for infilling). Autoregressive stages use temperature sampling with top-*k* filtering.

Constrained generation can be applied at any stage and the constraints compose freely. Layout constraints restrict allowed block types, block counts, and per-block length ranges. Content constraints fix specific monomers at user-selected positions, routed automatically to autoregressive prefix-forcing for GPT/LSTM/GRU or to frozen-mask infilling for BERT. Forced connections specify a connection type and endpoint positions, which are merged with the *Connection* stage’s predictions and rejected when the predicted type confidence falls below a configurable threshold.

Generated peptides undergo asynchronous SMILES roundtrip validation using a producer-consumer architecture: a GPU-bound generation thread feeds a bounded queue consumed by multiple CPU-bound RDKit validation workers. Generation terminates when the target number of valid peptides is reached or a configurable budget is exhausted. For large-scale screening, a streaming mode writes accepted candidates to disk through callbacks instead of accumulating in memory, keeping memory usage independent of run size.

Potency prediction can be optionally enabled by specifying one or more task names, each auto-discovering the corresponding ensemble configuration. A predict-only mode is also supported, in which an existing CSV of HELM peptides is scored without generation. An `ExternalPredictor` interface enables integration of third-party scoring tools (e.g., PeptiVerse[12] for safety and pharmacokinetic properties). Per-task post-processing filters, ranking, and candidate truncation can be configured to produce a final candidate list.

All inference parameters are specified through a unified JSON configuration schema, with full field definitions, constraint injection points, and web-interface details provided in Supplementary Section S6.

### 4.7 Large-Scale Screening

As a downstream application of the LCC cascade, the inference pipeline was run at scale: five independent shards of 2 million peptides each (distinct random seeds) were generated, yielding 10 million novel peptides in HELM format. The LCC cascade was instantiated with the production checkpoints selected in Section 4.3. The AMP ensemble and PeptiVerse predictions were invoked inline per candidate via the multi-task prediction interface, and outputs were written in streaming mode to keep memory usage independent of run size.

After generation, the 10 million candidates were merged across shards. Deduplication by InChIKey removed both inter-shard duplicates and training-set overlaps, yielding 4,783,266 novel unique peptides. These were then passed through two parallel screening filters: AMP ensemble classification (active classes 3 and 4, predicted MIC *<* 32 *µ*g/mL) retained 39,891 candidates, and PeptiVerse safety prediction (non-toxic and non-hemolytic) retained 545,481 candidates. The intersection of both filters yielded 799 hit AMP candidates. The shard-level sampling commands, postprocessing, and candidate-analysis pipeline are documented in Supplementary Section S7.5.

### 4.8 Evaluation Metrics

This section defines the metrics used to assess generation quality, constrained-generation fidelity, and potency prediction performance, together with their interpretation.

#### 4.8.1 Generation Quality

Generation quality is reported on the *de novo* output of the LCC cascade and the Flat GPT baseline. Let *G* denote the set of raw HELM strings sampled from a model, *V* ⊆ *G* the subset passing roundtrip validation, *U* ⊆*V* the subset retained after InChIKey deduplication, and *T* the InChIKey set of the training peptides. All MAP4C-based metrics use MinHash fingerprints computed via the `mapchiral` Python package, which implements the MAP4C algorithm[19], a stereochemistry-aware extension of MAP4[20], configured with maximum radius 2 and 2,048 MinHash permutations and operating on canonical SMILES of roundtrip-valid peptides.

##### Validity

Validity is defined as |*V* |*/*|*G*|. A HELM string is counted as valid if it can be converted to a SMILES through H2S, parsed back through S2H, and the round-tripped HELM is structurally consistent with the original. Structural consistency is verified by labeled-graph isomorphism between the original and round-tripped HELM (Supplementary Section S1.4). The metric captures notation correctness and chemical realisability jointly. A drop in validity signals that the model is producing strings that fail at the parser, monomer-resolution, or bond-formation stage.

##### Uniqueness

Uniqueness is defined as |*U*|*/*|*V*|. Each valid SMILES is canonicalised and converted to an InChIKey, and duplicates within the batch are collapsed. Low uniqueness reveals intra-batch redundancy: the model repeatedly samples the same peptide, the typical signature of mode collapse or insufficient stochasticity in decoding.

##### Novelty

Novelty is defined as |*U\T*|*/*|*U*|, the fraction of unique generated peptides whose InChIKey is absent from the training set. It separates *de novo* synthesis from memorisation: a model that has overfit produces high validity and uniqueness but low novelty, because it reconstructs known structures rather than exploring new ones.

##### Internal diversity

Internal diversity is the mean pairwise MAP4C MinHash Jaccard distance over a random sample *P* of up to 10,000 unordered pairs from *U* :

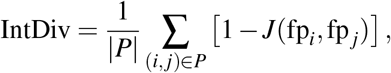

where 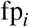 is the MAP4C fingerprint of peptide *i* and *J* the MinHash Jaccard similarity. Values close to 1 indicate that the batch spans heterogeneous chemical neighbourhoods, while values near 0 indicate that the peptides collapse into a narrow region. This metric is reported alongside the training set’s self-IntDiv as a reference distribution.

##### Nearest-neighbour MAP4C distance

For a generated peptide *x* and a reference set *R* (the training set), the nearest-neighbour distance is

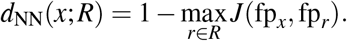

The reported statistic is the median of *d*_NN_ over all generated peptides. While IntDiv measures dispersion within the generated set, *d*_NN_ measures displacement from the training distribution. A median above the training set’s self-distance indicates that generations occupy regions of chemical space not covered by training data.

##### Per-connection-type metrics

Three statistics are computed per connection type *t* (amide, head-to-tail ring, disulfide, sulfanilamide, lanthionin, etc.). Each peptide contributes once per type regardless of multiplicity:

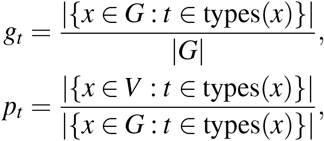

together with a per-type *d*_NN_ restricted to peptides containing *t*, computed against training peptides of the same type and capped at 500 samples per side. *g*_*t*_ exposes systematic under-sampling of rare connection types, *p*_*t*_ isolates per-type roundtrip fidelity from the overall validity, and per-type *d*_NN_ distinguishes models that satisfy a connection type by recalling training examples from those that explore new structures within that type.

#### 4.8.2 Constrained Generation Quality

Infilling metrics evaluate the *Content* stage’s ability to recover masked monomer positions while honouring user-supplied content constraints. For one infilled peptide, let *F* denote the set of free (non-constrained) positions, 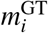 and 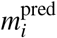 the ground-truth and predicted monomer at position *i*, and *s*(*a, b*) the cosine similarity between monomers *a* and *b* in the 384-dimensional ChemBERTa monomer embedding space defined in Section 4.2, with the convention *s*(*a, a*) = 1.

##### Validity (constrained)

Defined identically to *de novo* validity but applied to peptides generated under content constraints. It tests whether the model’s commitment to specific monomer positions still yields a chemically valid peptide after the connection stage and roundtrip validation.

##### Exact monomer accuracy

Per-peptide positional recovery rate over the free positions:

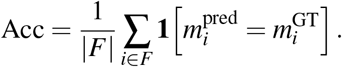

The mean of Acc across peptides at each reveal ratio is reported. This is the strictest position-level criterion. It does not credit chemically reasonable substitutions.

##### Compositional similarity

Per-peptide mean ChemBERTa cosine over all free positions:

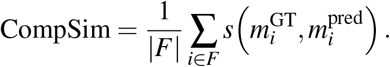

CompSim is a soft relaxation of exact accuracy: a substitution chemically close to the ground truth raises the score even when it does not match exactly. CompSim therefore captures the model’s ability to maintain compositional plausibility under low reveal ratios where exact recovery becomes statistically improbable.

##### Substitution similarity

Conditional cosine restricted to substitution events:

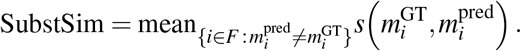

By excluding exact matches, SubstSim isolates the chemical judgment exhibited when the model deviates from the ground truth. A high SubstSim with a moderate Acc indicates a model that, when wrong, is wrong in chemically reasonable ways.

##### Perfect recovery rate

Molecule-level recovery rate:

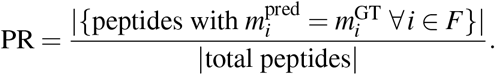

PR rises with reveal ratio for both architectures because, as more positions are fixed as context, fewer free positions remain and the task approaches verbatim recovery of the source peptide, making reproduction of entire training peptides increasingly likely. PR therefore co-varies with novelty: high PR at large reveal ratios partly reflects training-set recall rather than infilling accuracy.

##### Quality tier classification

Free positions are binned by their per-position cosine similarity into four tiers: exact (*s* = 1.0), high (0.9 *< s <* 1.0), moderate (0.7 *≤ s ≤* 0.9), and low (*s <* 0.7). Tier-wise differences (BERT−GPT) reveal whether one architecture concentrates its predictions in chemically tighter neighbourhoods, complementing the scalar similarity statistics.

#### 4.8.3 Antimicrobial Potency Prediction Quality

Potency prediction uses the 5-way ordinal MIC antimicrobial classification defined in Section 4.5. Let *y*_*i*_ and 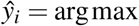 of the predicted logits be the ground-truth and predicted class for sample *i, N* the number of samples, *K* = 5 the number of classes, and *n*_*k*_ the support of class *k*. All metrics are computed via the scikit-learn[37] implementation.

##### Accuracy

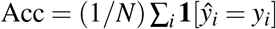. Overall fraction of correctly classified samples. It is sensitive to class imbalance and is dominated by the majority (background) class on AMP data, so it is reported only as a secondary reference.

##### Macro-F1

F1_macro_ = (1*/K*) ∑_*k*_ F1_*k*_, where F1_*k*_ is the harmonic mean of precision and recall on class *k*. Each class is weighted equally regardless of support, so the metric penalises failure on the rare active classes (3 and 4) that drive candidate prioritisation.

##### Weighted-F1

F1_weighted_ = ∑_*k*_(*n*_*k*_*/N*) F1_*k*_. The support-weighted variant of macro-F1, which reflects the expected F1 score on a sample drawn from the empirical class distribution. It is reported as a complement to macro-F1 and accuracy.

##### MCC

Multi-class Matthews correlation coefficient,

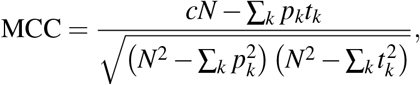

where *c* is the count of correctly classified samples, *t*_*k*_ the number of true class-*k* samples, and *p*_*k*_ the number of predicted class-*k* samples. The range is [−1, 1], with 1 indicating perfect agreement, 0 random, and negative values systematic disagreement. MCC remains balanced under class imbalance and penalises both intra-active confusion and background-to-active misassignments. It is the primary metric for selecting ensemble members and ranking individual predictors.

### 4.9 Software Implementation

PepForge is implemented in Python, with model training and inference built on PyTorch[38] and PyTorch Geometric[39] for sequence models and GNNs. Molecular processing relies on RDKit, MAP4C fingerprints on the `mapchiral` package, ChemBERTa monomer embeddings on the HuggingFace Transformers library[40], and classification metrics on scikit-learn.

The local web interface is built on a FastAPI backend that wraps the LCC cascade generation pipeline, AMP prediction ensemble, and PeptiVerse predictor as REST endpoints, paired with a React and TypeScript frontend (Vite and Tailwind CSS). The application runs entirely on the user’s machine and requires no cloud service.

## 5 Code and Data Availability

All source code, pre-trained models, training data, generated peptides, and figure-reproduction materials are released at https://github.com/wqx1999/PepForge, together with a one-click installer and a local web interface for interactive use. HuggingFace mirrors for the model weights and dataset archives are linked from the repository README. Code is released under the MIT license, and models and data under the CC-BY-4.0 license.

## 6 Acknowledgments

Qingxin Wang is supported by China Scholarship Council (CSC202408320133). The authors thank Dr. Erik Jung for his valuable review and helpful comments. The project was supported by grants from the Deutsche Forschungsgemeinschaft (DFG, RTG 2473 “Bioactive Peptides”, project number 392923329, R. D. S.).

