## Supplementary file for "PepForge: Hierarchical HELM-Based Peptide Generation"

### Supplementary Information
### PepForge: Hierarchical HELM-Based Peptide Generation

Qingxin Wang[1] Roderich D. Süssmuth[1,*]

[1]Institut für Chemie, Technische Universität Berlin, Straße des 17. Juni 115, D-10623 Berlin, Germany

**References** **47**

### S1 Data Curation and Conversion Pipeline

#### S1.1 SMILES-to-HELM Roundtrip Pipeline

The curated training set is built by a bidirectional SMILES-to-HELM[1] conversion pipeline with InChIKey-based roundtrip validation (Fig. S1). The forward direction adapts the backbone-detection and fragment-matching strategy of the `Structure2Sequence` module in the cyclicpeptide package[2]. We extend that strategy in three directions to fit our setting: (1) the output target is HELM notation instead of a one-letter sequence, so an additional HELM Construction stage is added to order monomers along the backbone and emit connection annotations, (2) fragments that fail direct monomer matching are routed through a special-connection search against a user-extensible SMARTS library, enabling coverage of non-peptide linkers (disulfide, lactone, thioether and five other connection types) beyond head-to-tail macrocyclization, and (3) a matching reverse direction (`HELM2SMILES`), adapted from the HELM-to-SMILES reconstruction logic of HELM-GPT[3] and extended to handle branched peptides and our full set of special connection types, is implemented so that the whole pipeline can be closed with an InChIKey roundtrip gate. The orchestrator `SMILES_HELM_roundtrip` drives each peptide through `SMILES2HELM` and then `HELM2SMILES`, retaining only entries whose reconstructed structure yields the same InChIKey as the input.

The forward direction decomposes a peptide into four stages (Fig. S1a). (1) *Molecule Preparation* parses the input, detects a peptide-like backbone, and rewrites the atom ordering into a canonical head-to-tail direction. (2) *Bond Break Points* enumerates cleavable peptide bonds along that backbone and any head-to-tail ring closure. (3) *Monomer Composition* matches every resulting fragment against the 425-entry monomer library. Fragments that still carry a non-peptide linkage are routed through a special-break search that tries to split them along one of the eight SMARTS-defined bond templates and then re-matches each sub-fragment against the same library. (4) *HELM Construction* reorders the matched monomers along the backbone and collects the inter-fragment connections into HELM annotations.

The reverse direction rebuilds the peptide in three stages (Fig. S1b). (1) *HELM Parsing* splits the HELM string into polymer blocks and connection specifications. (2) *Atom Mapping* walks each connection and assigns R-group indices to the monomer atoms flagged as leaving groups, so that downstream bond formation attaches at the correct position. (3) *Bond Formation* joins monomers with peptide bonds, reconstructs the special connections, and emits the final SMILES.

The pipeline is intentionally restricted in scope. Backbone detection in the forward direction matches the $\alpha$-amino acid pattern `C(=O)CN` only. As a consequence, molecules built on $\beta$- or $\gamma$-amino acid backbones, or on non-amide main-chain linkages, cannot be parsed with current pipeline. The forward direction also searches for a single continuous linear backbone and rejects branched topologies in which one residue's side chain extends into a second sub-chain. The library of special cross-links is fixed at the eight SMARTS patterns of Table S1 plus a head-to-tail ring rule (Fig. S1d), and a molecule requiring a bond type outside this set is rejected at the matching stage. Extending coverage to new chemistries reduces to adding entries to the monomer or SMARTS libraries (procedure in Section S7.6). These are scope limits rather than failure modes: they explain why only part of the raw data survives the conversion (Section S1.3), but they also guarantee that every retained peptide decomposes cleanly into three design primitives (an $\alpha$-amino acid backbone, a library monomer, and a library cross-link) and passes the

InChIKey roundtrip gate (Fig. S1c) unchanged.

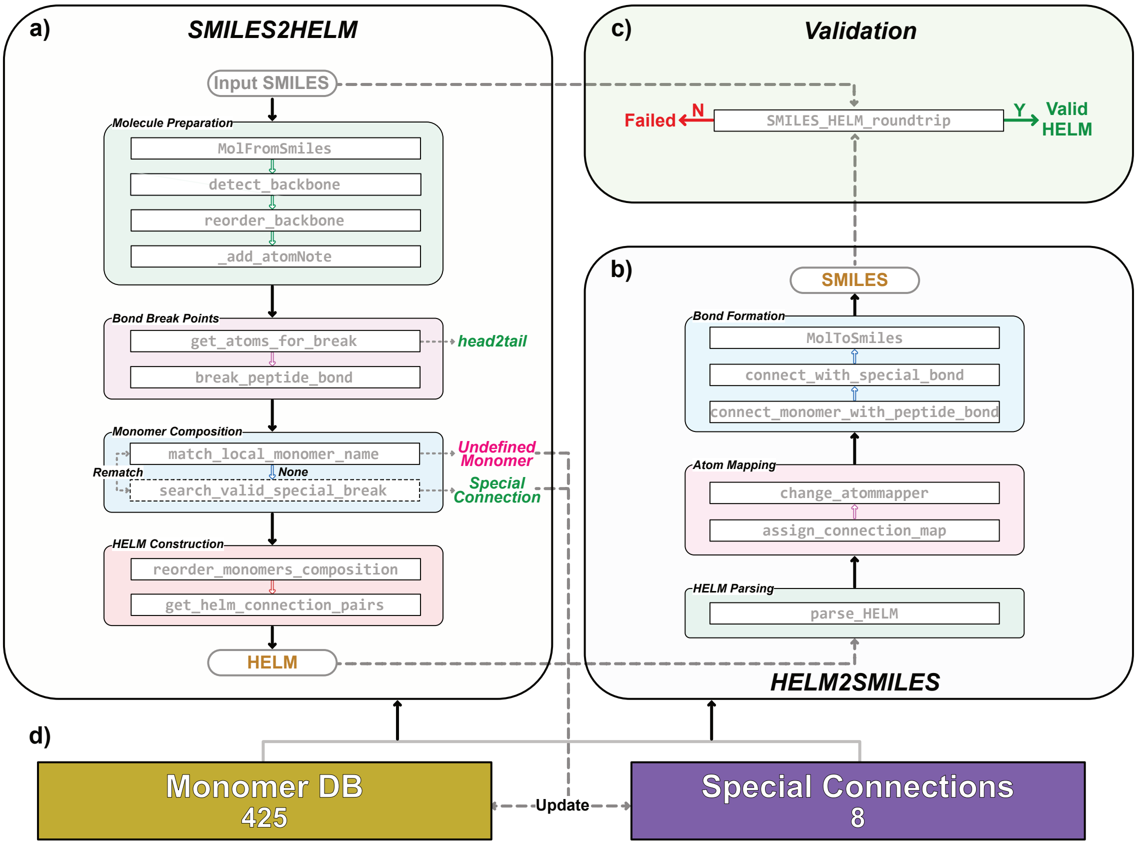

**Fig. S1: Bidirectional SMILES-to-HELM conversion and roundtrip validation pipeline.** **(a)** SMILES2HELM converts an input peptide through four sequential stages. *Molecule Preparation* parses the SMILES and canonicalizes the peptide backbone. *Bond Break Points* enumerates cleavable peptide bonds and head-to-tail closures. *Monomer Composition* matches each fragment against the monomer library, routing fragments that contain special bonds through a SMARTS-based special-break search and re-matching the resulting sub-fragments. *HELM Construction* orders the matched monomers along the backbone and assembles the connection annotations. **(b)** HELM2SMILES reverses the process in three stages. *HELM Parsing* splits the string into polymer blocks and connection records. *Atom Mapping* assigns R-group indices to the monomer leaving atoms. *Bond Formation* joins monomers with peptide bonds and reconstructs the special connections. **(c)** The orchestrator SMILES_HELM_roundtrip compares the InChIKey of the input SMILES and the SMILES reconstructed from the generated HELM. Only peptides passing this check are retained in the training set. **(d)** Both directions consult a shared 425-entry monomer library and an 8-entry SMARTS library of special connection types, both user-extensible. Individual function names of each stage are annotated in the corresponding figure box.

#### S1.2 SMARTS Patterns for Special Connection Detection

Table S1 specifies the eight SMARTS patterns that the SMILES2HELM forward direction uses to detect non-peptide linkages during fragment decomposition. Each entry provides a substructure match that flags the bond of interest, a *Break* tuple $(i, j)$ identifying the pair of SMARTS match atoms whose bond is cleaved, an optional *Add atom* tuple $(X, k)$ specifying an atom $X$ that is restored at match position $k$ during HELM2SMILES reconstruction so that forward decomposition and reverse rebuilding remain exact inverses, and a short chemical context indicating the typical use of the pattern. Together the eight patterns cover amide and carbamate linkers, sulfonamides, disulfides, esters and lactones, indole-thioether Trp–Cys cross-links, lanthionine bridges, and benzyl-thioether staples. Head-to-tail ring closure is handled separately by a backbone-adjacency rule rather than a SMARTS match, since its signature (a peptide bond between the C-terminal carboxyl carbon and the N-terminal amino nitrogen along the canonicalized backbone) depends on the full peptide topology rather than a local substructure.

**Table S1: SMARTS patterns for the eight special connection types.**

| Name | SMARTS | Break[a] | Add atom[b] | Chemical context |
|---|---|---|---|---|
| Amide | `[c,C]-C(=O)-[N;!$([N](-[C](=O))-[C](=O))]` | $(1,3)$ | (O, 1) | Amide bonds in non-peptide blocks |
| PG_Amide | `OC(=O)-[N;!$([N](-[C](=O))-[C](=O))]` | $(1,3)$ | (O, 1) | Carbamate linkages (Boc, Fmoc) |
| Sulfanilamide | `[c,C]-S(=O)(=O)-[N]` | $(1,4)$ | (O, 1) | Sulfonamide cross-links |
| Disulfide | `S-S` | $(0,1)$ | — | Cys–Cys disulfide bridges |
| Ester | `[c,C]-C(=O)[O;$([O]-[#6;!$(C=O)])]` | $(1,3)$ | (O, 1) | Ester and lactone cyclizations |
| Trp-Cys | `[S]-[c;$(*~[nH])]` | $(0,1)$ | — | Indole-thioether Trp–Cys cross-link |
| Lanthionin | `NC[C;H2]-[S]-CCN` | $(3,4)$ | (H, 4) | Lanthionine bridges (Dha/Dhb–Cys) |
| Bz_stapled | `[c]-[CH2]-S-C` | $(1,2)$ | — | Benzyl-thioether stapled peptides |

[a] $(i, j)$: the bond between atoms at zero-based SMARTS match positions $i$ and $j$ is cleaved during fragment decomposition.
[b] $(X, k)$: atom $X$ is restored at position $k$ during reconstruction. "—" denotes no restoration.

Fig. S2 walks through this convention on the *Amide* entry applied to a representative training peptide. The SMARTS matches an amide in a non-peptide context and assigns zero-based indices 0–3 to its four atoms in match order. The *Break* field $(1,3)$ cleaves the bond between atoms 1 (the carbonyl carbon) and 3 (the amide nitrogen), splitting the peptide into two fragments, and the *Add atom* field $(O,1)$ instructs `HELM2SMILES` to reinstall an oxygen at position 1 during reconstruction, recovering the original amide. *Add atom* is only needed when RDKit[4]'s sanitization cannot restore an atom on its own: heavy atoms lost at the cleavage site must always be listed, and hydrogens must be specified when they sit on or next to a stereocenter, since RDKit does not re-assign implicit hydrogens across stereo-tagged atoms. Lanthionin is the only entry where this applies. Ordinary hydrogens elsewhere are restored automatically.

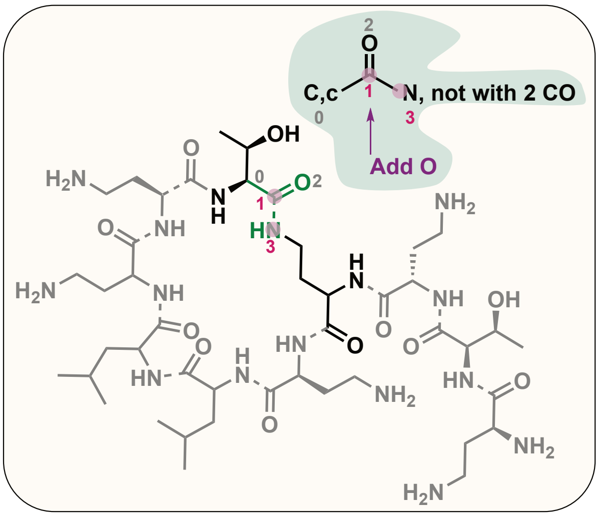

**Fig. S2: Worked example of SMARTS-based special connection detection on the *Amide* entry from Table S1.** A representative training peptide is shown with one amide match highlighted; the top-right inset is the SMARTS pattern with atom indices 0–3. Green shading marks atoms belonging to the SMARTS match; purple circles mark the two atoms whose bond is cleaved at this step; the purple *Add O* arrow denotes the oxygen atom restored during `HELM2SMILES` reconstruction. Dashed bonds elsewhere in the peptide would indicate bonds already broken by earlier decomposition steps. The seven remaining connection types in Table S1 follow the same visual convention.

#### S1.3 Data Sources, Conversion Outcomes, and Antimicrobial Potency Distribution

Table S2 summarizes per-source collection and pipeline-level conversion outcomes for the training dataset. Six public databases were queried through heterogeneous access routes, all wrapped in the data-fetch utility `Pipelines/Fetch_Peptides.py`. (1) PubChem[5] was searched through its PUG-REST substructure endpoint using the tripeptide backbone SMARTS `O=C(C)NCC(NCC=O)=O`, returning peptide-like compound identifiers (CIDs) and their canonical SMILES. (2) ChEMBL[6] was queried through its official Python client with the filter `molecule_type=Protein`, returning ChEMBL identifiers and canonical SMILES. (3) DBAASP[7] was retrieved entry-by-entry through its public REST API, restricted to single-chain peptides with measured minimum inhibitory concentration (MIC) values, returning DBAASP identifiers, SMILES, and MIC values. (4) UniProt[8] was queried through its REST search endpoint for reviewed (Swiss-Prot) entries of length 3–25 amino acids, returning UniProt accession numbers and one-letter amino acid sequences. (5) CycPeptMPDB[9] was obtained as a bulk CSV download from its website, providing molecular identifiers and SMILES for curated cyclic peptide entries. (6) Macrocycle-DB[10] was likewise obtained as a bulk CSV download, providing molecular identifiers and SMILES for its macrocycle entries. Collectively the six sources contributed 1,611,302 records, of which PubChem alone accounted for roughly 96% (Table S2).

All records were then funneled through a common cleanup and conversion pipeline exposed as `Pipelines/Processing_Raw_Data.py`. (1) UniProt sequences were converted to SMILES using `RDKit.Chem.MolFromSequence`. Sequences containing non-standard residues that could not be resolved by RDKit were silently dropped. (2) Salt fragments were removed from every SMILES by splitting on the "`.`" separator and retaining the longest component. (3) The pooled set was deduplicated by InChIKey, keeping the first occurrence across sources, which yielded 1,461,836 unique molecules. (4) The deduplicated set was passed through the `SMILES2HELM` pipeline described in Section S1.1. Of these, 383,817 entries (23.82% of the raw total and 26.26% of the pooled unique set) satisfied the InChIKey roundtrip gate and form the final training set. The DBAASP/MIC subset was processed through the same pipeline and, after restricting to entries with a numeric MIC value, yielded 11,026 potency-tagged HELM entries (76.0% of the 14,509 raw DBAASP records) for potency prediction. The same fetch-and-convert pipeline applies when users supply a new raw dataset for retraining (Section S7.6).

**Table S2: Data sources and pipeline-level conversion outcomes for the training dataset.**

| Source | Criteria | Count | Fraction (%) |
|---|---|---:|---:|
| PubChem | Substructure search with tripeptide backbone | 1,550,241 | 96.21 |
| ChEMBL | API query, `molecule_type=Protein` | 22,241 | 1.38 |
| DBAASP[a] | Single-chain peptides with SMILES and MIC | 14,509 | 0.90 |
| Macrocycle-DB | Bulk download | 11,077 | 0.69 |
| CycPeptMPDB | Bulk download | 8,466 | 0.53 |
| UniProt | Reviewed, length 3–25 aa (sequences) | 4,768 | 0.30 |
| Raw total | Sum across all sources | 1,611,302 | 100.00 |
| Pooled unique | UniProt sequence → SMILES, salt removal, InChIKey dedup | 1,461,836 | 90.72 |
| HELM conversion[b] | Training set | 383,817 | 23.82 |
| HELM conversion[c] | DBAASP / MIC subset (numeric MIC) | 11,026 | 76.00 |

[a] Only source providing measured antimicrobial activity (MIC values).

[b] Full training set used for the generation model.

[c] DBAASP subset used to train the antimicrobial potency prediction model, restricted to entries with a numeric MIC value. Fraction is computed relative to the 14,509 raw DBAASP records rather than the raw total across all sources.

Fig. S3 summarizes the composition of the training database at the monomer, connection, and activity levels. At the monomer level (Fig. S3a), 34 monomers meet the minimum-count cutoff of 100 in both sets. The six with the highest AMP enrichment are dH, dI, K, dK, I, and W, with $\log_2$(AMP/All) ratios ranging from +1.17 to +2.57. The six most depleted are dP, ac, Dab, dF, C, and D, with ratios ranging from −0.68 to −1.79. At the connection level (Fig. S3b), 59.2% of AMP peptides carry at least one special bond compared with 22.7% in the All-Peptides set, a 2.6× peptide-level enrichment. The per-type breakdown shows that amide chemistry accounts for 98.3% of special-bond instances in AMP and 63.1% in All-Peptides. The remaining 1.7% in AMP and 36.9% in All-Peptides comprise ester, disulfide, head-to-tail, and other non-amide chemistries. At the activity level (Fig. S3c), the DBAASP subset spans a wide MIC range. 18.1% of entries fall in the most potent bin (MIC ≤ 8 μg/mL), 12.9% to 16.4% in each of the five bins spanning 8 to 256 μg/mL, and 3.2% to 5.1% in the two highest MIC bins (> 256 μg/mL).

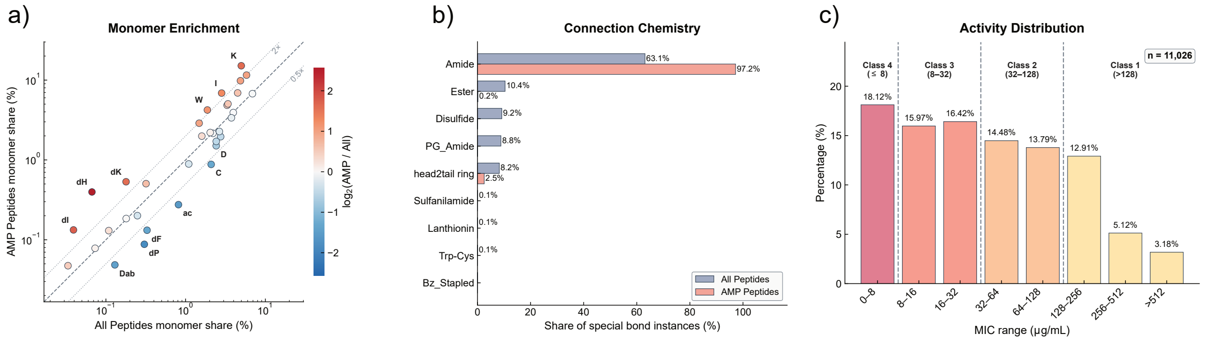

**Fig. S3: Composition of the HELM training database.** **(a)** Log-log scatter of monomer share in the All-Peptides set ($x$-axis) versus the AMP-Peptides subset ($y$-axis). Each point represents one monomer, and only monomers with at least 100 occurrences in both sets are shown. Colors encode $\log_2$(AMP share/All share) on a diverging blue–white–red scale. The dashed diagonal marks the no-enrichment baseline, and the dotted lines mark the 2× and 0.5× bounds. The six most enriched and the six most depleted monomers are labeled. **(b)** Share of special-bond instances per connection type for All-Peptides (blue) and AMP-Peptides (coral), computed across the eight SMARTS patterns defined in Table S1 plus the head-to-tail ring rule. **(c)** Distribution of MIC values in the DBAASP subset ($n$ = 11,026). Eight $\log_2$-spaced MIC bins are grouped into the four downstream potency classes by the thresholds 8, 32, and 128 μg/mL, colored with the yellow-to-red gradient used in the main-text potency prediction figure (deep red denotes the most potent class).

#### S1.4 Labeled-Graph Isomorphism for Generation Validity

Training-data curation (Section S1.1) accepts an entry only if the InChIKey of its reconstructed SMILES matches the InChIKey of the input SMILES, because both endpoints are SMILES strings. At generation time, however, the input is a HELM string rather than a SMILES, and the natural endpoints of the roundtrip are HELM → SMILES → HELM. Comparing the two HELM strings character-by-character is too strict (the same cyclic peptide admits multiple equivalent rotations of its monomer ordering, all of which should be accepted), while collapsing through InChIKey would discard HELM-level annotations such as connection-type labels that are part of what the generator is asked to produce. We therefore replace the InChIKey check with labeled-graph isomorphism on the monomer-level graph implicit in each HELM string.

Each HELM string is parsed into an undirected graph $G = (V, E, \ell_V, \ell_E)$. Each node $v \in V$ corresponds to one monomer occurrence and carries the monomer HELM symbol as its label $\ell_V(v)$ (e.g. `A`, `dC`, `Mod1`). Each edge $e \in E$ connects two monomer nodes that share a backbone or special bond and carries the connection-type string as its label $\ell_E(e)$ (one of `backbone` or the eight special-bond types of Table S1).

To make the labeling rotation-invariant on cyclic peptides, the head-to-tail amide bond closing a ring is renormalized from `R1-R2` to `backbone` so that all rotations of the same ring share an identical edge labeling. Only true head-to-tail (R1-R2) closures are renormalized, while disulfide and side-chain cross-links between the first and last residue keep their original labels. Two HELM strings are declared structurally consistent if and only if their parsed graphs are isomorphic under both node and edge labels. Formally, $h_a$ and $h_b$ are consistent iff there exists a bijection $\phi : V_a \to V_b$ such that $\ell_V^a(v) = \ell_V^b(\phi(v))$ for every $v \in V_a$, $(u,v) \in E_a \iff (\phi(u),\phi(v)) \in E_b$, and $\ell_E^a(u,v) = \ell_E^b(\phi(u),\phi(v))$ for every edge.

Generation validity in Section 4.8.1 of the main text requires both that the input HELM round-trips through SMILES and that the resulting HELM passes the labeled-graph equivalence defined above. Failures of either condition exclude the candidate from $V$.

Backbone renormalization preserves the cyclic-vs-linear distinction: a cyclic $n$-mer has $n$ backbone edges while a linear $n$-mer has $n-1$, and isomorphism requires equal edge counts.

### S2 Monomer Embeddings

#### S2.1 Monomer Library and ChemBERTa Featurization

The monomer library used by the SMILES-to-HELM[1] pipeline of Section S1.1 was seeded from the HELMCoreLibrary, the public monomer library distributed with the HELM software by the Pistoia Alliance[11]. From the seed we retained only entries within peptide chemistry and edited the SMILES of selected monomers to expose the side-chain atoms recruited by our special-connection library (for example, the lanthionine thioether of Ala variants whose side-chain position serves as one end of a lanthionine bridge). We then added a set of terminal capping groups frequently used in solid-phase peptide synthesis and enriched every entry with an InChIKey for downstream deduplication. The curated library is treated as open-ended: monomers that recur in the forward pipeline but are not yet covered can be added in batches. The user lists new monomers in a CSV (symbol, SMILES, monomer type, natural analog, R-group cap groups) and runs the one-click utility `Pipelines/Add_Monomer.py`, which deduplicates the input against `Configs/Monomer/HELMLibrary.json` by InChIKey and assigns a continuous identifier to each accepted entry. After several rounds of this scan-and-extend cycle over the pooled training set, the library converged at 425 monomers, consisting of 344 backbone residues, 76 terminal groups, and 5 link monomers. Panels (a)–(d) and (f) of Fig. S4 together cover the 344 backbone residues, while panel (e) covers the 81 capping groups and linker connectors (76 terminal + 5 link, 81 entries).

Fig. S4 partitions the full 425-entry library into six groups. The largest group comprises mixed type amino acids, non-proteinogenic residues with side-chain or scaffold modifications relative to the canonical 20 amino acids ($n = 208$, e.g., homocysteine and 3-thienylalanine). The 37 stereochemistry-unspecified amino acids (asterisk-prefixed HELM symbols, Fig. S4d) capture chirality omissions in the source SMILES and remain connectivity-equivalent to their L- and D-counterparts. The model treats them as a separate token class so that downstream sequences inherit the source ambiguity rather than silently committing to either configuration.

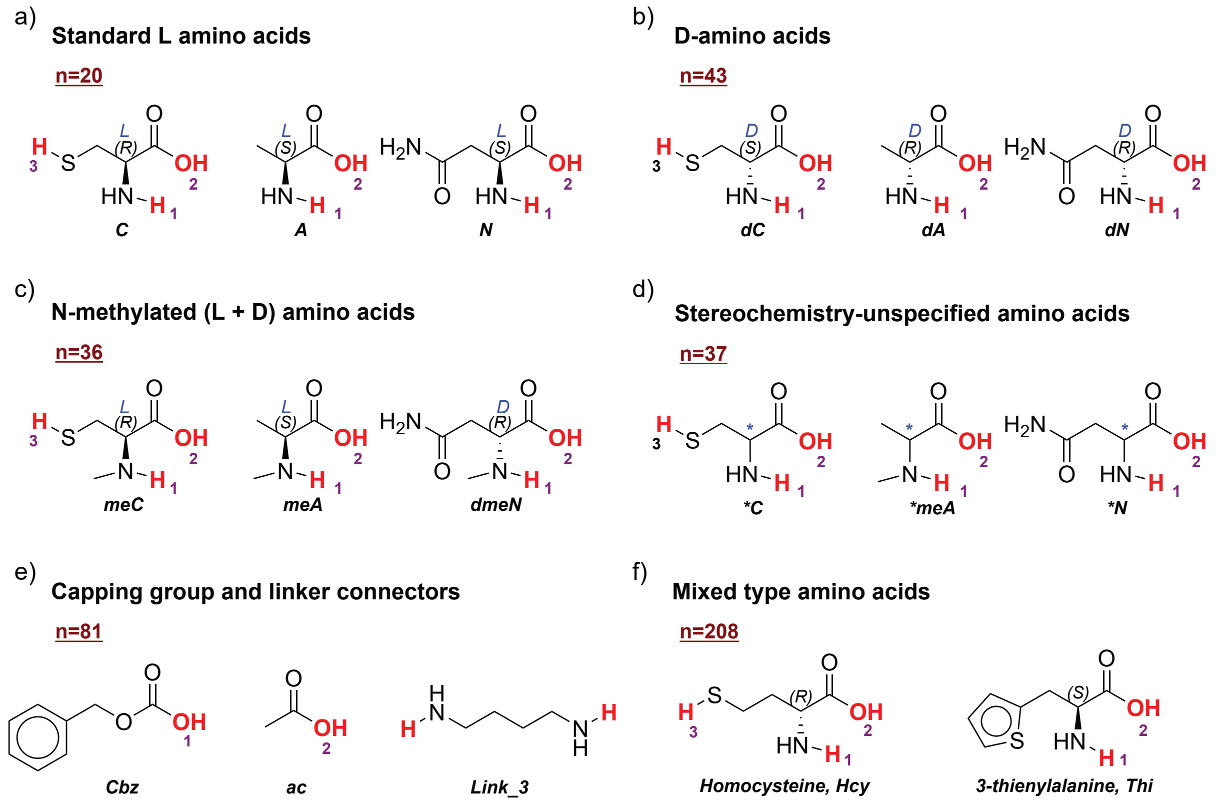

**Fig. S4: Chemical composition of the 425-monomer library.** The 425 entries are generally divided into six chemical classes: **(a)** standard L amino acids ($n = 20$); **(b)** D-amino acids ($n = 43$); **(c)** N-methylated (L + D) amino acids ($n = 36$); **(d)** stereochemistry-unspecified amino acids (asterisk-prefixed HELM symbols) whose source SMILES carries no explicit chirality annotation ($n = 37$); **(e)** capping groups and linker connectors ($n = 81$); and **(f)** mixed type amino acids, non-proteinogenic residues with side-chain or scaffold modifications relative to the canonical 20 ($n = 208$). Two to three representative structures are shown per class, with R-group attachment positions annotated as $H_1/H_2/H_3$ following HELM library conventions.

Each monomer is featurized into a continuous vector with ChemBERTa[12], a pre-trained chemical language model for SMILES. These vectors initialize the monomer embedding tables of the three generation modules and of the HELM-based AMP predictor (Section S5), which are then updated jointly with the rest of each host model during training. The motivation for pre-training over learning from scratch is a coverage gap between the two training sets: the generation set covers 386 of the 425 library monomers, whereas the AMP prediction set covers only 283. The 103 monomers seen in generation but not in AMP training are in-distribution for the peptides the generation model emits at inference, yet their rows in the AMP predictor never receive a gradient update. Random initialization would in principle leave those rows as noise at scoring time, whereas ChemBERTa initialization keeps them chemically coherent by construction. We use the `DeepChem/ChemBERTa-77M-MLM` checkpoint loaded through the HuggingFace Transformers library[13], feed each monomer's canonical SMILES through the associated tokenizer, and mean-pool the last hidden states of the non-padding tokens into a single 384-dimensional vector. The collected vectors and their monomer symbols are stored in `Configs/Embedding/Artifacts/monomer_embeddings.pt`. Tokens absent from the file (special tokens and any future monomers added after embedding generation) fall back to random initialization.

#### S2.2 Embeddings Encode Physicochemistry

Fig. S5 shows the 425 ChemBERTa embeddings on a shared $t$-SNE[14] layout, with each of its eight panels colored by one RDKit[4] physicochemical descriptor: molecular weight, LogP, topological polar surface area, hydrogen-bond donor count, hydrogen-bond acceptor count, number of rotatable bonds, number of aromatic rings, and fraction $CSP^3$. Monomers for which RDKit cannot compute the full descriptor panel are drawn in light grey at the background layer. The panels show an imprecise but visible trend: monomers with similar descriptor values tend to colocate, most clearly along an aromaticity and hydrophobicity axis, and more loosely for the other descriptors. ChemBERTa thus provides a partial chemical prior rather than a clean physicochemical encoding, which is still preferable to random initialization for the AMP-unseen rows of Section S2.1.

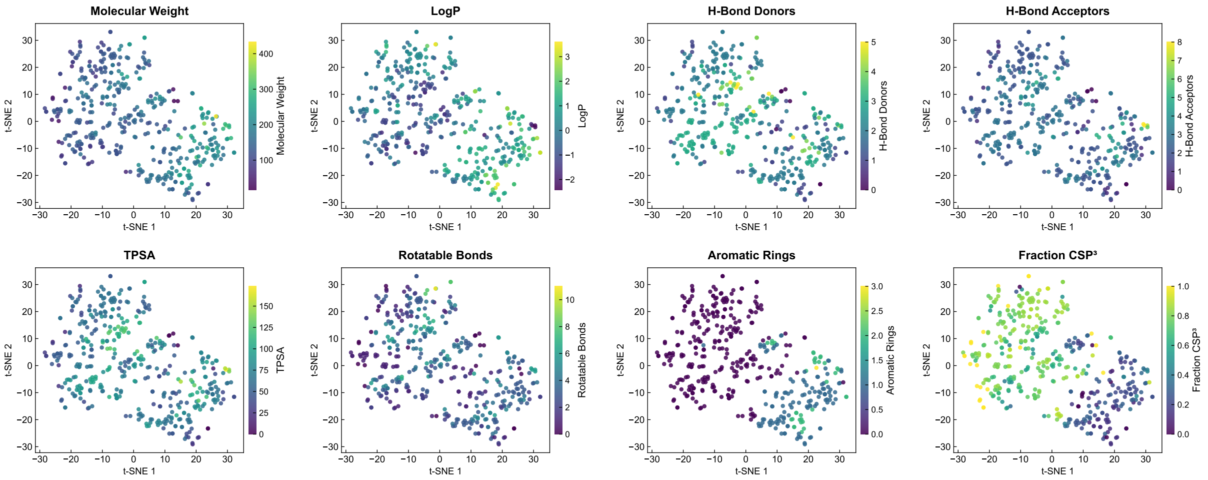

**Fig. S5: ChemBERTa embeddings organized by RDKit physicochemical descriptors.** Two-dimensional $t$-SNE projection of the 384-dimensional ChemBERTa embeddings for the 425 monomers in the library. Each panel is colored by one RDKit descriptor: molecular weight, LogP, topological polar surface area, hydrogen-bond donors, hydrogen-bond acceptors, number of rotatable bonds, number of aromatic rings, and fraction $CSP^3$. All panels share the same $t$-SNE coordinates; only the color channel differs. Monomers for which RDKit cannot compute the full descriptor panel are drawn in light grey at the background layer.

#### S2.3 Rescaling Before Downstream Use

Applied to the 425 library monomers, the ChemBERTa procedure of Section S2.1 returns a stacked tensor with a standard deviation of approximately 0.19, corresponding to an average per-row L2 norm of about 3.7. This is roughly five times smaller than the expected L2 norm of a PyTorch[15] default `nn.Embedding` row drawn from $\mathcal{N}(0,1)$, which is $\sqrt{384} \approx 19.6$. Downstream models build their monomer embedding layer from this tensor and fall back to a random $\mathcal{N}(0,1)$ initialization for any token without a matching entry. The resulting magnitude mismatch would bias early-training activations and gradients toward the larger random rows, suppressing the structural prior that ChemBERTa contributes. We therefore rescale the tensor to unit standard deviation once at embedding-generation time, before writing it to disk. To keep the table in sync with the HELM library, `Pipelines/Add_Monomer.py` runs the same procedure for every newly added monomer, computing its ChemBERTa vector, rescaling with the `raw_std` recorded in the embedding manifest, and appending the new row after the existing ones so that earlier checkpoints stay reproducible.

Fig. S6a confirms that the rescaling places the ChemBERTa distribution onto the `nn.Embedding` reference scale. The raw per-monomer L2 norms (coral) sit near 3.7, while the rescaled norms (teal) concentrate around $\sqrt{384} \approx 19.6$, matching the expected norm of a 384-dimensional $\mathcal{N}(0,1)$ vector (dashed line). Fig. S6b reports the pairwise cosine similarity over 20,000 randomly sampled monomer pairs, which is invariant under positive scalar rescaling and therefore characterizes the intrinsic angular structure inherited from ChemBERTa. The distribution is centered near 0.73 with a standard deviation of 0.10, indicating a uniformly positive baseline similarity among monomers rather than an isotropic spread around zero. This positive offset is a well-known property of contextual language-model embeddings and does not, on its own, impede downstream learning, since what matters for the content, connection, and prediction models is the relative geometry among monomers rather than the absolute mean.

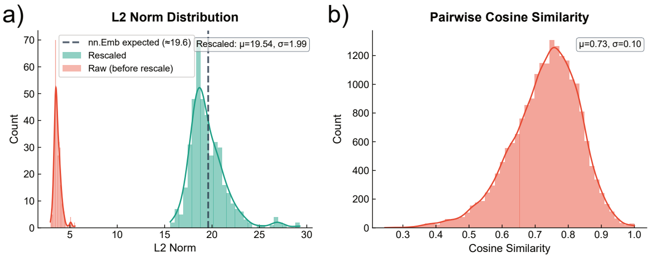

**Fig. S6: L2-norm rescaling of the ChemBERTa monomer embeddings.** **(a)** Per-monomer L2-norm distribution across the 425 monomers. Coral: raw ChemBERTa embeddings (standard deviation $\approx 0.19$). Teal: embeddings after dividing the raw tensor by its scalar standard deviation, which brings the norm distribution onto the same scale as PyTorch's default `nn.Embedding` initializer $\mathcal{N}(0,1)$. The dashed line marks the expected L2 norm of a 384-dimensional $\mathcal{N}(0,1)$ vector, $\sqrt{384} \approx 19.6$. **(b)** Pairwise cosine similarity distribution over 20,000 randomly sampled monomer pairs. Cosine is invariant under positive scalar rescaling, so this panel reports the intrinsic angular structure of the ChemBERTa embedding. The distribution is centered near 0.73 with a standard deviation of 0.10, reflecting the uniformly positive baseline similarity characteristic of contextual language-model embeddings.

### S3 Generation Model Evaluation

The three generation modules share a single `BaseTrainer` class and a common training infrastructure, and every run reported in this section was launched through the unified pipeline wrapper `Pipelines/Train_Generation.py`, which sweeps architecture and size combinations through train, evaluation, and inference-test in one call. Default architecture dimensions and training hyperparameters are defined in `Scripts/Generation_Model/Configs/arch_presets.py` and `params.py`, and can be overridden by CLI flags or a JSON config file passed to the wrapper. Table S3 lists the settings held fixed across all three modules. Architecture hyperparameters and the few module-specific knobs (label smoothing, exponential moving average (EMA) decay) are reported in the subsections below. All metrics in this section are computed on the held-out test split of the data splits constructed in Section S1.3.

The early-stopping criterion is patience-based with strict-improvement reset. The per-stage validation metric is `val_loss` for *Layout* and *Content*, and `exist_mcc`, the Matthews correlation coefficient (MCC) on the EDGE EXIST GATE, for *Connection*. The GRAPH GATE pre-filters whether the EDGE EXIST GATE runs, and the EDGE TYPE GATE fires only when an edge exists. Structural fidelity therefore reduces to `exist_mcc`. Training stochasticity (model initialisation, DataLoader shuffling, *Connection*'s dynamic edge sampling) was not seeded in the runs reported here. The released codebase configures global seeds (Python, NumPy, PyTorch CPU/CUDA RNGs, default seed 42) at the entry of every training script.

**Table S3: Default training settings shared across the three generation modules.**

| Setting | Value[a] |
|---|---|
| optimizer | AdamW[b] |
| LR schedule | cosine annealing, $\eta_{\min} = 1 \times 10^{-6}$, warmup 5% of epoch budget |
| gradient clipping | $\|\nabla\|_2 \leq 1.0$ |
| mixed precision | off |
| early stopping | per-stage validation metric, patience 20 epochs, strict-improvement criterion |
| epoch budget | 200 |
| EMA decay | 0.99[e] |

[a] Defaults applied to every run unless overridden in the per-module tables of Sections S3.1, S3.2, and S3.3.
[b] PyTorch AdamW defaults: $\beta_1 = 0.9$, $\beta_2 = 0.999$, weight decay $= 0.01$.
[e] Both raw and EMA weights are stored. Loaders default to EMA.

#### S3.1 *Layout* Stage

The *Layout* stage operates on a small vocabulary of block-type markers (`<PEP>`, `<CHEM>`) and length tokens (`<LEN_N>`) consumed as alternating pairs. To check whether this task requires architectural effort at all, we train three single-layer autoregressive backbones (GPT[16], GRU[17], LSTM[18]) with next-token cross-entropy over the block-type-and-length token sequence, on top of the common training infrastructure of Table S3. The architecture hyperparameters and Layout-specific training settings are summarized in Table S4, and all three runs complete the full 200-epoch budget without triggering the stop criterion. Generation panels are based on unconditional samples drawn at the default decoding temperature.

**Table S4: *Layout*-stage architecture and module-specific training settings.**

| | Architecture | | | | | | Training (module-specific) | | | |
|---|---|---|---|---|---|---|---|---|---|---|
| **Config** | `d_model` | `num_layers` | `n_heads` | `dim_ff` | `dropout` | `max_len` | batch | lr | label smoothing | EMA decay[a] |
| GPT | 64 | 1 | 2 | 128 | 0.1 | 20 | 40,960 | $1 \times 10^{-3}$ | 0.0 | 0.0 |
| GRU | 64 | 1 | n/a | n/a | 0.1 | 20 | 40,960 | $1 \times 10^{-3}$ | 0.0 | 0.0 |
| LSTM | 64 | 1 | n/a | n/a | 0.1 | 20 | 40,960 | $1 \times 10^{-3}$ | 0.0 | 0.0 |

[a] Value 0.0 disables EMA tracking.

Fig. S7a plots validation loss per epoch. The three curves collapse onto a shared asymptote within roughly 25 epochs and then drift only by milli-units, ending at a held-out test cross-entropy of 0.8055 (GPT), 0.8062 (GRU), and 0.8073 (LSTM). GPT attains both the lowest test loss and, visually, the earliest plateau, and the GPT-to-LSTM margin on test is 0.0018 nats per token. The *Layout* vocabulary exposes too few independent decisions for architecture choice to register meaningfully at this scale.

Generation-time fidelity is equally tight. Fig. S7b overlays the PEPTIDE-block length distribution produced by each architecture on the training distribution. The three density curves are visually indistinguishable from the ground-truth envelope, and the Jensen–Shannon divergences against the training distribution sit at $8.6 \times 10^{-4}$, $8.2 \times 10^{-4}$, and $8.7 \times 10^{-4}$ for GPT, GRU, and LSTM respectively. Pattern frequencies (Fig. S7c) are recovered to within one percentage point on every segment. All four bars place 0.81 to 0.82 of their mass on PEP-only layouts, 0.11 to 0.12 on PEP-1CHEM, 0.06 to 0.07 on PEP-2CHEM, and the remainder in the Other bin. CHEM blocks in the HELM library are length-one anchors by construction, so the per-block length axis reduces to the PEPTIDE length shown in panel (b) and is not plotted separately.

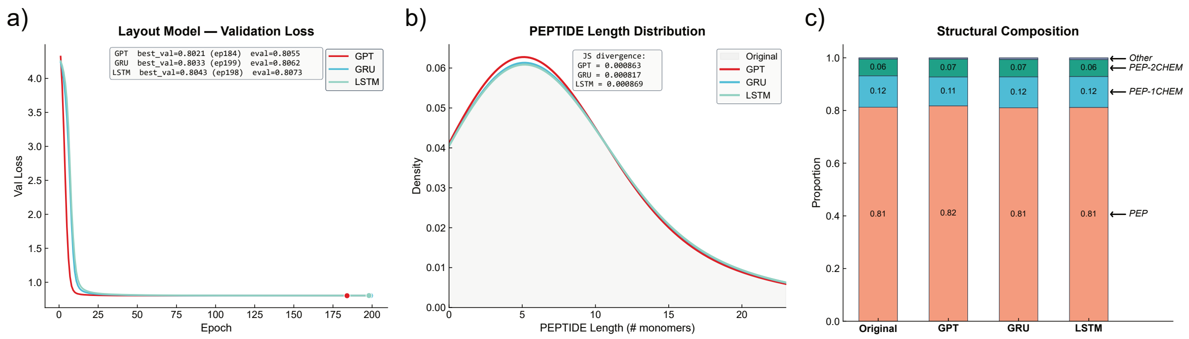

**Fig. S7: *Layout*-stage evaluation. (a)** Validation loss per epoch for the three *Layout* architectures (GPT, GRU, LSTM), trained with matched optimization settings over a 200-epoch budget. Filled dots mark the best validation epoch. The text box reports best-val loss, its epoch index, and held-out test-set loss for each architecture. **(b)** PEPTIDE-block length distribution. The grey shaded area is the training distribution and the colored curves are samples from the trained *Layout* models. Kernel density estimates use bandwidth 0.8 and are truncated at the 95th percentile of all observations. The inset reports Jensen–Shannon divergence between each generated distribution and the training distribution, computed on integer-valued histograms over the full length range. **(c)** Frequency of block-level patterns. Labels denote block sequences along the peptide: PEP for a single PEPTIDE block, PEP-1CHEM and PEP-2CHEM for a PEPTIDE block followed by one or two CHEM anchors. The three most common patterns in the training distribution are labeled and the residual mass is pooled into the Other segment.

We therefore carry the GPT configuration forward into the downstream LCC cascade on the grounds that it attains both the lowest test loss and the earliest convergence, while recognizing that the three architectures are effectively interchangeable at this scale. The broader takeaway is that the *Layout* module is a solved component in the small single-layer regime, which pushes the quality bottleneck of the full

LCC cascade onto the *Content* and *Connection* modules in the next two subsections.

#### S3.2 *Content* Stage

The *Content* stage ingests the block-type and length context produced by the *Layout* stage and fills in a monomer sequence of the specified length for each block. The four backbones span two training regimes. GPT, GRU, and LSTM are autoregressive, with each monomer predicted from the prior blocks and the intra-block prefix, and are trained with next-token cross-entropy over the full monomer sequence. BERT[19] is a masked language modeling (MLM) encoder, with masked monomer positions predicted from bidirectional context over all other blocks and the surrounding intra-block monomers, and is trained with cross-entropy over the masked positions only. Each architecture is instantiated at three sizes (small, medium, large), giving 12 configurations. All 12 share the training infrastructure of Table S3, and architecture dimensions together with module-specific training settings appear in Table S5. BERT additionally samples `mlm_probability` uniformly from $[0.15, 0.85]$ per batch, spanning gentle infilling at the low end to nearly *de novo* generation at the high end.

**Table S5: *Content*-stage architecture and module-specific training settings.**

| | Architecture | | | | | | Training (module-specific) | | | |
|---|---|---|---|---|---|---|---|---|---|---|
| **Config** | `d_model` | `num_layers` | `n_heads` | `dim_ff` | `dropout` | `max_len` | batch | lr | weight decay | label smoothing |
| GPT-small | 256 | 4 | 4 | 512 | 0.25 | 256 | 1,024 | $1\times10^{-4}$ | 0.05 | 0.25 |
| GPT-medium | 512 | 8 | 8 | 1024 | 0.35 | 512 | 384 | $1\times10^{-4}$ | 0.05 | 0.25 |
| GPT-large | 768 | 12 | 12 | 2048 | 0.40 | 512 | 128 | $8\times10^{-5}$ | 0.05 | 0.25 |
| BERT-small[a] | 256 | 4 | 4 | 512 | 0.25 | 256 | 960 | $1\times10^{-4}$ | 0.06 | 0.30 |
| BERT-medium[a] | 512 | 8 | 8 | 1024 | 0.35 | 512 | 384 | $1\times10^{-4}$ | 0.06 | 0.30 |
| BERT-large[a] | 768 | 12 | 12 | 2048 | 0.40 | 512 | 64 | $8\times10^{-5}$ | 0.06 | 0.30 |
| LSTM-small | 256 | 2 | n/a | n/a | 0.25 | 256 | 2,048 | $1\times10^{-4}$ | 0.05 | 0.25 |
| LSTM-medium | 512 | 3 | n/a | n/a | 0.30 | 512 | 1,024 | $1\times10^{-4}$ | 0.05 | 0.25 |
| LSTM-large | 768 | 4 | n/a | n/a | 0.35 | 512 | 512 | $8\times10^{-5}$ | 0.05 | 0.25 |
| GRU-small | 256 | 2 | n/a | n/a | 0.20 | 256 | 3,072 | $1\times10^{-4}$ | 0.03 | 0.20 |
| GRU-medium | 512 | 3 | n/a | n/a | 0.25 | 512 | 960 | $1\times10^{-4}$ | 0.03 | 0.20 |
| GRU-large | 768 | 4 | n/a | n/a | 0.30 | 512 | 512 | $8\times10^{-5}$ | 0.03 | 0.20 |

[a] BERT is trained with masked language modeling rather than next-token prediction. During training, `mlm_probability` is sampled uniformly from $[0.15, 0.85]$ per batch, spanning infilling at the low end to nearly *de novo* generation at the high end.

Fig. S8a plots per-epoch validation loss at each size, and Fig. S8b summarizes held-out test performance across architectures and sizes. Among the autoregressive backbones, the cross-entropy ranking GPT $<$ GRU $<$ LSTM holds at every scale, with GPT-large attaining the overall minimum (1.889, 6.61 perplexity). Convergence behaviour separates the scales cleanly. At small and medium sizes, all four architectures consume the full 200-epoch budget. At large, the autoregressive backbones plateau early (best-validation epoch 70 for LSTM, 83 for GPT, 84 for GRU), while BERT continues to improve until epoch 166. BERT's test cross-entropy sits about 0.4 nats per token above GPT, but the two losses are scored over different position sets (BERT averages over masked positions only, the autoregressive backbones over every target position) and are not directly comparable.

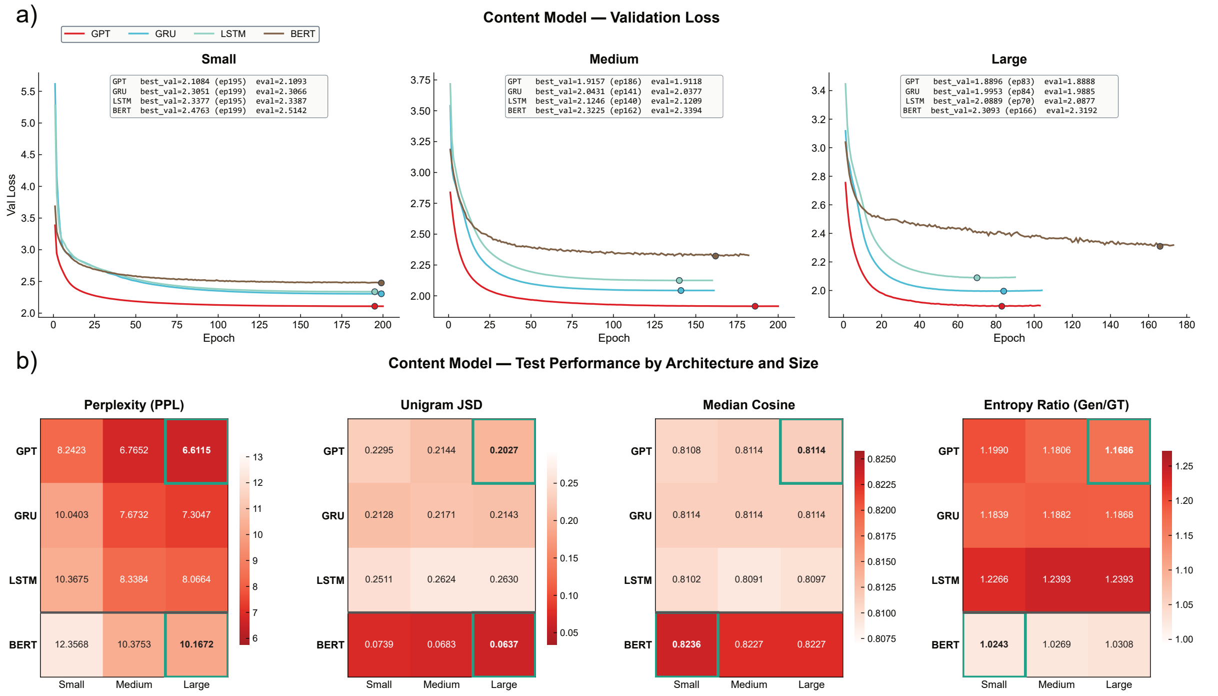

**Fig. S8: *Content*-stage convergence and held-out test performance.** **(a)** Validation loss per epoch for the 12 configurations, grouped by size (small, medium, large). Filled dots mark the best validation epoch. Text boxes report best-validation loss with its epoch index and held-out test loss for each architecture at that size. **(b)** Held-out test performance across architectures and sizes. Perplexity and unigram Jensen–Shannon divergence are lower-is-better, and the Reds colormaps are inverted so that darker saturation marks the better value. Median cosine is the embedding similarity between generated and reference monomers. Entropy ratio is generated-over-ground-truth unigram entropy, with a target of 1. Green borders mark the best cell within each architecture (GPT/GRU/LSTM compared to each other, BERT compared to itself across sizes).

Distribution-matching metrics reverse the perplexity ordering. On unigram Jensen–Shannon divergence between generated and training monomer distributions (Fig. S8b), BERT sits at 0.064–0.074 across sizes and the three autoregressive backbones at 0.20–0.26. Fig. S9a cross-checks this at the level of per-monomer log-frequency: the Spearman correlation between generated and ground-truth frequency at the large size is 0.821 for BERT, 0.777 for GPT, 0.744 for GRU, and 0.721 for LSTM. BERT's scatter lies on the diagonal across four decades, whereas the autoregressive scatters push low-frequency monomers above the diagonal. The entropy ratio (generated-over-ground-truth) is near unity for BERT (1.03) and above unity for the autoregressive backbones (1.17–1.24), the same over-dispersion signal seen in the frequency scatter. Fig. S9b narrows the lens to substitution events. At positions where the generated monomer differs from the reference, the median embedding cosine is 0.810 for the autoregressive backbones and 0.823 for BERT, both well above the random-pair baseline of 0.73. The two regimes differ mainly in substitution rate: 95–96% of positions for the autoregressive backbones and 52–57% for BERT.

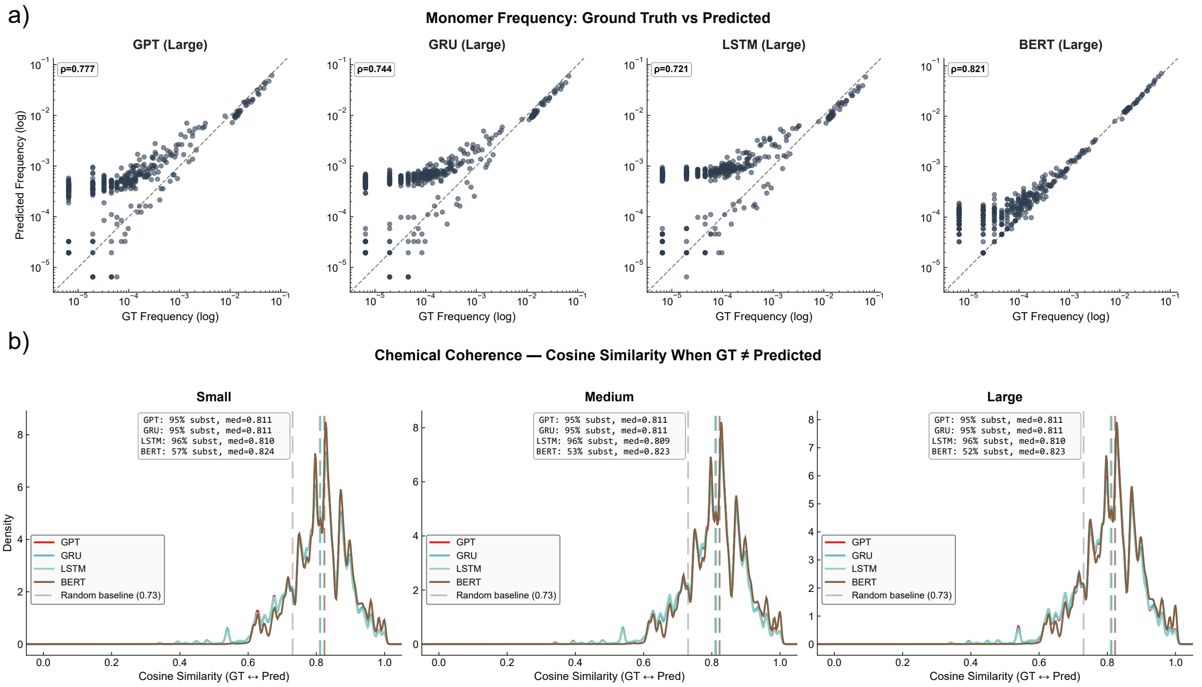

**Fig. S9:** ***Content*-stage monomer-level fidelity at the best size per architecture.** **(a)** Predicted versus ground-truth monomer frequency on log–log axes for each architecture at its best size (all four peak at the large configuration). The dashed line is the $y = x$ diagonal, and the inset reports Spearman's $\rho$ between generated and reference log-frequency. **(b)** Kernel density of the embedding cosine similarity between the generated and reference monomer at each position where the two differ, separately for small, medium, and large. The inset reports the per-architecture substitution rate (fraction of positions with generated $\neq$ reference) and the median substitution cosine. The grey dashed line at 0.73 marks the mean cosine over random monomer pairs from the library and serves as a chance-level reference.

A PEPTIDE-versus-CHEM breakdown sharpens the architectural contrast. All four backbones recover the full PEPTIDE monomer vocabulary, but CHEM vocabulary coverage rises from 0.79 (GPT) through 0.85 (GRU) and 0.96 (LSTM) to 1.00 (BERT) (Fig. S10b). On PEPTIDE positions, the median substitution cosine is tight at 0.81–0.82 across all four backbones. On CHEM positions, the autoregressive backbones drop to 0.54. Every CHEM block in the HELM library is length one, so a wrong prediction replaces the entire chemical anchor rather than one residue inside a longer run, and the resulting embedding distance is correspondingly larger. BERT has no CHEM substitution cosine to report because it predicts every test CHEM anchor exactly (28/28 at each size), so the "cosine when predicted $\neq$ reference" statistic is undefined. At the sequence level (Fig. S10a), Shannon entropy over PEPTIDE monomers is 5.45 bits in the reference, 5.63 bits for BERT, and 6.44–6.86 bits for the autoregressive backbones. Mean normalized Levenshtein distance between generated PEPTIDE sequences is 0.96–0.97 for every architecture, so pairwise sequence-identity diversity is effectively architecture-insensitive.

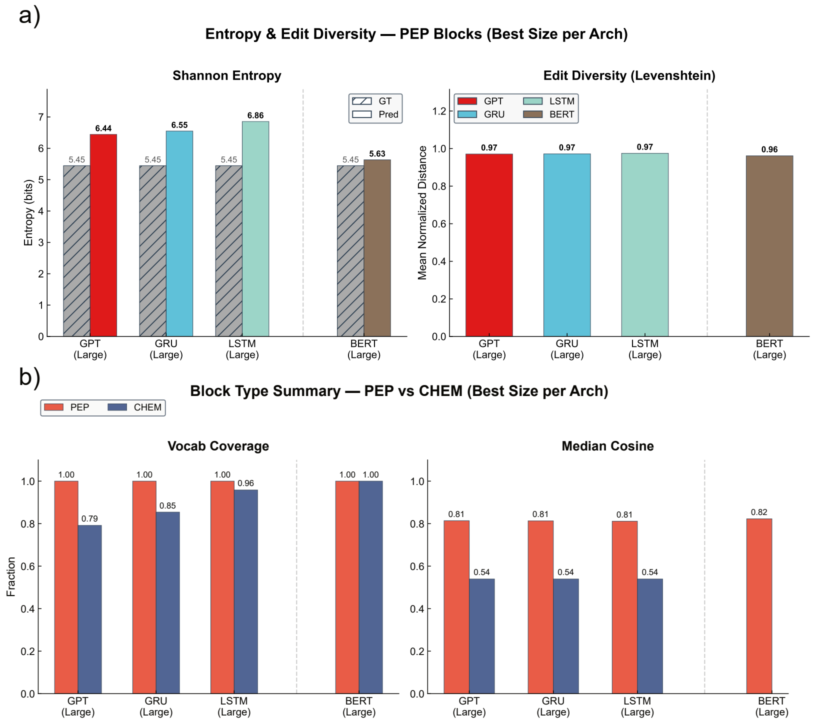

**Fig. S10: *Content*-stage diversity and block-type breakdown at the best size per architecture.** **(a)** Per-architecture sequence-level diversity on PEPTIDE blocks. Left: Shannon entropy over the monomer distribution aggregated across generated sequences (hatched grey) against the ground-truth reference (colored). Right: mean normalized Levenshtein distance between pairs of generated PEPTIDE sequences, reported as a single Pred bar per architecture. **(b)** Vocabulary coverage and median embedding cosine, split by block type (PEP versus CHEM). Vocabulary coverage is $|\text{pred} \cap \text{ground truth}|/|\text{ground truth}|$. The CHEM median cosine is omitted for BERT because BERT predicts every test CHEM anchor exactly and the "cosine when predicted $\neq$ reference" statistic is therefore undefined.

Two operating points are worth carrying forward. GPT-large attains the lowest test perplexity and the cleanest autoregressive convergence, and its broader unigram spread (JSD 0.20, entropy ratio 1.17) is the expected profile of a temperature-sampled *de novo* generator. We therefore adopt it as the default Content backbone for unconditional LCC cascade inference. BERT-large attains the tightest fit to the training monomer distribution (JSD 0.06, entropy ratio 1.03), exact recall of CHEM anchors, and parallel masked decoding, which together make it the natural choice for user-specified partial designs where only selected positions are to be re-sampled. Both configurations are registered as default Content models for downstream use, and the LCC cascade quality question shifts to the *Connection* module in the next subsection.

#### S3.3 *Connection* Stage

The *Connection* stage operates on a fully connected token graph over the monomers of a generated HELM[1] peptide and predicts special (non-backbone) bonds through three heads: a GRAPH GATE that decides whether any special connection exists, an EDGE EXIST GATE that scores each candidate edge, and a 9-class EDGE TYPE GATE covering the eight SMARTS-based linker patterns of Section S1.2 together with the head-to-tail ring rule. Five graph neural network (GNN) backbones are compared (GAT[20], GCN[21], GIN[22], MPNN[23], and Graph Transformer[24]), each instantiated at three sizes (small, medium, large) for 15 configurations total. All 15 share the training infrastructure of Table S3. Architecture dimensions and module-specific training settings appear in Table S6.

The three heads are trained jointly with a fixed-weight multi-task objective:

$$\mathscr{L}_{\text{conn}} = \mathscr{L}_{\text{exist}}^{\text{focal}} + \mathscr{L}_{\text{type}}^{\text{CE-w}} + 0.5\,\mathscr{L}_{\text{graph}}^{\text{BCE}}. \tag{S1}$$

$\mathscr{L}_{\text{exist}}^{\text{focal}}(p_t) = -(1-p_t)^{\gamma}\log p_t$ with $\gamma = 2$[25] is applied to every candidate edge in non-empty graphs, where $p_t$ is the sigmoid score on positive edges and one minus the sigmoid score on negatives. The focal factor down-weights easy negatives under the severe candidate-vs-positive imbalance. $\mathscr{L}_{\text{type}}^{\text{CE-w}}$ is class-frequency-weighted cross-entropy over the nine positive edge types of Section S1.2, applied to positive edges only. The per-class weights are $w_c \propto \sqrt{\sum_k n_k/n_c}$, clipped at $12\times$ the most-frequent class to address the long-tailed type distribution. $\mathscr{L}_{\text{graph}}^{\text{BCE}}$ is binary cross-entropy on all graphs and carries a 0.5 down-weight as an auxiliary head. The EDGE EXIST GATE is the primary task and drives both checkpoint selection and downstream structural fidelity (Section S3).

Two data-construction ratios complement the loss by targeting the imbalances that the gates and the edge classifier see during training. The empty-to-non-empty graph ratio is fixed at 0.5 in each batch so the GRAPH GATE sees a balanced binary signal. Within each non-empty graph, the negative-to-positive edge sampling ratio is 4:1 per positive edge. Negative candidates are drawn first from monomer-pair types observed as positive somewhere in training ("plausible-but-absent" edges) and then from random pairs, and are resampled every epoch so successive epochs train on different negatives.

**Table S6: *Connection*-stage architecture and module-specific training settings.**

| | Architecture | | | | | Training (module-specific) | |
|---|---|---|---|---|---|---|---|
| **Config** | `d_model` | `hidden` | `num_layers` | `n_heads` | `dropout` | batch | lr |
| GAT-small | 384 | 768 | 3 | 4 | 0.20 | 512 | $1\times10^{-4}$ |
| GAT-medium | 512 | 1024 | 4 | 8 | 0.25 | 256 | $5\times10^{-5}$ |
| GAT-large | 768 | 1536 | 6 | 8 | 0.30 | 128 | $5\times10^{-5}$ |
| GCN-small | 384 | 768 | 3 | n/a | 0.20 | 512 | $1\times10^{-4}$ |
| GCN-medium | 512 | 1024 | 4 | n/a | 0.25 | 256 | $5\times10^{-5}$ |
| GCN-large | 768 | 1536 | 6 | n/a | 0.30 | 128 | $5\times10^{-5}$ |
| GIN-small | 384 | 768 | 3 | n/a | 0.20 | 512 | $1\times10^{-4}$ |
| GIN-medium | 512 | 1024 | 4 | n/a | 0.25 | 256 | $5\times10^{-5}$ |
| GIN-large | 768 | 1536 | 6 | n/a | 0.30 | 128 | $5\times10^{-5}$ |
| MPNN-small | 384 | 768 | 3 | n/a | 0.20 | 512 | $1\times10^{-4}$ |
| MPNN-medium | 512 | 1024 | 4 | n/a | 0.25 | 256 | $5\times10^{-5}$ |
| MPNN-large | 768 | 1536 | 6 | n/a | 0.30 | 128 | $5\times10^{-5}$ |
| GraphTrans-small | 384 | 768 | 3 | 4 | 0.20 | 512 | $1\times10^{-4}$ |
| GraphTrans-medium | 512 | 1024 | 4 | 8 | 0.25 | 256 | $5\times10^{-5}$ |
| GraphTrans-large | 768 | 1536 | 6 | 8 | 0.30 | 128 | $5\times10^{-5}$ |

Fig. S11a plots per-epoch validation `exist_mcc` for the 15 configurations. *Connection* is trained with the multi-head joint loss of Eq. S1, but deployment quality reduces to `exist_mcc` because the auxiliary heads function as pre-filter and conditional refinement around edge existence. All 15 runs reach `exist_mcc` saturation around epoch 50 and continue to climb slowly thereafter by approximately $1\times10^{-4}$ per epoch. Fig. S11a shows the deployed best checkpoint of each configuration is at the `exist_mcc` max (epoch 94–173). Early stopping halts each run between epoch 114 (Graph Transformer-medium) and

193 (GAT-large). The held-out `exist_mcc` is tightly clustered across architectures and sizes (0.96–0.97), confirming that the multi-head joint loss reliably optimises the primary objective for every backbone.

Fig. S11b reports held-out test performance for each of the three heads at the deployed checkpoint. Model selection is anchored on the primary task (EDGE EXIST GATE F1); the two auxiliary heads are reported as a sanity check that joint training does not sacrifice them too much. On the GRAPH GATE (binary classification of whether the peptide carries any special connection), the MCC clusters tightly in 0.920–0.945, with GAT-medium on top (0.9451). On the EDGE EXIST GATE, the primary task, F1 is even tighter at 0.960–0.971, with GAT-large best (0.9714). The EDGE TYPE GATE (9-class macro-F1) shows the largest spread, from 0.851 (Graph Transformer-medium) to 0.929 (MPNN-large), and benefits most from increased capacity, with MPNN climbing from 0.890 at small to 0.929 at large and GAT from 0.873 to 0.912. Conditional on a correctly-predicted edge, the EDGE TYPE GATE is essentially perfect at every configuration (accuracy = 1.000 for all 15 configurations, not plotted), so the EDGE TYPE GATE macro-F1 spread is driven by the EDGE EXIST GATE's per-type recall rather than by any confusion between type classes. GAT-large is the primary-task winner, and stays within 0.012 on the GRAPH GATE and 0.017 on the EDGE TYPE GATE of the per-head winners on the two auxiliary heads, so it carries forward as the candidate backbone subject to the per-type and structural-validity analyses below.

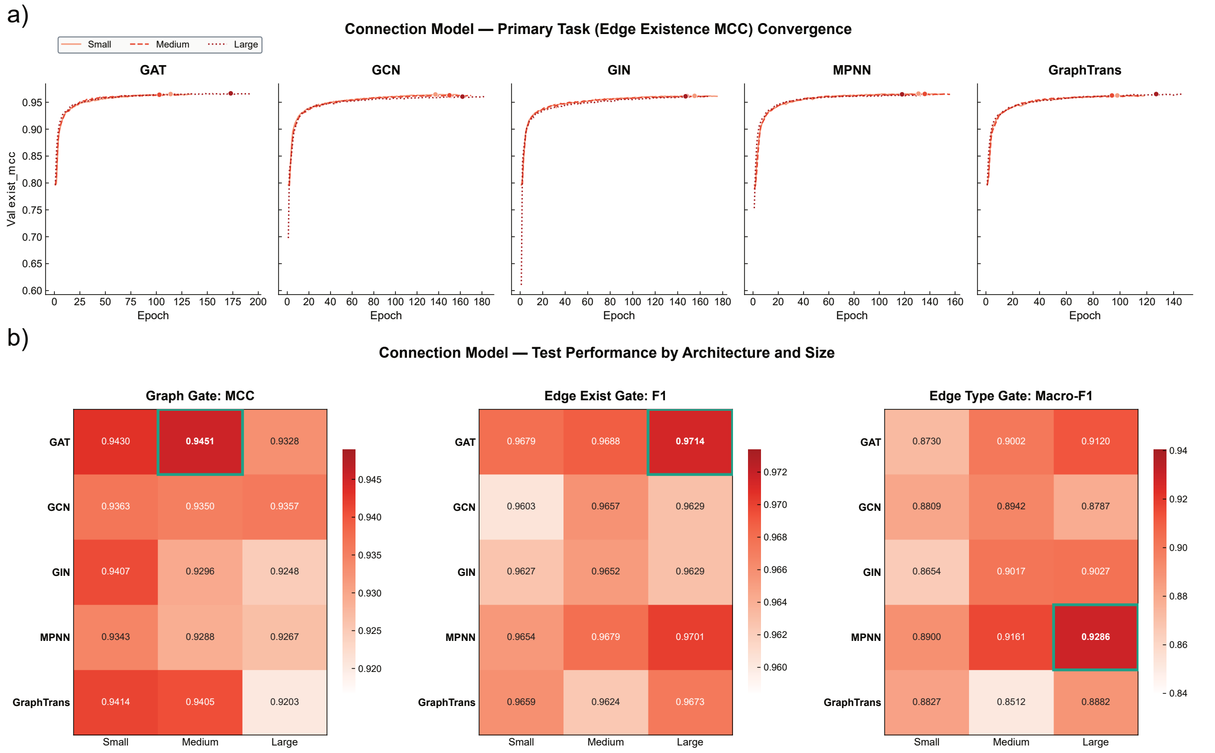

**Fig. S11: *Connection*-stage convergence and head-wise test performance.** **(a)** Validation `exist_mcc` per epoch for the 15 configurations, one panel per architecture with small, medium, and large curves overlaid. *Connection* is trained with a multi-head joint loss; we visualise the primary task's metric (edge-existence MCC; rationale in Section S3). Filled dots mark the deployed best checkpoint of each run (`exist_mcc` max). Early stopping halts each run between epoch 114 (Graph Transformer-medium) and 193 (GAT-large). **(b)** Held-out test performance for each of the three heads: MCC on the GRAPH GATE, F1 on the EDGE EXIST GATE, and macro-averaged F1 over nine edge-type classes on the EDGE TYPE GATE. All metrics are higher-is-better. Reds colormaps are used across panels and each panel has its own tight scale. Green borders mark the best cell per panel.

Fig. S12 resolves EDGE EXIST GATE performance by connection type at the best size per architecture (selected by EDGE EXIST GATE F1, the deployment criterion), to check whether each architecture's aggregate F1 is uniform across types or driven by a subset. The five common types (Amide $n = 8{,}232$, Ester $n = 1{,}366$, PG-Amide $n = 1{,}218$, Disulfide $n = 1{,}141$, H2T Ring $n = 1{,}023$) cluster tightly at $P, R \geq 0.9$ across all five architectures, and the per-architecture dumbbells overlap within the visible line width. The four rare types (Lanthi. $n = 17$, Bz_stapled $n = 10$, Sulfanil. $n = 9$, Trp-Cys $n = 3$) show wide per-architecture dispersion. Most of this spread is a support problem rather than an architecture problem, because single-digit test support provides little statistical stability: one missed Trp-Cys call out of three shifts recall from 1.0 to 0.67, and arch-to-arch differences on $n \leq 17$ types are comparable to the resolution of the metric itself. The aggregate-F1 advantage of GAT-large therefore reflects uniform high performance on common types, not an outlier on a single rare type.

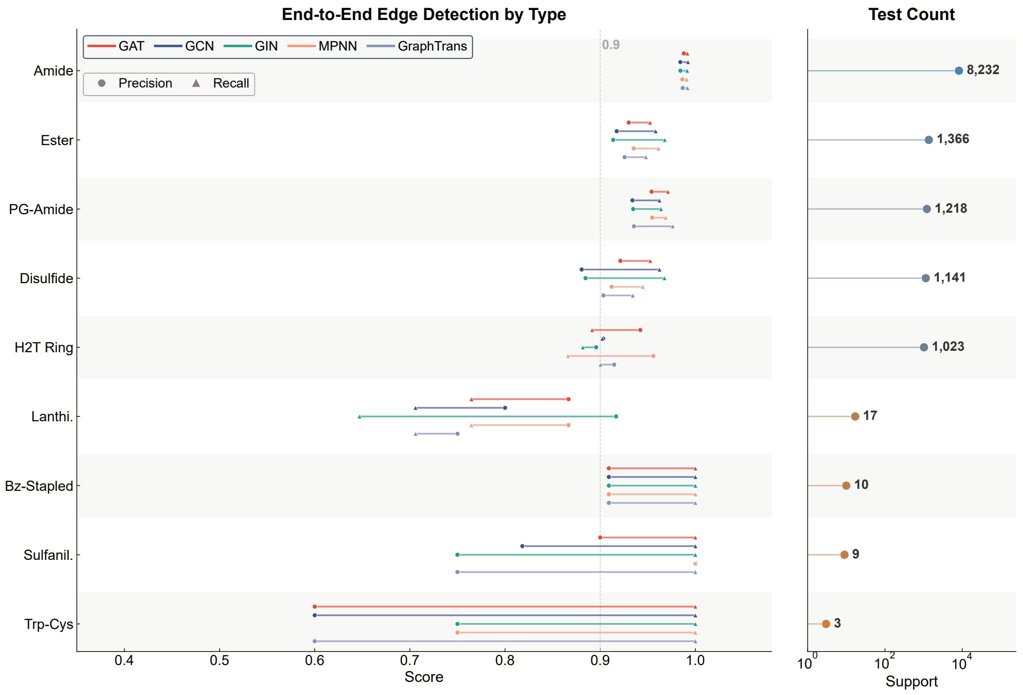

**Fig. S12: Per-type precision and recall for the *Connection* stage, best size per architecture (selected by EDGE EXIST GATE F1, the deployment criterion).** Each row is one connection type, sorted by test-set support (bars on the right, edge counts). For each architecture (colour-coded, see legend), a dumbbell connects the precision (●) and recall (▲) of the EDGE EXIST GATE on edges of that type.

Fig. S13 closes the per-architecture comparison by reporting four peptide-level error rates, each computed as a fraction of all test graphs (a single graph may contribute to more than one rate, so the four rates overlap rather than partition failures). The chemistry-invalid and missing-required rates both remain below 0.5% at every configuration, but for different reasons. The chemistry-invalid rate measures graphs whose predicted edges would violate a physical or topological constraint: an R-group atom used twice, an R-group label not on the monomer's allowed list, a bond from a monomer to itself, or a multi-block peptide left as disconnected fragments. Its near-zero value is a structural guarantee from the decoder rather than chemistry learned by the model. The greedy edge selector calls `fast_graph_validity_check` on every candidate and refuses any edge that would violate (1) R-group occupancy, (2) R-group-label compatibility, or (3) self-loop constraints, so no such edge ever reaches the output. The residual $< 0.5\%$ is almost entirely multi-block peptides whose predicted edges fail to bridge all blocks into a single connected structure, a condition tested only on the completed graph. Missing-required events capture the opposite failure: a reference peptide that demands at least one special connection but receives zero predicted edges, so a cyclic or crosslinked reference is degraded to a linear prediction. Their rarity indicates that the EDGE EXIST GATE and EDGE TYPE GATE together almost always find at least one plausible bond to propose on peptides that need one.

The two edge-count mismatch rates capture the residual error that survives decoder-level filtering, and their interpretation is geometric rather than constraint-based. Over-connection (5.0–6.9%) is a chemically valid peptide with more bonds than the reference, such as a bicyclic variant of a mono-cyclic reference, or a second staple added across an already-cyclized backbone. Under-connection (4.7–5.4%) is the reverse: a chemically valid peptide with fewer bonds, such as a linear variant of a reference cycle. The near-equality of the two rates indicates that errors are distributed symmetrically around the correct bond count, without a systematic tendency to over- or under-predict, so any residual improvement would need to reduce overall edge-decision error rather than only shift the EDGE EXIST GATE threshold.

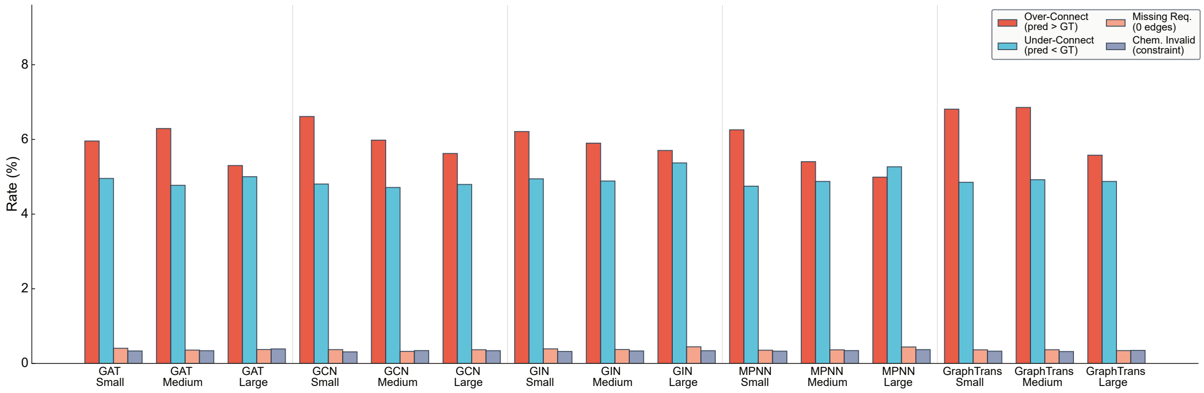

**Fig. S13: Structural validity of *Connection*-stage predictions, all 15 configurations.** Four chemistry-aware failure rates are reported as a percentage of test graphs. *Over-Connect* and *Under-Connect* are mismatches in predicted edge count relative to ground truth. *Missing Required* is the subset where ground truth has at least one edge but the model predicted none. *Chem. Invalid* is the subset where predicted edges violate an R-group, self-loop, or block-connectivity constraint. All 15 configurations cluster inside a $\pm 1\%$ band on every rate, indicating that aggregate structural validity is architecture-insensitive. Per-type precision and recall vary across architectures, especially on rare types (Fig. S12).

All 15 configurations cluster inside a narrow band on every aggregate rate, so architecture choice does not move the aggregate structural-validity dial. We adopt GAT-large as the default *Connection* backbone on three grounds: (1) it attains the highest EDGE EXIST GATE F1 (0.9714), the primary task whose MCC drives both checkpoint selection and structural fidelity (Section S3); (2) on the auxiliary heads its performance stays within 0.012 (GRAPH GATE vs GAT-medium, 0.9451) and 0.017 (EDGE TYPE GATE vs MPNN-large, 0.9286) of the per-head winners, so the primary-task choice does not compromise auxiliary quality; (3) structural-validity rates are architecture-insensitive, so the choice carries no downstream penalty. This closes the three-stage generator: *Layout* uses GPT, *Content* defaults to GPT-large with BERT-large available for user-specified infilling, and *Connection* uses GAT-large.

#### S3.4 End-to-End Monomer Vocabulary Coverage

The three preceding subsections decompose the LCC cascade into its stages. We close Section S3 with an LCC-cascade-level measurement read off at the output boundary. Fig. 3a, b established that the LCC cascade produces more novel peptides than Flat GPT by MAP4C[26, 27] nearest-neighbor distance, but that metric operates on whole-molecule fingerprints and leaves open whether the novelty comes from recombining familiar monomers into new arrangements or from drawing on a broader monomer vocabulary. We resolve this by enumerating the set of monomers each model produces (parsing each HELM string with the regex `(?:PEPTIDE|CHEM)\d+\{([^}]+)\}` and taking the union over all five runs on validity-passing peptides) and comparing against the 386-monomer training vocabulary collected the same way from the 383,817 training HELM sequences.

In *de novo* generation (Fig. S14a, 100K samples per run, 5 runs), the LCC cascade covers 385 of the 386 training monomers (99.7%) and additionally produces 21 monomers that do not appear in any training sequence, whereas Flat GPT covers only 300 training monomers (77.7%) and produces 1 out-of-training monomer. The 85-monomer gap is the long tail of low-frequency monomers that an end-to-end decoder assigns near-zero probability at the 100K scale, while the factorized Content decoder samples position-by-position from the full monomer alphabet and therefore retains non-trivial mass on rare building blocks. We attribute this to factorization rather than to two unrelated differences between the arms: Flat GPT uses nucleus sampling and a GPT-medium backbone, while the LCC *Content* stage uses top-$k$ sampling and a GPT-Large backbone (full configurations in Section S7.1). Neither difference can close the 85-monomer gap on its own, because both only reweight monomers the model already assigns non-trivial probability. Loosening the nucleus cutoff or scaling up the backbone cannot rescue a monomer whose Flat-GPT joint probability has been multiplied down to near-zero across the full sequence. Lifting such a monomer from "never sampled" to "sampled at least once" over 100K draws requires a structural change to the decoder, not a different sampler or a wider Transformer.

In the infilling setting (Fig. S14b, reveal ratio 0.5 on the same test peptides used for Fig. 3g–i, 5 runs), both architectures cover nearly all training monomers (GPT $368/386 = 95.3\%$, BERT $370/386 = 95.9\%$) and produce a small, nearly equal number of out-of-training monomers (GPT 9, BERT 10). The near-parity follows from the protocol: half the monomer positions are revealed from the test peptide, pinning a large fraction of the output vocabulary to already-seen tokens regardless of the filling architecture. At this reveal ratio the vocabulary-level footprint of BERT and GPT is therefore indistinguishable, and the two models differ only in how they fill the free positions.

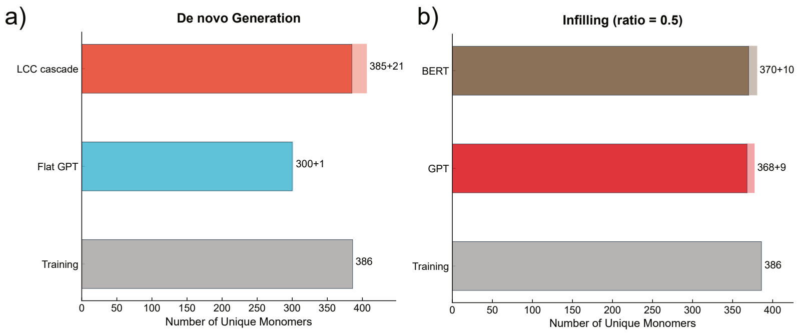

**Fig. S14: Monomer vocabulary coverage (a)** *De novo* generation (100K samples per run, 5 runs, union over runs). Each model bar has a solid segment (monomers shared with the 386-monomer training vocabulary) and a hatched segment (monomers not present in any training sequence), with the numeric label `shared+novel` at the bar end. **(b)** Infilling on the held-out test set (reveal ratio 0.5, 5 runs, union over runs), same segmentation applied to GPT and BERT as Content backbones.

### S4 Constrained Generation Scenarios

The inference pipeline is driven by `Pipelines/Inference.py`, which instantiates `InferencePipeline` (defined in `Scripts/Inference/pipeline.py`) and invokes its `.run()` method. It exposes three structural constraint axes plus one post-hoc filter, all configured through the JSON request:

1. `input.layout_tokens` or `constraints.layout_constraints` fixes the *Layout*-stage block-type and length prefix (exact sequence) or admissible range (block types and lengths), respectively.
2. `constraints.content_constraints` fixes specific monomers at specific 1-indexed positions inside each polymer block (HELM[1] convention; the first monomer is position 1), injected into the *Content* stage as a positional prefix during decoding.
3. `constraints.forced_connections` injects a specific edge into the *Connection* decoder via three field groups, `from_polymer+from_pos`, `to_polymer+to_pos`, and `connection_type`, bypassing the EDGE EXIST GATE for that edge. Positions use the same 1-indexed convention.
4. `constraints.filter_connection_type` applies post-hoc filtering to the generated pool, retaining only peptides whose EDGE TYPE GATE realizes the requested type(s).

Each run draws raw samples until either `params.num_samples` = 1000 accepted peptides are collected or `raw_budget` = 100,000 pipeline attempts are exhausted, whichever comes first. Table S7 summarizes nine scenarios chosen to sweep constraint-specification strategies, with particular attention to how each strategy interacts with rare connection types.

**Table S7: Constrained-generation summary across nine scenarios.**

| Scenario | Strategy | Description | Attempts | Validity[a] (%) | Yield[b] (%) |
|---|---|---|---|---|---|
| CG1 | L | PEP×7 | 1,119 | 89.4 | 89.4 |
| CG5 | L (multi-block) | PEP×9 + CHEM×1 | 1,241 | 80.6 | 80.6 |
| CG8 | L (range) | PEP len ∈ [5,10], free block types | 1,158 | 86.4 | 86.4 |
| CG9 | L (range, PEP-only) | PEP-only, len ∈ [5,12] | 1,121 | 89.2 | 89.2 |
| CG2 | L+C | PEP×10; pos3=C, pos8=C | 1,139 | 87.8 | 87.8 |
| CG3 | L+C+Conn | PEP×9; pos2=C, pos8=C; Disulfide(2,8) | 1,417 | 90.4 | 70.6 |
| CG4 | L+C+Conn (rare) | PEP×7; pos1=C, pos5=dA; Lanthionin(1,5) | 100,000 | 76.7 | 0.158 |
| CG6 | L+Conn (rare, C free) | PEP×10 + CHEM×1; Sulfanilamide(PEP:5, CHEM:1) | 100,000 | 100.0 | 0.004 |
| CG7 | L+Filter (rare) | PEP×12; filter: Trp-Cys & head2tail ring | 100,000 | 85.5 | 0.009 |

[a] Fraction of generated peptides passing HELM → SMILES → HELM roundtrip.

[b] Fraction of total pipeline attempts producing an accepted peptide (satisfies constraints AND passes roundtrip). Runs terminate at `target` = 1000 accepted OR `raw_budget` = 100,000 attempts.

The six scenarios that combine either a single-axis Layout constraint (CG1, CG5, CG8, CG9) or the full Layout+Content+Connection specification over a common connection (CG2, CG3) reach the 1000-accepted target within 1,100–1,420 attempts, giving Yield values between 70.6% and 89.4%. Validity tracks Yield closely in this regime because generation-stage rejection is rare. The residual gap comes from a small fraction of generated HELM strings that fail the HELM → SMILES → HELM roundtrip. CG3, which forces a Disulfide between two Cys at fixed positions 2 and 8, loses additional attempts at the generation stage (1000 accepted from 1,417 attempts) because the *Content* stage occasionally samples a sequence that conflicts with the forced edge, but Validity remains at 90.4%.

The three rare-connection scenarios (CG4, CG6, CG7) all exhaust the 100,000-attempt budget without reaching the 1000-accepted target, and their failure distributions differ by more than an order of magnitude in ways that map directly onto constraint-specification strategy. CG4 (L+C+Conn, Lanthionin between Cys at position 1 and dA at position 5) fully specifies the Layout, Content, and Connection axes. Two hundred and six of 100,000 attempts survive the generation stage, of which 158 pass roundtrip and satisfy the forced edge, giving Yield $= 0.158\%$. The generation-stage attrition reflects the joint requirement that the *Content* stage samples a sequence compatible with the forced Lanthionin endpoint constraints and R-group compatibility; once through, Validity stays at $76.7\%$. CG6 (L+Conn, Sulfanilamide between `PEPTIDE:5` and `CHEM:1`, content free) forces the edge but leaves Content unconstrained. Only 4 of 100,000 attempts survive the generation stage because the *Content* stage samples residues at the two endpoint positions without knowledge of the Sulfanilamide R-group compatibility, so the edge-injection check rejects almost every sample. Yield collapses to 0.004%, a factor of $\sim$40 below CG4. CG7 (L+Filter, filter = "Trp-Cys $\wedge$ head2tail ring") uses no structural constraint beyond Layout and applies a post-hoc filter. Generation-stage acceptance is 100% and Validity reflects roundtrip only (85.5%), but the conjunctive filter over two rare patterns (Trp-Cys training support $n = 3$, head2tail ring support $n = 1023$) matches only 9 of 100,000 samples, giving Yield $= 0.009\%$.

The CG4 $\rightarrow$ CG6 $\rightarrow$ CG7 progression quantifies the cost of loosening specification for rare types. Moving from full specification (CG4) to connection-only injection with free content (CG6) lowers Yield by $\sim$40$\times$ because the *Content* stage samples endpoint residues blind to the connection's R-group requirements, triggering an edge-injection rejection on almost every sample. Moving from full specification (CG4) to post-hoc filtering with free generation (CG7) lowers Yield by $\sim$18$\times$ because the unconstrained generation distribution concentrates on common motifs rather than the requested rare pattern. Among the three rare-type strategies tested, combined Layout+Content+Connection specification (CG4) yields 18–40$\times$ more accepted peptides per attempt than connection-only injection (CG6) or filter-only post-processing (CG7), and is therefore the most efficient strategy within a fixed compute budget. Users targeting rare-type peptides through the web interface should fix the endpoint residues alongside the forced edge, rather than rely on connection-only injection or filter-only post-processing.

### S5 Antimicrobial Potency Prediction Model Comparison

The prediction model is a drop-in scorer invoked after the LCC cascade (Fig. 2e) and is trained once on fixed supervised data, independently of the generation modules of Section S3. The 48 predictors compared in this section share a single `BaseTrainer` class and a common training infrastructure, and every run was launched through the unified pipeline wrapper `Pipelines/Train_Prediction.py`, which sweeps architecture, size, and encoding combinations through train and evaluation in one call. Default architecture dimensions and training hyperparameters are defined in `Scripts/Prediction_Model/Configs/arch_presets.py` and `params.py`, and can be overridden by CLI flags or a JSON config file passed to the wrapper. Table S8 lists the settings held fixed across all 48 runs. Architecture hyperparameters and the few per-architecture overrides appear in the unified Table S9 of Section S5.1. All metrics in this section are computed on the held-out test split of Section S1.3.

Confidence intervals on the test-set metrics reported in Section S5.4 (Figs. 4a–b) were computed by paired bootstrap. Specifically, 1,000 index sets of size $n_{\text{test}} = 2{,}206$ were drawn with replacement from the test split using a fixed RNG seed of 42, and the same set of resampled indices was applied to every entry (each individual member and each ensemble strategy) so that pairwise differences (e.g., weighted vs. soft vote) operate on identical resampled subsets. For each of the 1,000 resamples we recomputed accuracy, macro-F1, weighted-F1, MCC, and per-class F1, and report the 2.5% and 97.5% percentiles of each metric's bootstrap distribution as a 95% CI. The implementation is the helper `_bootstrap_ci()` in `Scripts/Prediction_Model/Evaluation/eval_ensemble.py`. The bootstrap step is enabled by the CLI flag `--bootstrap_n 1000`, and the resulting CI fields are appended to the same `ensemble_comparison.json` report consumed by the figure-rendering code.

**Table S8: Default training settings shared across all 48 antimicrobial potency predictors.**

| Setting | Value[a] |
|---|---|
| optimizer | AdamW[b] |
| LR schedule | cosine annealing, $\eta_{\min} = 1 \times 10^{-6}$, warmup 5% of epoch budget |
| gradient clipping | $\|\|\nabla\|\|_2 \leq 1.0$ |
| mixed precision | off |
| early stopping | validation MCC, patience 15 epochs, min delta 0.001 |
| epoch budget | 200 |
| EMA decay | 0.99 (best-EMA checkpoint retained by validation MCC) |
| classification loss | inverse-frequency-weighted cross-entropy on ordinal-Gaussian soft targets[28] ($\sigma = 1.0$), with focal modulation[25] ($\gamma = 2.0$) |
| negative sampling | dynamic 1:1 unlabelled-to-active, resampled every epoch[c] |

[a] Values listed are defaults applied to every run unless explicitly overridden in the per-architecture Table S9 of Section S5.1.
[b] PyTorch AdamW defaults: $\beta_1 = 0.9$, $\beta_2 = 0.999$, weight decay = 0.01.
[c] Implemented by `UnlabeledNegativeSampler.set_epoch()`; draws 8,820 class-0 background negatives from the 47,467-peptide training pool (Section S5.1) at the start of every epoch.

#### S5.1 Architectures and Training Data

Potency is cast as an ordinal 5-class label. The four potency classes are derived from the DBAASP MIC column described in Section S1.3 with thresholds `activity_bins = [8, 32, 128]` $\mu$g/mL: class 1 (MIC $\geq$ 128, least potent), class 2 (32 $\leq$ MIC $<$ 128), class 3 (8 $\leq$ MIC $<$ 32), class 4 (MIC $<$ 8, most potent). Class 0 is the background class, defined in the construction below, and carries no DBAASP

MIC. The 11,026 potency-tagged HELM entries of Table S2 are split 80/10/10 by InChIKey into 8,820 training, 1,103 validation, and 1,103 test actives. InChIKey distinguishes stereoisomers, so L/D-isomer pairs of the same backbone may end up on opposite sides of the split, and the reported test-set MCC should be read as an upper bound under stereo-blind generalisation. The training actives distribute across classes 1–4 as 1,858/2,495/2,847/1,620, and the underlying MIC histogram is shown in Fig. S3c.

Class 0 (the background class) is a positive-unlabelled (PU) negative, not a low-potency bin below class 1: the four MIC-defined classes are the labelled positives, and class 0 supplies the negative-side training signal from peptides with no reported antimicrobial activity. The distinction from class 1 is one of label provenance, not potency: a class 1 peptide carries a measured MIC $\geq 128$ $\mu$g/mL and is an in-distribution weak active, whereas a class 0 peptide has no measured MIC at all and lies off the potency axis. Class 0 therefore acts as an out-of-distribution (OOD) detector that, at scoring time, absorbs generated peptides unlike known actives instead of forcing them onto the four-bin scale, trading recall on unreported actives for screening precision. These PU negatives are sampled from an unlabelled pool built from the 383,817-peptide generation training set of Table S2 by (1) filtering to HELM sequences of 12 to 30 monomers, the length window that matches the active set, and (2) InChIKey-deduplicating against every active peptide to prevent leakage. Val and test each receive a fixed negative sample matched 1:1 to their active count (1,103 peptides each). The remaining 47,467 unlabelled peptides form the training pool from which the dynamic sampler of Table S8 redraws an independent 1:1 negative set each epoch.

We sweep a factored design of (encoding) $\times$ (family) $\times$ (architecture) $\times$ (size). The encoding axis has two values, monomer-level HELM[1] tokens and atom-level SMILES. The family axis has two values: language models (LLM) consuming the tokenized string (GPT[16], GRU[17], LSTM[18]) and GNNs consuming a molecule graph (GAT[20], GCN[21], GIN[22], MPNN[23], Graph Transformer[24]). In HELM mode the GNN operates on a monomer-level graph (one node per monomer, edges following the backbone and the special connections of Section S1.2) while in SMILES mode it operates on an atom-level graph built via `RDKit.from_smiles` and featurized with the OGB atom and bond encoders[29]. All 8 architectures are instantiated at three sizes (small: hidden 256, medium: hidden 512, large: hidden 768), giving $8 \times 3 \times 2 = 48$ trained predictors. Architecture dimensions and per-architecture training settings appear in Table S9. All 48 configurations are trained with inverse-frequency-weighted cross-entropy on ordinal-Gaussian soft targets and focal modulation (Table S8).

**Table S9: Prediction model architecture and training settings across LLM and GNN families.**

| | | Architecture | | | | | | Training (per configuration)[a] | | |
|---|---|---|---|---|---|---|---|---|---|---|
| **Family** | **Config** | `hidden` | `num_layers` | `heads` | `dim_ff` | `edge_dim`[b] | `dropout` | batch | lr | wd |
| | GPT-small | 256 | 4 | 4 | 512 | n/a | 0.25 | 128 | $1 \times 10^{-4}$ | 0.01 |
| | GPT-medium | 512 | 8 | 8 | 1024 | n/a | 0.35 | 64 | $5 \times 10^{-5}$ | 0.01 |
| | GPT-large | 768 | 12 | 12 | 2048 | n/a | 0.40 | 32 | $5 \times 10^{-5}$ | 0.02 |
| | GRU-small | 256 | 2 | n/a | n/a | n/a | 0.25 | 128 | $1 \times 10^{-3}$ | 0.01 |
| LLM | GRU-medium | 512 | 3 | n/a | n/a | n/a | 0.30 | 64 | $5 \times 10^{-4}$ | 0.01 |
| | GRU-large | 768 | 4 | n/a | n/a | n/a | 0.35 | 32 | $2.5 \times 10^{-4}$ | 0.01 |
| | LSTM-small | 256 | 2 | n/a | n/a | n/a | 0.25 | 128 | $1 \times 10^{-3}$ | 0.01 |
| | LSTM-medium | 512 | 3 | n/a | n/a | n/a | 0.30 | 64 | $5 \times 10^{-4}$ | 0.01 |
| | LSTM-large | 768 | 4 | n/a | n/a | n/a | 0.35 | 32 | $2.5 \times 10^{-4}$ | 0.01 |
| | GAT-small | 256 | 3 | 4 | n/a | 64 | 0.25 | 128 | $1 \times 10^{-3}$ | 0.01 |
| | GAT-medium | 512 | 4 | 8 | n/a | 128 | 0.30 | 64 | $1 \times 10^{-3}$ | 0.01 |
| | GAT-large | 768 | 6 | 8 | n/a | 128 | 0.30 | 32 | $5 \times 10^{-4}$ | 0.01 |
| | GCN-small | 256 | 3 | n/a | n/a | n/a | 0.25 | 128 | $1 \times 10^{-3}$ | 0.01 |
| | GCN-medium | 512 | 4 | n/a | n/a | n/a | 0.30 | 64 | $1 \times 10^{-3}$ | 0.01 |
| | GCN-large | 768 | 6 | n/a | n/a | n/a | 0.30 | 32 | $5 \times 10^{-4}$ | 0.01 |
| | GIN-small | 256 | 3 | n/a | n/a | 64 | 0.25 | 128 | $1 \times 10^{-3}$ | 0.01 |
| GNN | GIN-medium | 512 | 4 | n/a | n/a | 128 | 0.30 | 64 | $1 \times 10^{-3}$ | 0.01 |
| | GIN-large | 768 | 6 | n/a | n/a | 128 | 0.30 | 32 | $5 \times 10^{-4}$ | 0.01 |
| | MPNN-small | 256 | 3 | n/a | n/a | 64 | 0.25 | 128 | $1 \times 10^{-3}$ | 0.01 |
| | MPNN-medium | 512 | 4 | n/a | n/a | 128 | 0.30 | 64 | $1 \times 10^{-3}$ | 0.01 |
| | MPNN-large | 768 | 6 | n/a | n/a | 128 | 0.30 | 32 | $5 \times 10^{-4}$ | 0.01 |
| | GraphTrans-small | 256 | 3 | 4 | n/a | 64 | 0.25 | 128 | $1 \times 10^{-3}$ | 0.02 |
| | GraphTrans-medium | 512 | 4 | 8 | n/a | 128 | 0.30 | 64 | $1 \times 10^{-3}$ | 0.02 |
| | GraphTrans-large | 768 | 6 | 8 | n/a | 128 | 0.30 | 32 | $5 \times 10^{-4}$ | 0.02 |

[a] "wd" is weight decay.

[b] Learned edge-feature dimensionality; unused by GCN (edge-type-indexed relational convolution).

#### S5.2 Architecture×Size×Encoding Selection

We first inspect training convergence. Fig. S15 overlays per-epoch validation loss for the 48 runs, split into a GNN panel (5 architectures × 3 sizes × 2 encodings, Fig. S15a) and an LLM panel (3 architectures × 3 sizes × 2 encodings, Fig. S15b). All 48 runs reach their best-validation-MCC checkpoint within the 200-epoch budget and trigger early stopping without divergence, but the convergence quality is uneven. GNN curves descend smoothly and monotonically yet saturate at a higher validation-loss plateau than the LLMs. LLM curves are noisier throughout training, with GPT-large in both encoding modes showing the largest epoch-to-epoch oscillation. Despite this, GPT-large is at least as good as GPT-medium on the held-out test set (MCC 0.529 vs 0.495 for SMILES, 0.499 vs 0.495 for HELM, the latter difference within bootstrap noise reported in Section S5.4), so the apparent training instability does not translate to a generalization deficit. Several HELM-encoded LLM runs, most visibly GPT-HELM, also drift upward in validation loss after their best-validation epoch, consistent with late-stage overfitting.

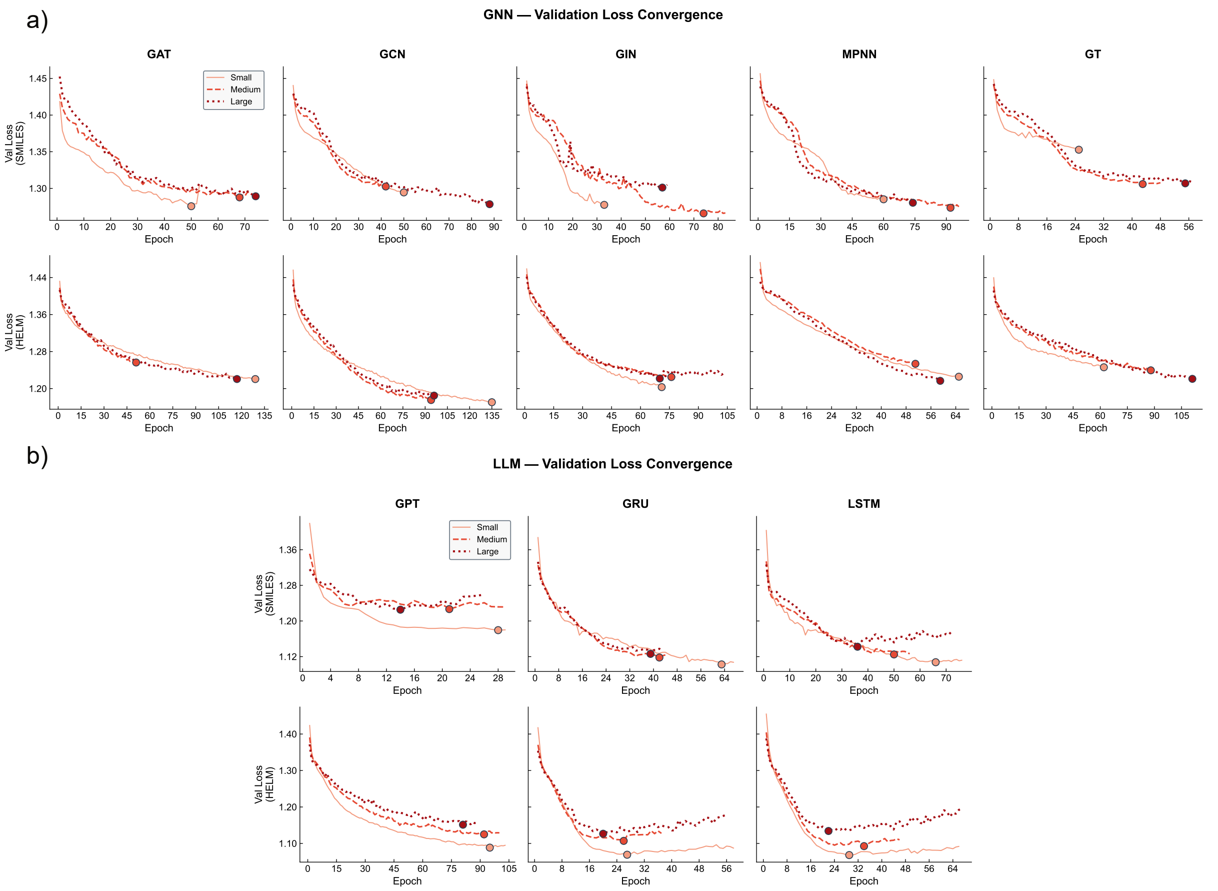

**Fig. S15: Validation loss per epoch for the 48 prediction configurations.** **(a)** GNN family: rows are encoding modes (SMILES, HELM) and columns are the five backbones (GAT, GCN, GIN, MPNN, Graph Transformer); each subplot overlays small, medium, and large sizes. **(b)** LLM family: rows are encoding modes and columns are the three backbones (GPT, GRU, LSTM); each subplot overlays the three sizes.

Fig. S16 plots four held-out test metrics (accuracy, macro-F1, weighted-F1, MCC) across the 48 configurations, arranged as 8 architectures $\times$ 6 (encoding, size) columns (SMILES-S/M/L on the left, HELM-S/M/L on the right). Performance is compressed into a narrow band: the best single configuration reaches MCC 0.587 (LSTM-large with SMILES encoding). At the family level, the best LLM configuration outperforms the best GNN configuration on MCC by 0.091 (LSTM-large-SMILES, 0.587 vs GCN-medium-HELM, 0.496), and 28 of 30 GNN configurations sit at or below the lowest-scoring LLM configuration on MCC (HELM-GPT-small, 0.476). Large is the best size in 9 of 16 (architecture, encoding) cells. The remaining 7 are best at Medium (GRU-HELM, LSTM-HELM, GCN-HELM, Graph Transformer-HELM, Graph Transformer-SMILES) or Small (GAT-SMILES, GIN-SMILES). No configuration in the sweep exceeds test MCC 0.59, motivating the ensemble construction examined in Section S5.4.

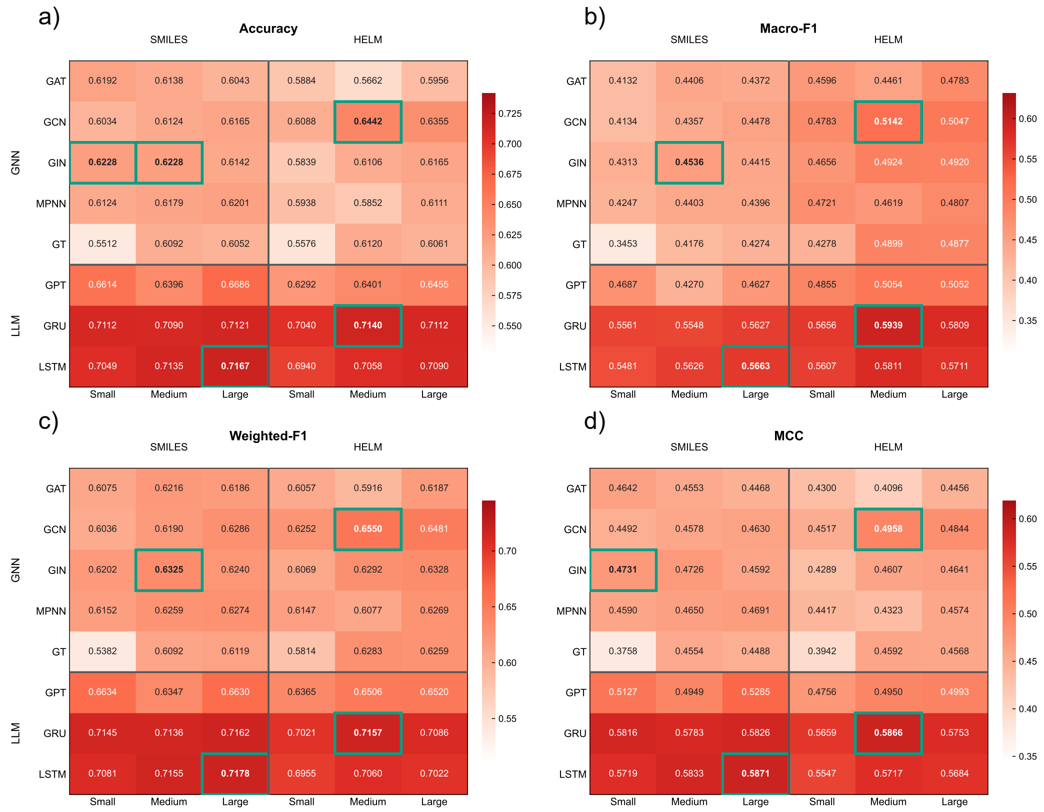

**Fig. S16: Held-out test performance of the 48 prediction configurations.** Four metrics (accuracy, macro-F1, weighted-F1, MCC) arranged as a $2 \times 2$ grid of heatmaps. Within each panel, rows are the 8 architectures (GNN family above, LLM family below, separated by a horizontal rule) and columns are the 6 (encoding, size) combinations (SMILES-S/M/L $\|$ HELM-S/M/L). Each panel has its own colour scale; within each panel, the highest-scoring cell of each (family, encoding) quadrant under that panel's metric is highlighted with a bold border (four borders per panel). These borders mark per-quadrant test-set maxima and are not the same selection as the four ensemble members of Section S5.4, which are chosen by validation MCC and therefore differ from the bordered cells in this figure.

To characterize how each of these four winners fails, Fig. S17 shows their recall-normalized confusion matrices. LSTM-large-SMILES (LLM-SMILES) classifies background cleanly (recall 0.973) but spreads off-diagonal mass broadly across the four active bins (class_1 F1 0.478, class_2 F1 0.404, class_3 F1 0.469, class_4 F1 0.504). LSTM-medium-HELM (LLM-HELM) softens background recall slightly (0.919) but is more balanced across active bins, peaking at class_4 F1 0.568. GCN-large-HELM (GNN-HELM) loses about 16% of background recall (0.844): true negatives spill primarily into class_1, a distinctive background-to-class_1 leakage less prominent in the LLM winners. GCN-large-SMILES (GNN-SMILES) has the most uneven profile: class_3 is over-called (recall 0.594) while class_4 collapses (recall 0.166), with the model systematically pulling class_4 predictions into the class_3 bin. The four patterns are partially complementary (the two LLM members conserve background precision and confuse within the active bins, the two GNN members sacrifice background purity but redistribute active-bin errors differently from the LLMs), which is the structural reason to combine them in Section S5.4 rather than deploy the best single model.

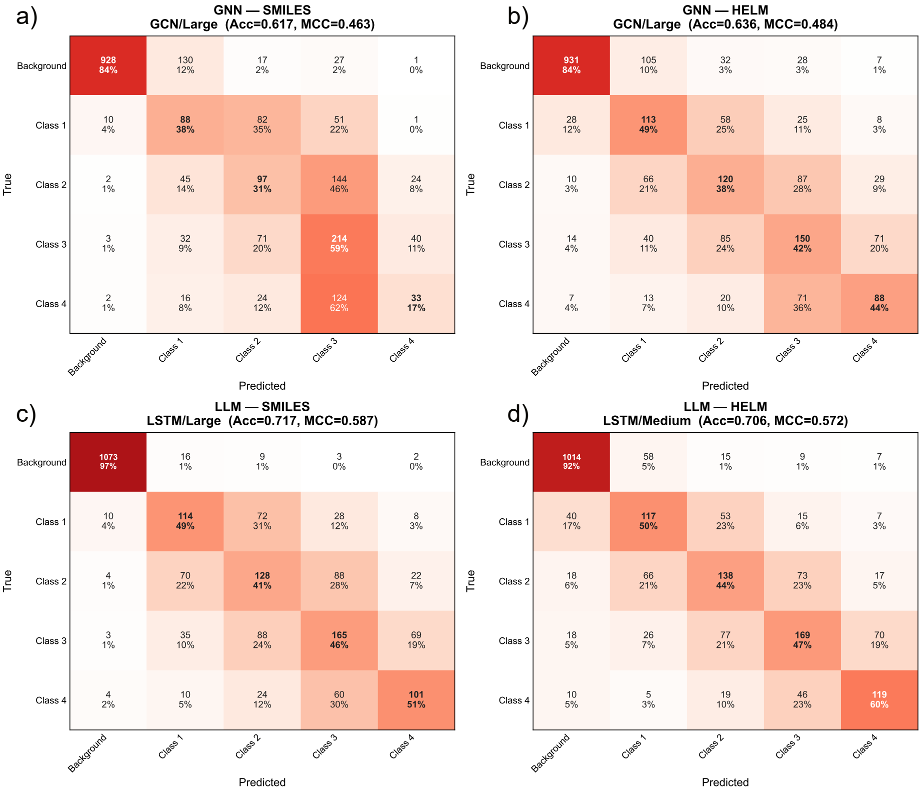

**Fig. S17: Recall-normalized confusion matrices for the four single-model quadrant winners.** $2 \times 2$ layout: top row, GNN winners ((a) SMILES: GCN-large; (b) HELM: GCN-large); bottom row, LLM winners ((c) SMILES: LSTM-large; (d) HELM: LSTM-medium). Rows of each matrix are reference classes and columns are predicted classes; each row sums to 1.0. Darker cells indicate larger fraction of the reference class predicted as the corresponding column class.

#### S5.3 HELM vs. SMILES Encoding Effect

The encoding axis does not have a consistent direction, but the pattern differs between the two families. Fig. S18 shows $\Delta\text{MCC} = \text{MCC(HELM)} - \text{MCC(SMILES)}$ for each (architecture, size) cell on a diverging red-blue scale. Within the LLM family, SMILES wins 7 of 9 (architecture, size) cells (HELM wins at GPT-medium and GRU-medium). The quadrant winners by validation MCC are LSTM-large SMILES (test 0.587) and LSTM-medium HELM (test 0.572), with a median cell-level test $\Delta$MCC of $-0.016$. Within the GNN family, the direction flips across architectures: GCN and Graph Transformer prefer HELM at all three sizes, whereas GAT and MPNN prefer SMILES in all sizes and GIN prefers SMILES in two of three; the quadrant winners by validation MCC split accordingly (GCN-large HELM, test 0.484 vs GCN-large SMILES, test 0.463, test $\Delta\text{MCC} = +0.021$). Across all 24 cells SMILES wins 15 and HELM wins 9. No cell shows a single-encoding dominance large enough to justify dropping the other. The ensemble of Section S5.4 therefore retains one member from each of the four (encoding, family) quadrants rather than committing to a single encoding.

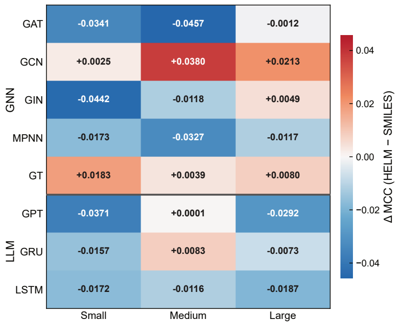

**Fig. S18: Encoding effect on test-set MCC across the 8-by-3 architecture-size grid.** Heatmap of ΔMCC = MCC(HELM) − MCC(SMILES). Red cells indicate HELM outperforming SMILES; blue cells indicate SMILES outperforming HELM. Rows are the 8 architectures (GNN family above, LLM family below, separated by a horizontal rule); columns are the three sizes (small, medium, large).

#### S5.4 Ensemble Strategy Comparison

We combine four single-model members into the ensemble, one per (encoding, family) quadrant: LSTM-large-SMILES (LLM-SMILES), LSTM-medium-HELM (LLM-HELM), GCN-large-HELM (GNN-HELM), and GCN-large-SMILES (GNN-SMILES). Selection is anchored on validation MCC rather than test MCC: test MCC must remain a held-out evaluation of the final ensemble, so test scores cannot steer member or weight choices. The four selected members reach val MCC 0.612, 0.602, 0.514, and 0.479 respectively, which double as the weights of the MCC-weighted ensemble below (normalized to sum to one).

Fig. S19a-c compares these four members against two ensemble aggregators: soft voting (equal weights) and MCC-weighted voting (weights proportional to each member's validation MCC, normalized to sum to one). The global gain from either ensemble is limited. Soft voting reaches MCC 0.618 (+0.031 over the best individual LSTM-large-SMILES at 0.587) and weighted voting reaches MCC 0.622 (+0.035), with the MCC-weighted variant adding only +0.004 over soft. On macro-F1 the pattern is similar (LSTM-large-SMILES 0.566 → soft 0.605 → weighted 0.609). The +0.035 MCC improvement over the best individual is supported by the bootstrap: the weighted-ensemble CI lower bound (0.598) lies above the LSTM-large-SMILES point estimate (0.587), placing the gain outside the best individual's bootstrap variability. We adopt MCC-weighted voting as the production aggregator because it at least matches soft voting on every reported metric. However, the 95% bootstrap CIs of the two strategies overlap heavily on the test set (soft: [0.594, 0.641]; weighted: [0.598, 0.644]), so this choice is principled rather than empirically decisive.

Per-class, the picture is less uniform and shows an explicit trade-off. Relative to LSTM-large-SMILES, the weighted ensemble drops background precision by 0.021 (0.981 → 0.960) while background recall is essentially unchanged (+0.001, from 0.973 to 0.974). The ensemble therefore becomes slightly less clean at the background boundary, not more aggressive in calling active. In exchange, the ensemble gains F1 on every active bin: class_1 +0.053 (0.478 → 0.531), class_2 +0.057 (0.404 → 0.461), class_3 +0.049 (0.469 → 0.518), and class_4 +0.065 (0.504 → 0.569). These active-bin gains do not come from a sharper background-versus-active boundary but from better resolution among the four active classes themselves: individual members confuse the four active classes broadly and in different ways (cf. Fig. S17), and the ensemble confusion matrices show a more balanced distribution across the four active bins (Fig. S19d-e). The net trade-off is acceptable because background F1 stays at 0.967 (down from 0.977) while the four active bins move from F1 0.40–0.50 to F1 0.46–0.57. The improvement is concentrated where the predictor matters for downstream filtering.

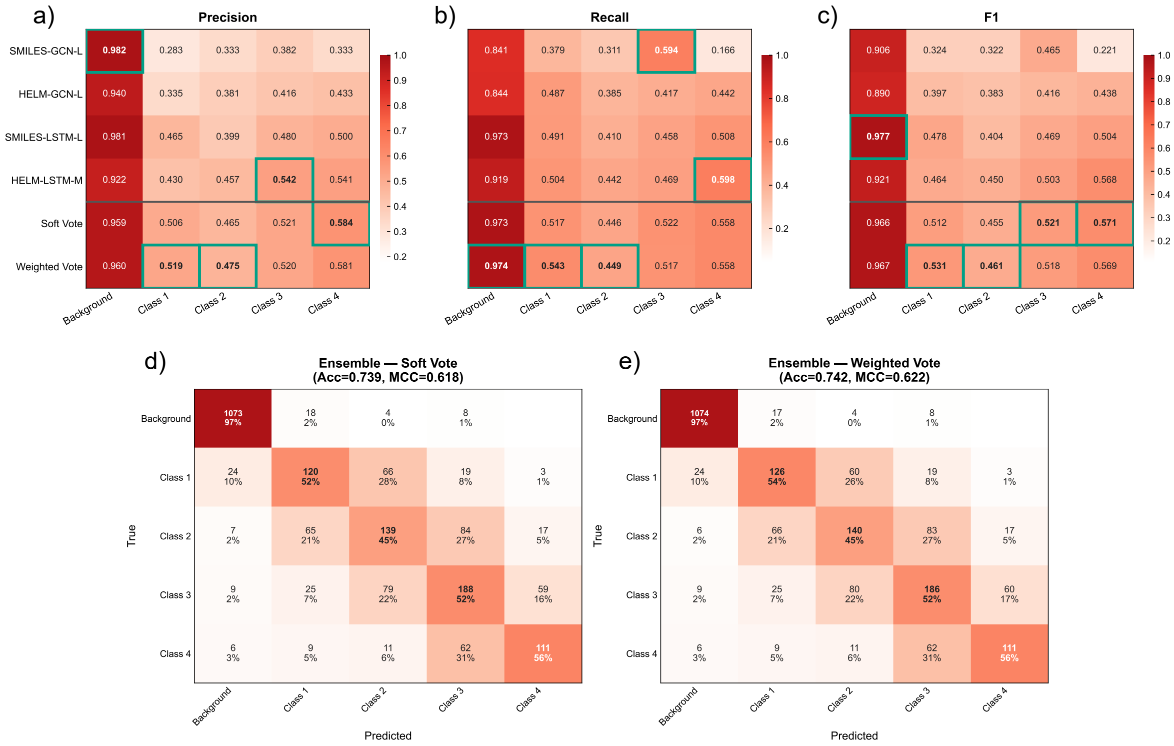

**Fig. S19: Per-class metrics and ensemble confusion matrices.** Five panels. Panels (a)–(c): per-class precision, recall, and F1 heatmaps. Within each panel, rows are the 6 entries (four individual members: LSTM-large-SMILES, LSTM-medium-HELM, GCN-large-HELM, GCN-large-SMILES; two ensemble methods: soft vote, MCC-weighted vote) and columns are the 5 classes (background, class_1, class_2, class_3, class_4). Per-column best cells are highlighted with a bold border. Row-label size suffix S/M/L = small/medium/large. Panels (d)–(e): recall-normalized confusion matrices for the soft-vote (d) and MCC-weighted (e) ensembles, with rows = reference classes, columns = predicted classes, and each row summing to 1.0.

The prediction head is treated as a downstream filter applied after the LCC cascade rather than as a standalone AMP classifier, and the search reported in this section is not exhaustive. We did not sweep alternative loss formulations beyond the focal plus ordinal-Gaussian combination of Table S8, nor alternative ensemble aggregators beyond soft and MCC-weighted voting, nor architectures outside the 8 covered in Table S9. The 5-class MCC plateau around 0.62 should therefore be read as the ceiling of this specific configuration rather than of the AMP prediction task in general. The background-precision-for-active-F1 trade-off of Section S5.4 is what justifies deploying the weighted ensemble in the generation pipeline's filter role, where the predictor's job is to rank generated candidates by predicted potency rather than to return a pure background call. An application that used the background prediction itself as a hard filter for downstream assays (accepting only peptides the model assigns to an active bin) would prefer the LSTM-large-SMILES single-model configuration, whose background predictions are very clean (precision 0.981) at the cost of the active-bin F1 improvements reported above. This PU setup has one weakness: the unlabelled pool can contain real but unreported AMPs, which are then trained as class 0. The model therefore sees some true actives labelled as background, so the active-class recall reported here is a conservative estimate. It should be used to rank and shortlist likely actives, not as a final non-AMP decision.

### S6 Inference Pipeline and Web Interface

Inference is exposed through a single entry point (`Pipelines/Inference.py`) that wraps the LCC cascade, an optional prediction layer, and a post-generation filter into one JSON-driven run. Figure S20 summarises the full flow and marks the points at which user-supplied constraints enter the pipeline.

#### S6.1 Inference Flow and Constraint Injection Points

The generator executes the three trained stages in sequence (*Layout* $\rightarrow$ *Content* $\rightarrow$ *Connection*) for each sampled candidate. Three input levels determine the entry point: Level 0 samples a HELM[1] peptide *de novo*, Level 1 supplies a user-specified layout token list (<PEP><LEN_7><CHEM><LEN_1>, . . . ) that skips the layout model, and Level 2 supplies a polymer-only HELM string so only the connection model is invoked. User constraints inject per stage: the layout model accepts `allowed_block_types`, global and per-type `length_range`, and `count_range` to bound block composition. The content model accepts sparse `{polymer:{position:monomer}}` maps that become AR prefix-forcing for autoregressive decoders (GPT, LSTM, GRU) or frozen-mask inputs for BERT. The connection model accepts a list of `forced_connections` whose R-group compatibility is checked against the monomer library before any generation work, together with a `forced_type_threshold` that rejects forced bonds whose model-assigned type probability falls below the cut-off.

Each generated HELM candidate then passes through an inline per-candidate check: HELM-to-SMILES-to-HELM round-trip validation (Section S1.1), `InChIKey` deduplication against the accumulated pool, and an optional `filter_connection_type` that retains only candidates containing every listed special-connection type. Accepted candidates stream to disk via `candidates.csv` with a companion `raw.jsonl` recording rejected drafts for diagnostics. Prediction is opt-in: the `prediction.task` field in the JSON attaches one or more scorers to the running pool, spanning the internal AMP ensemble of Section S5, the external PeptiVerse[30] ADMET predictor, and arbitrary user-registered plugins that follow the `predictor_config.json` contract. A final pool-level filter (`postprocess.prediction_filters`, `ranking`, `max_candidates`) selects the shortlist written to `filtered.csv`. This section carries the active-peptide thresholds (`accept_classes`), the safety constraints (`min_extras` keyed on `hemolysis_confidence` and `toxicity_confidence`), and the sort keys that drive the ranking.

Registration of a new predictor follows a lightweight two-file contract. A `predictor_config.json` declares the path to an adapter module, the conda environment in which the adapter is to be invoked, and the set of properties the predictor returns with their types (classification or regression). An accompanying `adapter.py` exposes two entry points: `init(config)` loads the underlying model once at startup, and `predict(model, smiles, properties)` returns a per-candidate extras dictionary whose keys follow a fixed naming convention (`{prop}_label` and `{prop}_confidence` for classification, `{prop}_{unit}` for regression) so that any returned field can be referenced directly in `postprocess.prediction_filters` and `ranking`. Each plugin runs in a dedicated subprocess bound to its declared conda environment, isolating dependency stacks (e.g., conflicting PyTorch or RDKit versions) between PepForge and third-party tools. A working template (`Configs/Inference/External_predictor_template/`) and an end-to-end tutorial (`Docs/Tutorials/custom_predictor.md`) are distributed with the repository.

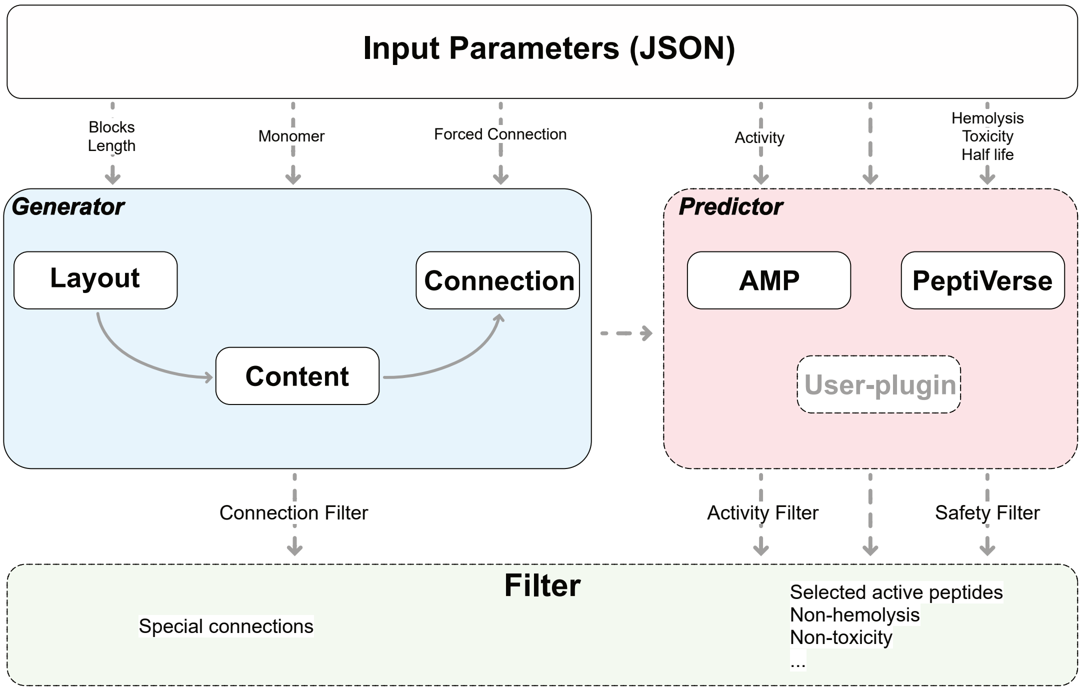

**Fig. S20: Inference pipeline and constraint injection points.** Top: the JSON configuration fans out into the pipeline, with per-stage fields driving the generator (block layout, monomer content, forced connections) and per-task fields driving the predictor and filter. Generator: the three trained stages (*Layout*, *Content*, *Connection*) run as a fixed LCC cascade. The dashed arrow to the predictor indicates that prediction is opt-in. Predictor: the internal AMP ensemble, the external PeptiVerse ADMET scorer, and user-registered plugins can be attached in parallel. Filter: a single post-processing stage that applies connection-type constraints from the generator and potency or safety thresholds from the predictor to the candidate pool.

#### S6.2 JSON Input Schema

A single JSON object controls the entire run. The seven top-level sections group options by role: (1) `enable_generation` and `input_csv` choose between full generation and predict-only mode, (2) `input` sets the LCC cascade entry level, (3) `constraints` holds the three per-stage injection slots of Section S6.1, (4) `params` collects sampling hyperparameters (`num_samples`, `temperature`, `top_k`, `validation_workers`, `seed`), (5) `models` overrides default checkpoint paths, (6) `prediction` selects tasks and per-task options, and (7) `postprocess` carries the filter, ranking, and truncation rules. Listing S1 shows a typical request that generates 1,000 disulfide-bridged candidates with `Cys` fixed at positions 1 and 7, scores them with the AMP ensemble and PeptiVerse, and retains the top-ranked active, non-hemolytic, non-toxic peptides. A complete, annotated template with every field documented in place is distributed as `Configs/Inference/inference_request_template.json`, and end-to-end usage examples together with the corresponding web-form mappings are collected in the online tutorial linked from the project README.

**Listing S1:** Example JSON inference request covering all seven top-level sections.

```
{
  "name": "disulfide_amp_screen",
  "enable_generation": true,
  "input": { "helm": null, "layout_tokens": null },          // Level 0 (de novo)
  "constraints": {
    "layout_constraints":  { "PEP_length_range": [7, 12] },
    "content_constraints": { "PEPTIDE1": { "1": "C", "7": "C" } },
    "forced_connections":  [ { "from_polymer": "PEPTIDE1", "from_pos": 1,
                               "to_polymer":   "PEPTIDE1", "to_pos":   7,
                               "connection_type": "Disulfide" } ],
    "forced_type_threshold": 0.5
  },
  "params": { "num_samples": 1000, "temperature": 1.0, "seed": 42 },
  "prediction": { "task": ["amp", "peptiverse"] },
  "postprocess": {
    "filter_connection_type": ["Disulfide"],
    "prediction_filters": {
      "amp":        { "accept_classes": ["class_3", "class_4"], "min_confidence": 0.6 },
      "peptiverse": { "accept_extras":  { "hemolysis_label": ["non_hemolytic"],
                                          "toxicity_label":  ["non_toxic"] } }
    },
    "ranking":        { "sort_by": ["amp.predicted_class", "amp.confidence"] },
    "max_candidates": 100
  }
}
```

#### S6.3 Web Interface

A local web interface wraps the same JSON schema in an interactive form so that users without a scripting background can drive generation, prediction, and post-filtering from the browser. The application is distributed as part of the repository (FastAPI backend, React/Vite frontend, launched by `App/run.sh`) and runs entirely on the user's own machine with no cloud deployment. Figure S21 shows the two views that together cover the supported workflows.

The Generate+Predict panel (Fig. S21a) maps every field of the JSON schema to a form widget. Numbered callouts in the left sidebar mark (1) the LCC cascade entry-mode selector (`De Novo`, `From Layout`, `From Content`, corresponding to the three values of `input` in the JSON schema), (2) the sampling-parameter controls (`num_samples`, `temperature`), (3) the three per-stage constraint toggles (*Layout*, *Content*, *Connection*), all off in this Level 0 *de novo* example, and (4) the optional built-in predictor section, collapsed in the main view. The left inset (5) shows this section expanded, exposing the AMP Activity scorer (the four-model MCC-weighted ensemble of Section S5) and the PeptiVerse ADMET scorer as independent toggles with per-scorer post-filter controls stacked below. Callout (6) marks the expanded output card for a deliberately borderline candidate: the generated tripeptide `PEPTIDE1{[G].[Mnm_22].[G]}$$$$` is assigned AMP class 4 with only 25% peak confidence (flat class distribution 21%/19%/17%/18%/25% over `background` and `class_1`–`class_4`), PeptiVerse reports a 1.0 h half-life, and both hemolysis and toxicity flags read `Risk`, so a downstream `min_confidence` $>$ 0.25 rule would reject this row even though its argmax class lies in the active range.

The Monomer Library view (Fig. S21b), reached from the Library tab, exposes the 425-monomer catalogue that backs the `content_constraints` field. Callouts mark (1) the three search-mode tabs (`Browse`, `Search by Name`, `Search by SMILES`), (2) the per-monomer detail card (here `*A`, Alanine) listing the HELM symbol, natural analog, polymer class, canonical SMILES, `InChIKey`, and R-group specifications, (3) the scrollable catalogue with filter tabs for `All`, `Backbone`, `Terminal`, and `Link`, and (4) an inline structure preview that tracks the currently highlighted row (showing `1Nal`, 3-(1-naphthyl)-alanine in this screenshot), letting users scan structures without opening each card individually. Generated candidates stream back into an inline results table mirroring `candidates.csv` and `filtered.csv`, with per-candidate cards expanding to show the full prediction breakdown.

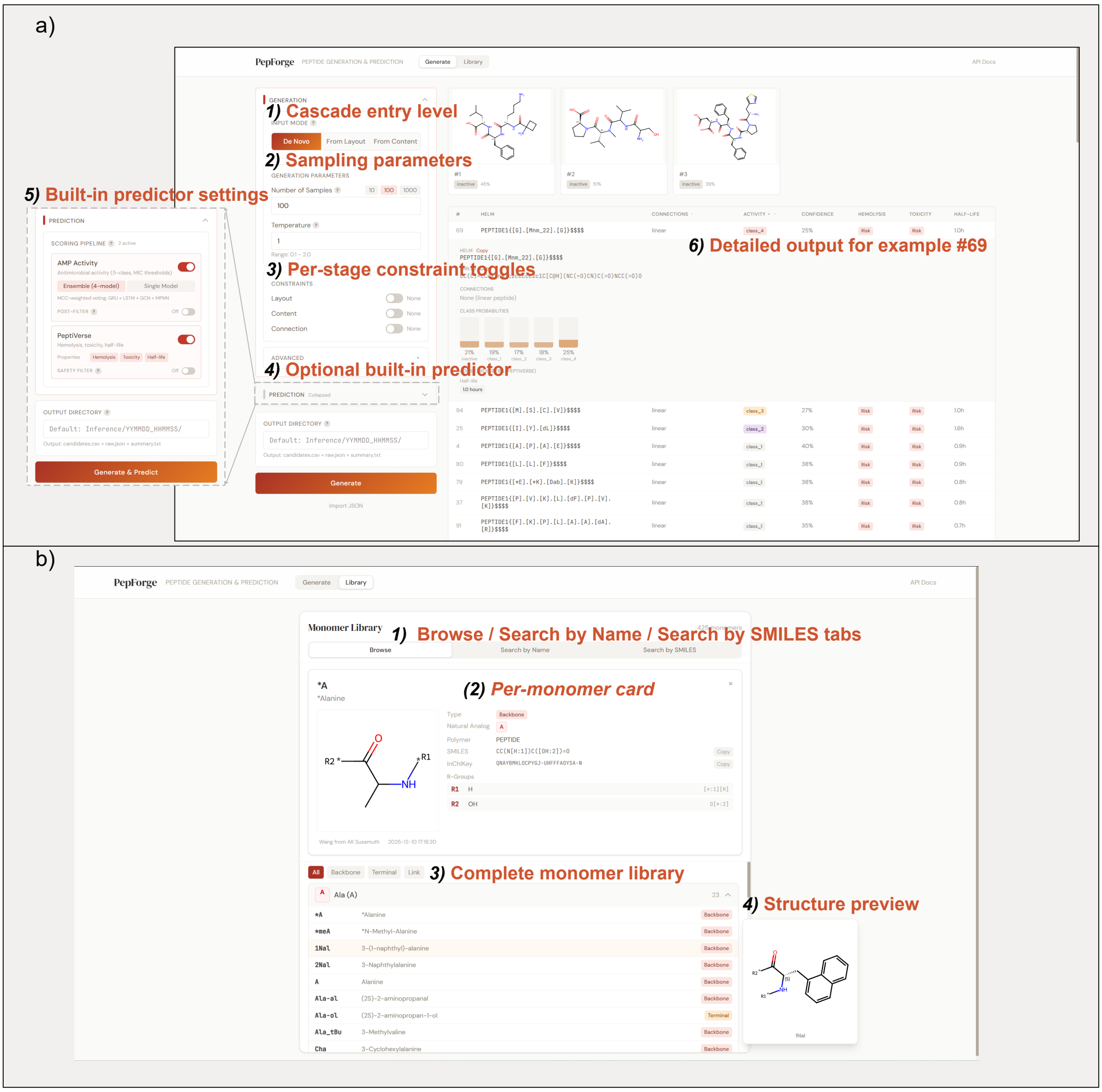

**Fig. S21: Local web interface for inference and monomer lookup. (a)** Generate+Predict panel. **(b)** Monomer Library view. Numbered callouts reference the walkthroughs in Section S6.3.

### S7 Reference Configurations and Reproducibility

This section collects reference configurations that support reproducibility of experiments reported in the main text but do not belong to any per-module evaluation section. Per-module architecture and training hyperparameters are embedded in their respective sections: *Layout* in Section S3.1 (Table S4), *Content* in Section S3.2 (Table S5), *Connection* in Section S3.3 (Table S6), and the prediction architectures in Section S5.1 (Table S9).

#### S7.1 Flat GPT Baseline Configuration

The Flat GPT baseline evaluated against the LCC cascade in Fig. 3a–f is a single autoregressive GPT[16] decoder trained on the same 383,817-peptide training split used by every module of the LCC cascade, but exposed to HELM[1] as a flat token stream over a full-HELM vocabulary of 510 tokens (block delimiters, monomer symbols, R-group labels, connection notation, and special tokens). Architecturally it mirrors the Content GPT-medium backbone, providing a representative end-to-end baseline trained on the same HELM corpus as the LCC cascade. The LCC cascade's *Content* stage uses a larger GPT-Large backbone (d=768, L=12), so the comparison varies decoding structure (factorized vs flat) and backbone capacity together rather than holding capacity fixed. The implementation lives under `Scripts/Flat_Generation/` and reuses the same `BaseTrainer` infrastructure (cosine learning-rate schedule, EMA with decay 0.99, 20-epoch early-stopping patience on validation loss, max-gradient-norm clipping at 1.0) that underlies every module in Section S3. Table S10 summarises architecture and module-specific training settings alongside the Content GPT-medium reference row.

**Table S10: Flat GPT architecture and training hyperparameters, with Content GPT-medium as a reference.**

| | Architecture | | | | | | | Training[d] | | | | | | | |
|---|---|---|---|---|---|---|---|---|---|---|---|---|---|---|---|
| **Model** | vocab | `d_model` | heads | layers | `dim_ff` | dropout | `max_len` | batch | epochs | lr | wd | ls | EMA | pat | clip |
| Flat GPT | 510[b] | 512 | 8 | 8 | 1024 | 0.35 | 512 | 256 | 200 | $1\times10^{-4}$ | 0.05 | 0.1 | 0.99 | 20 | 1.0 |
| Content GPT-medium[a] | 498[c] | 512 | 8 | 8 | 1024 | 0.35 | 512 | 384 | 200 | $1\times10^{-4}$ | 0.05 | 0.25 | 0.99 | 20 | 1.0 |

[a] Full per-architecture *Content*-model table is Table S5 in Section S3.2.

[b] Full-HELM tokenisation: block delimiters, monomer symbols, R-group labels, connection notation, and special tokens.

[c] *Content* stage sees only the monomer-symbol subset plus special tokens. The *Layout* stage operates on its own 70-token vocabulary of block types and length tokens (Section S3.1).

[d] "wd" is weight decay, "ls" is uniform label smoothing, "pat" is early-stopping patience (epochs), and "clip" is the max-gradient-norm threshold.

The exported checkpoint, exposed to downstream inference as `best.pt`, drives every Flat GPT sampling run reported in the main text. Its path is given in Listing S3.

#### S7.2 Free Generation at 100K (Fig. 3a–f)

The 100K comparison between the LCC cascade and Flat GPT was executed as two SLURM arrays of five independent runs with shared seeds $\{42, 123, 456, 789, 1024\}$, under `jobs/Inference/Cascade_vs_Flat/`. Each run targets 100,000 raw samples and every candidate passes the HELM-to-SMILES round-trip gate. Peptides failing the gate are dropped before validity, uniqueness, and novelty are computed. Evaluation of both models is performed by the same script `Scripts/Inference/`

`evaluate_generation.py`, taking the raw JSONL output and the training CSV `Data/all_peptides/HELM.csv` as reference.

The LCC cascade arm is driven by the full pipeline `Pipelines/Inference.py` with a per-run JSON request. Listing S2 shows the request for run 1 (seed 42). The other four runs differ only in the `seed` field and the per-run `output_dir`. Sampling is temperature 1.0 with top-$k = 64$ and a per-sample cap of 8 blocks and 25 monomers, identical to the production run of Section S7.5. Checkpoints are the best per-stage models (Layout GPT, Content GPT-Large, Connection GAT-Large) selected in Sections S3.1–S3.3.

**Listing S2:** LCC cascade 100K free-generation request (run 1 of 5, seed 42).

```
{
  "name": "cascade_run1",
  "enable_generation": true,
  "input": { "helm": null, "layout_tokens": null },
  "constraints": {
    "layout_constraints":    null,
    "content_constraints":   null,
    "forced_connections":    null,
    "forced_type_threshold": 0.5
  },
  "params": {
    "num_samples":             100000,
    "temperature":             1.0,
    "top_k":                   64,
    "max_blocks":              8,
    "max_monomers_per_block": 25,
    "validation_workers":      8,
    "no_roundtrip":            false,
    "save_raw":                true,
    "seed":                    42,       // runs 2..5 -> 123, 456, 789, 1024
    "output_dir":              "Paper/Figures/Data/.../Cascade/run1"
  },
  "models": {
    "layout_ckpt":      "Models/Generation/Layout/260210_GPT.pt",
    "content_ckpt":     "Models/Generation/Content/GPT_L_260226.pt",
    "connection_ckpt": "Models/Generation/Connection/GAT_L_260226.pt"
  },
  "postprocess": null
}
```

The Flat GPT arm calls a dedicated CLI script `infer_flat_gpt.py` that bypasses the LCC cascade and decodes the flat-HELM vocabulary end to end. Listing S3 shows the invocation for run 1; the other four runs differ only in the `-seed` value and the `-output_dir`. Decoding defaults to nucleus sampling (`top_p` = 0.9) with `top_k` disabled, which is the Flat-GPT inference default rather than the LCC cascade's `top_k` = 64. Temperature stays at 1.0 and `max_len` at 256 tokens. A matched-decoding ablation (identical `top_k` or `top_p` across the two arms) is left to future work, and the reported novelty ordering should be read with this decoding-policy caveat in mind.

**Listing S3:** Flat GPT 100K free-generation CLI (run 1 of 5, seed 42). Runs 2–5 differ only in the -seed value.

```
python Scripts/Flat_Generation/Inference/infer_flat_gpt.py \
    --ckpt Scripts/Flat_Generation/Models_ckpt/260225_flat_gpt/best.pt \
    --num_samples 100000 \
    --batch_size 512 \
    --validation_workers 8 \
    --seed 42 \
    --output_dir Paper/Figures/Data/.../Flat/run1
# Internal defaults: temperature=1.0, top_p=0.9, top_k disabled, max_len=256
```

#### S7.3 *Content* Infilling (Fig. 3g–i)

The GPT-vs-BERT infilling comparison operates on a frozen evaluation subset of 10,000 peptides drawn from the held-out test split `test.jsonl` with a fixed random seed of 42, and exposes monomer positions at one of three reveal ratios in $\{0.3, 0.5, 0.7\}$. Constraints are materialised once per ratio by `prepare_constraints.py` and re-read by every run so that the mask pattern is identical across architectures. Each ratio is sampled five times with seeds $\{43, 44, 45, 46, 47\}$, giving a SLURM array of $3 \times 5 = 15$ jobs. Both architectures share the *Connection* stage (GAT-Large, identical checkpoint) and differ only in the Content checkpoint, GPT-Large or BERT-Large. Listing S4 shows the invocation for one array task. Full paths are under `jobs/Inference/GPT_vs_BERT_Infilling/`.

**Listing S4:** Infilling comparison CLI (array task 1 of 15: ratio 0.3, run 1).

```
python Paper/Figures/Generation_Model/End2end/GPT_vs_BERT_Infilling/run_comparison.py \
    --constraints Paper/Figures/Data/.../GPT_vs_BERT_Infilling/constraints/ratio_0.3.jsonl
    \
    --output_dir  Paper/Figures/Data/.../GPT_vs_BERT_Infilling/ratio_0.3 \
    --run_id      1 \
    --seed        43 \
    --temperature 1.0 \
    --gpt_ckpt        Models/Generation/Content/GPT_L_260226.pt \
    --bert_ckpt       Models/Generation/Content/BERT_L_260301.pt \
    --connection_ckpt Models/Generation/Connection/GAT_L_260226.pt \
    --training_csv    Data/all_peptides/HELM.csv
# Array task i -> ratio_idx = (i-1)/5, run_id = (i-1)%5 + 1, seed = 42 + run_id
```

#### S7.4 Constrained Generation (Table S7)

The nine constrained-generation scenarios of Section S4 are executed as a SLURM array of nine jobs (`submit_cascade_constraint.sh` under `jobs/Inference/Cascade/Constraint_G/`). Every scenario reuses the best-of-stage LCC cascade checkpoints (Content GPT-Large, Connection GAT-Large; Layout is skipped in scenarios that ship a `layout_tokens` prefix) and the common sampling parameters shared across all nine requests, but reduces the target pool to 1,000 accepted peptides with a raw-sample budget of 100,000 attempts (Section S4). A single seed of 42 is used per scenario, since the nine experiments are qualitative sweeps over constraint-specification strategies rather than statistical averages. Listing S5 records the shared `params`/`models` block, and Listings S6–S7 give the per-scenario `input` and `constraints` fields for CG1 through CG9.

**Listing S5:** Shared `params` and `models` block, identical across CG1–CG9.

```
"params": {
  "num_samples":            1000,
  "temperature":            1.0,
  "top_k":                  64,
  "max_blocks":             8,
  "max_monomers_per_block": 25,
  "no_roundtrip":           false,
  "save_raw":               true,
  "seed":                   42,
  "output_dir":             "Paper/Figures/Data/.../Constrained_Generation/<CGk>"
},
"models": {
  "layout_ckpt":     null,    // Layout skipped when "layout_tokens" is provided
  "content_ckpt":    "Models/Generation/Content/GPT_L_260226.pt",
  "connection_ckpt": "Models/Generation/Connection/GAT_L_260226.pt"
}
```

**Listing S6:** Per-scenario `input` and `constraints` fields, CG1–CG5. Every request also carries the shared `params`/`models` block of Listing S5 and the fixed `"forced_type_threshold": 0.5`.

```
// --- CG1_L: layout-only, PEP x 7 ---
"input":       { "layout_tokens": ["<PEP>","<LEN_7>"] },
"constraints": { }

// --- CG2_LC_Cys: layout + content, Cys fixed at positions 3 and 8 ---
"input":       { "layout_tokens": ["<PEP>","<LEN_10>"] },
"constraints": {
  "content_constraints": { "PEPTIDE1": { "3": "C", "8": "C" } }
}

// --- CG3_LCC_SS: layout + content + forced Disulfide(2,8) ---
"input":       { "layout_tokens": ["<PEP>","<LEN_9>"] },
"constraints": {
  "content_constraints": { "PEPTIDE1": { "2": "C", "8": "C" } },
  "forced_connections":  [ { "from_polymer": "PEPTIDE1", "from_pos": 2,
                             "to_polymer":   "PEPTIDE1", "to_pos":   8,
                             "connection_type": "Disulfide" } ]
}

// --- CG4_LCC_Lanthi: layout + content + rare Lanthionin(1,5) ---
"input":       { "layout_tokens": ["<PEP>","<LEN_7>"] },
"constraints": {
  "content_constraints": { "PEPTIDE1": { "1": "C", "5": "dA" } },
  "forced_connections":  [ { "from_polymer": "PEPTIDE1", "from_pos": 1,
                             "to_polymer":   "PEPTIDE1", "to_pos":   5,
                             "connection_type": "Lanthionin" } ]
}

// --- CG5_L_multi: layout-only, multi-block PEP + CHEM ---
"input":       { "layout_tokens": ["<PEP>","<LEN_9>","<CHEM>","<LEN_1>"] },
"constraints": { }
```

**Listing S7:** Per-scenario `input` and `constraints` fields, CG6–CG9 (continuation of Listing S6).

```
// --- CG6_LCC_multi_Sulfa: layout + forced Sulfanilamide(PEP:5, CHEM:1), content free ---
"input":       { "layout_tokens": ["<PEP>","<LEN_10>","<CHEM>","<LEN_1>"] },
"constraints": {
  "forced_connections": [ { "from_polymer": "PEPTIDE1", "from_pos": 5,
                            "to_polymer":   "CHEM1",    "to_pos":   1,
                            "connection_type": "Sulfanilamide" } ]
}

// --- CG7_L_filter_SS: layout + post-hoc filter on rare types ---
"input":       { "layout_tokens": ["<PEP>","<LEN_12>"] },
"constraints": {
  "filter_connection_type": ["Trp-Cys", "head2tail ring"]
}

// --- CG8_LayoutRange: layout range, free block types ---
"input":       { "layout_tokens": null },
"constraints": {
  "layout_constraints": { "PEP_length_range": [5, 10] }
}

// --- CG9_PEPonly: layout range, PEP-only ---
"input":       { "layout_tokens": null },
"constraints": {
  "layout_constraints": { "allowed_block_types": ["PEP"],
                          "PEP_length_range":    [5, 12] }
}
```

#### S7.5 10M-Scale Generation for Candidate Discovery (Fig. 4)

The production-scale run behind Fig. 4 candidate discovery was executed as a SLURM array of five independent shards (`jobs/Inference/G+AMP_10M/`), each generating 2,000,000 *de novo* HELM peptides with the same constraints-free configuration but a distinct random seed (shard $k$ uses `seed` $= k$ for $k \in \{1, 2, 3, 4, 5\}$), yielding 10M raw candidates in total. Sampling reuses the same parameters as the 100K LCC cascade run in Section S7.2 (temperature 1.0, top-$k =$ 64, per-sample cap of 8 blocks and 25 monomers, `forced_type_threshold` $=$ 0.5), with `validation_workers` raised to 12 to keep the round-trip gate CPU-bound rather than GPU-bound. Each shard runs on a single GPU with compute capability $\geq$ 8.6 (Ampere or Ada Lovelace generation, e.g., RTX A6000 or RTX 6000 Ada) under a wall-clock budget of four days. Prediction is invoked in the same job so AMP and PeptiVerse scores are emitted alongside the raw candidates, but no `postprocess` filter is applied at this stage. A separate merge step (`jobs/Inference/G+AMP_10M/submit_merge.sh`) unions the five per-shard `candidates.csv` files, deduplicates by `InChIKey`, subtracts training-set structures, and produces the 4.78M-peptide novel pool reported in the main text. Listing S8 shows the JSON for shard 0. The remaining four shard configs differ only in the `seed` field and the per-shard `output_dir`.

**Listing S8:** Inference request for shard 0 of the 10M production run. Shards 0–4 differ only in the seed (1–5) and output directory.

```
{
  "name": "G+AMP_10M_shard0",
  "enable_generation": true,
  "input": { "helm": null, "layout_tokens": null },
  "constraints": {
    "layout_constraints":     null,
    "content_constraints":    null,
    "forced_connections":     null,
    "forced_type_threshold":  0.5
  },
  "params": {
    "num_samples":            2000000,   // 2M per shard x 5 shards = 10M
    "temperature":            1.0,
    "top_k":                  64,
    "max_blocks":             8,
    "max_monomers_per_block": 25,
    "validation_workers":     12,
    "seed":                   1,         // shard 0..4 -> seed 1..5
    "output_dir":             "Inference/260307_G+AMP_10M/shard0"
  },
  "models": {
    "layout_ckpt":     "Models/Generation/Layout/260210_GPT.pt",
    "content_ckpt":    "Models/Generation/Content/GPT_L_260226.pt",
    "connection_ckpt": "Models/Generation/Connection/GAT_L_260226.pt"
  },
  "prediction": {
    "task":       ["amp", "peptiverse"],
    "device":     "cuda",
    "batch_size": 64
  },
  "postprocess": null                    // filtering applied post-merge
}
```

#### S7.6 Reproducing Training from Scratch

Retraining is typically motivated by one of three library-level changes: adding new monomers, adding new special-connection types, or replacing the training set with an updated HELM CSV. Monomer extension is handled by the one-click utility `Pipelines/Add_Monomer.py` described in Section S2.1, which deduplicates a user-supplied CSV against the existing `HELMLibrary.json` by `InChIKey`, extends the ChemBERTa embedding tensor using the `raw_std` recorded in the embedding manifest so that earlier checkpoints stay reproducible (Section S2.3), and assigns continuous identifiers to each accepted entry. A new special-connection type is added by appending a `SPECIAL_BOND_INFO` entry to `Configs/Special_connection/special_connections.py` with four fields: a short `Name`, a `SMARTS` pattern identifying the bond, the pair of SMARTS atom indices marking the bond-break positions (`start_atom_pos`, `end_atom_pos`), and an optional `add_atom` record for an atom restored during `HELM2SMILES` reconstruction (for example, the carbonyl oxygen reintroduced on amide cleavage). The roundtrip pipeline of Section S1.1 loads every entry at import time, so the new type becomes available to both the forward decomposer and the reverse reconstructor without further wiring. After any of these three changes, a single call to `Scripts/Generation_Model/Scripts/prepare_data.py` (with `--input_csv` pointing to the HELM source of choice) regenerates the hierarchical splits, the tokeniser vocabularies, and the auto-loaded connection-type list consumed by the *Connection* stage, so that the downstream training wrappers see the updated chemical space without further manual configuration.

Environment provisioning uses `install.py`, documented in the top-level `README.md`. Training is launched through the same three wrappers used throughout this SI: `Pipelines/Train_Generation.py` for the LCC cascade (Section S3), `Pipelines/Train_Prediction.py` for the prediction ensemble (Section S5, invoked once per encoding mode), and `train_flat_gpt.py` under `Scripts/Flat_Generation/` for the Flat GPT baseline of Section S7.1. Listing S9 shows the library-update steps (A)–(C) and the minimal retraining invocations (D), and full per-architecture flag lists are in `README.md`. Promoting the retrained checkpoints to the production LCC cascade of Section S7.5 follows the convention recorded in `Models/Generation/MODEL_REGISTRY.md`.

Compute expectations follow a predictable pattern on the reference hardware (NVIDIA RTX 6000 Ada, 48 GB) used throughout this SI. The *Layout* stage trains end-to-end in under one GPU-hour per architecture because the vocabulary is small and convergence is reached within 25 epochs (Section S3.1). Content-large and BERT-large each take 10–14 GPU-hours on the full 383,817-peptide split, dominated by the autoregressive pass over the 498-monomer vocabulary at batch size 128 (Section S3.2). Connection GAT-large completes in 6–8 GPU-hours, limited by R-group-compatible edge enumeration rather than by GNN forward cost (Section S3.3). Each AMP prediction ensemble member trains in 2–4 GPU-hours because the tagged pool is two orders of magnitude smaller than the generation pool (Section S5.1). End-to-end retraining of the full production LCC cascade plus the four ensemble members therefore fits within one overnight run on a single GPU, and the per-stage checkpoints can be swapped into the production pipeline of Section S7.5 without retraining the stages that were not affected by the library change. Users whose changes are confined to a single stage (for example, extending only the monomer library without adding a new special-connection type) should retrain only that stage, because the three stages consume disjoint supervision signals and a retrained Content or Connection checkpoint can be combined with an unchanged Layout checkpoint at inference time without loss of validity.

**Listing S9:** Retraining workflow after a chemistry extension. (A)–(B) update the monomer library or connection-type registry; (C) regenerates splits and tokenizers (always required after (A) or (B)); (D) retrains the production LCC cascade and prediction ensemble. See `README.md` for full per-architecture flag lists.

```
# (A) Add new monomers: populate Configs/Monomer/Add_Monomer.csv
# (symbol, SMILES, type, natural analog, R-group caps), then:
python Pipelines/Add_Monomer.py

# (B) Add a new connection type: append a dict entry to
# SPECIAL_BOND_INFO in Configs/Special_connection/special_connections.py,
# following the schema of existing entries (Name, SMARTS,
# start_atom_pos, end_atom_pos, add_atom).

# (C) Regenerate splits, tokenizers, connection-type list
python Scripts/Generation_Model/Scripts/prepare_data.py \
    --input_csv  Data/all_peptides/HELM.csv \
    --output_dir Scripts/Generation_Model/Data/processed

# (D) Retrain. Production default: Layout GPT + Content GPT-large
# + Connection GAT-large. Expand *_arch_list / *_size_list to sweep.
python Pipelines/Train_Generation.py \
    --layout_arch_list gpt \
    --content_arch_list gpt --content_size_list large \
    --connection_arch_list gat --connection_size_list large

# Prediction ensemble (4 members, one per encoding x family quadrant):
# LSTM-medium-HELM, GCN-large-HELM, LSTM-large-SMILES, GCN-large-SMILES.
# Train the two HELM members in one call (LSTM medium, GCN large):
python Pipelines/Train_Prediction.py \
    --llm_arch_list lstm --llm_size_list medium --llm_use_smiles False \
    --gnn_arch_list gcn  --gnn_size_list large
# Train the two SMILES members in one call (LSTM large, GCN large):
python Pipelines/Train_Prediction.py \
    --llm_arch_list lstm --llm_size_list large  --llm_use_smiles True \
    --gnn_arch_list gcn  --gnn_size_list large  --gnn_use_smiles True
```

[12] Seyone Chithrananda, Gabriel Grand, and Bharath Ramsundar. ChemBERTa: large-scale self-supervised pretraining for molecular property prediction. *arXiv preprint arXiv:2010.09885*, 2020. Preprint.

[13] Thomas Wolf, Lysandre Debut, Victor Sanh, Julien Chaumond, Clement Delangue, Anthony Moi, Pierric Cistac, Tim Rault, Remi Louf, Morgan Funtowicz, Joe Davison, Sam Shleifer, Patrick von Platen, Clara Ma, Yacine Jernite, Julien Plu, Canwen Xu, Teven Le Scao, Sylvain Gugger, Mariama Drame, Quentin Lhoest, and Alexander Rush. Transformers: State-of-the-art natural language processing. In *Proceedings of the 2020 Conference on Empirical Methods in Natural Language Processing: System Demonstrations*, pages 38–45. Association for Computational Linguistics, 2020. doi: 10.18653/v1/2020.emnlp-demos.6.

[14] Laurens van der Maaten and Geoffrey Hinton. Visualizing data using t-SNE. *Journal of Machine Learning Research*, 9(86):2579–2605, 2008.

[15] Adam Paszke, Sam Gross, Francisco Massa, Adam Lerner, James Bradbury, Gregory Chanan, Trevor Killeen, Zeming Lin, Natalia Gimelshein, Luca Antiga, et al. PyTorch: An imperative style, high-performance deep learning library. In *Advances in Neural Information Processing Systems*, volume 32, 2019.

[16] Ashish Vaswani, Noam Shazeer, Niki Parmar, Jakob Uszkoreit, Llion Jones, Aidan N. Gomez, Łukasz Kaiser, and Illia Polosukhin. Attention is all you need. In *Advances in Neural Information Processing Systems*, volume 30, 2017.

[17] Kyunghyun Cho, Bart van Merriënboer, Caglar Gulcehre, Dzmitry Bahdanau, Fethi Bougares, Holger Schwenk, and Yoshua Bengio. Learning phrase representations using RNN encoder–decoder for statistical machine translation. In *Proceedings of the 2014 Conference on Empirical Methods in Natural Language Processing (EMNLP)*, pages 1724–1734. Association for Computational Linguistics, 2014. doi: 10.3115/v1/D14-1179.

[18] Sepp Hochreiter and Jürgen Schmidhuber. Long short-term memory. *Neural Computation*, 9(8): 1735–1780, 1997. doi: 10.1162/neco.1997.9.8.1735.

[19] Jacob Devlin, Ming-Wei Chang, Kenton Lee, and Kristina Toutanova. BERT: Pre-training of deep bidirectional transformers for language understanding. In *Proceedings of the 2019 Conference of the North American Chapter of the Association for Computational Linguistics: Human Language Technologies (NAACL-HLT)*, pages 4171–4186. Association for Computational Linguistics, 2019. doi: 10.18653/v1/N19-1423.

[20] Petar Veličković, Guillem Cucurull, Arantxa Casanova, Adriana Romero, Pietro Liò, and Yoshua Bengio. Graph attention networks. In *International Conference on Learning Representations*, 2018.

[21] Thomas N Kipf and Max Welling. Semi-supervised classification with graph convolutional networks. In *International Conference on Learning Representations*, 2017.

[22] Keyulu Xu, Weihua Hu, Jure Leskovec, and Stefanie Jegelka. How powerful are graph neural networks? In *International Conference on Learning Representations*, 2019.

[29] Weihua Hu, Matthias Fey, Marinka Zitnik, Yuxiao Dong, Hongyu Ren, Bowen Liu, Michele Catasta, and Jure Leskovec. Open graph benchmark: Datasets for machine learning on graphs. In *Advances in Neural Information Processing Systems*, volume 33, 2020.

[30] Yinuo Zhang, Sophia Tang, Tong Chen, Elizabeth Mahood, Sophia Vincoff, and Pranam Chatterjee. PeptiVerse: A unified platform for therapeutic peptide property prediction. *bioRxiv*, 2026. doi: 10.64898/2025.12.31.697180. Preprint.
